# Sequencing 1206 genomes reveals origin and movement of *Aedes aegypti* driving increased dengue risk

**DOI:** 10.1101/2024.07.23.604830

**Authors:** Jacob E. Crawford, Dario Balcazar, Seth Redmond, Noah H. Rose, Henry A. Youd, Eric R. Lucas, Rusdiyah Sudirman Made Ali, Ashwaq Alnazawi, Athanase Badolo, Chun-Hong Chen, Luciano V. Cosme, Jennifer A. Henke, Kim Y. Hung, Susanne Kluh, Wei-Liang Liu, Kevin Maringer, María Victoria Micieli, Evlyn Pless, Aboubacar Sombié, Sinnathamby N. Surendran, Isra Wahid, Peter A. Armbruster, David Weetman, Carolyn S. McBride, Andrea Gloria-Soria, Jeffrey R. Powell, Bradley J. White

## Abstract

The number of dengue cases worldwide has increased ten-fold over the past decade as *Aedes aegypti*, the primary vector of this disease, thrives and expands its distribution, revealing limitations to current control methods. To better understand how *Ae. aegypti* evolved from a forest dwelling, generalist species to a highly anthropophilic urban species and the impact of contemporary gene flow on the future of dengue control, we sequenced 1,206 genomes from mosquitoes collected at 74 locations around the globe. Here we show that after evolving a preference for humans in the Sahel region of West Africa, the origin of the fully domesticated, anthropophilic subspecies *Ae. aegypti aegypti* (*Aaa*) occurred in the Americas during the Atlantic Slave Trade era and was followed by its explosive expansion around the globe. In recent decades, *Aaa* has invaded coastal Africa, the ancestral home range, introducing insecticide resistance mutations and an affinity for human hosts. Evidence of back-to-Africa migration is found in regions with recent dengue outbreaks, raising concern that global movement of *Aaa* could increase transmission risk of arboviruses including dengue in urban Africa. These data provide a platform to further study this important mosquito vector species and underscore developing complexity in the fight to limit the spread of dengue, Zika, and chikungunya diseases.

## Main text

The primary mosquito vector of dengue, *Aedes aegypti*, has evolved to thrive in and around human habitats and prefer human blood meals, putting nearly 4 billion people at risk of contracting dengue each year in tropical and subtropical regions (*1*). *Ae. aegypti* also transmits Zika, chikungunya, and yellow fever, and its distribution continues to increase (*2*), further expanding the public health burden caused by this mosquito. While the ancestral form of this mosquito, *Ae. aegypti* formosus (*Aaf*), has lived in the forests of Africa feeding on a variety of animals for the last 85k years (*3*), significant viral transmission to humans only began after the emergence of a human-preferring specialist subspecies, *Ae. aegypti aegypti* (*Aaa*) (*4*) in the Sahel region of West Africa within the last 5,000 years (*5*). Most dengue cases are transmitted outside of Africa by highly invasive *Aaa* populations that are genetically distinct from African *Ae. aegypti,* suggesting the expansion of *Aaa* around the globe followed a more recent shift that occurred within the last 500 years (*3*, *6*, *7*). The West African specialist populations can therefore be thought of as precursors to the widespread out-of-Africa specialist and referred to as *Ae. aegypti* proto-*aegypti* (proto-*Aaa*; (*4*)). We sequenced *Ae. aegypti* genomes from around the globe to identify the historical shifts and genetic changes underlying the explosive global invasion of this species that led to the global scourge of viral infections we face today. Following a similar effort in *Anopheles gambiae* (*8*), we named this endeavor “the Aaeg1200 genomes project”.

### Aaeg1200 genome sequencing dataset

We generated 31.4 terabases of DNA sequence with the Illumina HiSeq platform from 1,304 specimens representing 74 populations from throughout the global distribution of *Ae. aegypti* (Fig 1A). Special emphasis was given to sampling *Aaf* inside the native sub-Saharan African range of this species, with 510 samples representing 32 populations of this subspecies. A subset of our full dataset, including many of the *Aaf* populations, has been described previously (*9*). Short-read data were aligned to the updated versions of the AaegL5 (*10*) genome assembly (see Methods), resulting in an average read depth at accessible, robust sites of 11.24X across the genome (Figure S1). After quality control and removal of closely related individuals, the panel included 1,206 individuals. We identified 141.42 million single nucleotide polymorphisms (SNP) across the panel. Our dataset represents a significant step forward in genomic resources for this public health pest.

**Figure 1.**
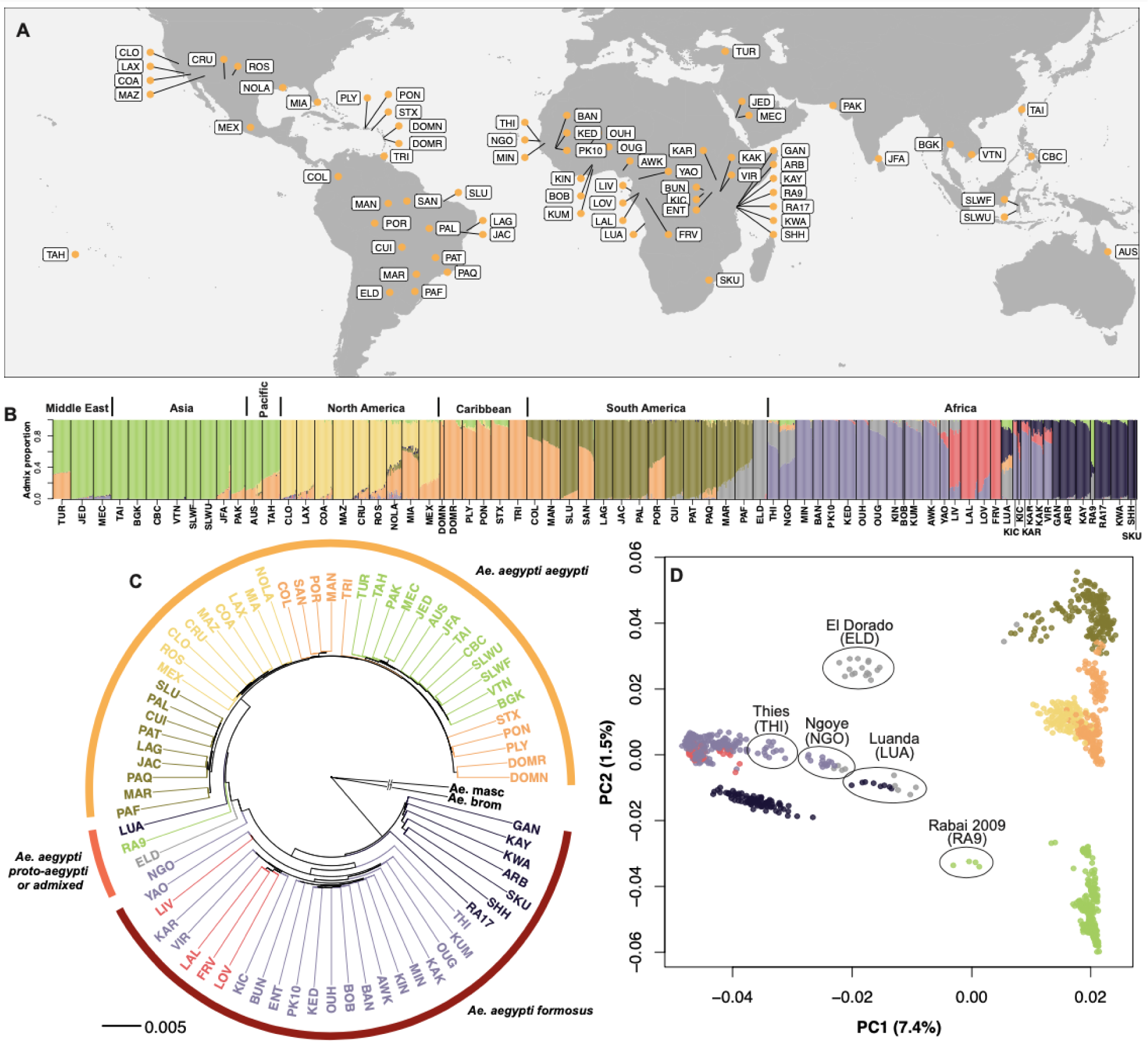
A) Global map showing locations of population samples included in Aaeg1200 sequencing. Details for individual population samples and full names can be found in Table S4. B) Barplot showing admixture proportions assuming eight ancestral populations (K=8). Each vertical bar shows admixture proportions for an individual mosquito with each color corresponding to an ancestral population. C) Neighbor-joining tree calculated based on pairwise genetic distance (*D_xy_*) among three representative individuals from each population and averaged at the population level. Outgroup branches are broken for ease of visualization. Branch and label colors correspond to admixture plot colors and outer band indicates subspecies designation with colors corresponding to Figure 2D. D) Principal Components Analysis of genetic variation in all 1,206 individuals. The first two principal component axes are shown with the percent of genetic variation explained shown in parentheses. Each dot represents one individual mosquito with color indicating largest admixture component from B).

### Genetic diversity and differentiation

Genetic diversity is concentrated within *Aaf,* with the majority of variants (53.3%) segregating exclusively in *Aaf*, 14.7% found only in *Aaa*, and close to a third (32.0%) shared between the two subspecies (Figure S2). Nucleotide diversity in *Aaf* varies along each chromosome with diversity lowest around the centromeres and increasing with distance from centromere until it reaches a plateau at the center of each chromosomal arm (Figure S3), a pattern characteristic of the interaction between natural selection and recombination along chromosomal arms (*11*). Nucleotide diversity in *Aaa*, in contrast, lacks this distinctive pattern with far less difference in diversity levels among the chromosomal arms and centromeric regions (Figure S4), suggesting a major role of demographic events impacting diversity patterns in these populations.

Large shifts in the frequency pattern of derived mutations in a population indicate historical changes in population size. We summarized derived SNP frequencies in each population and revealed strong deficits of rare SNPs in many *Aaa* populations compared to *Aaf* populations and a hypothetical population with a constant population size (Figure S5). We also measured correlations among SNP frequencies as a function of distance, or linkage disequilibrium, and found evidence of disequilibrium at more than two times greater distances in *Aaa* populations (mean LD decay distance in *Aaa* = 14,571 bp) than in *Aaf* populations (mean LD decay distance in *Aaf* = 5,669 bp; Figure S6). Both results suggest a history of strong population bottleneck events in *Aaa* populations.

To understand how populations are related to each other, we estimated genetic ancestry and genetic exchange among populations. Genetic structure analysis of the Aaeg1200 panel primarily separates *Aaf* and *Aaa*, with substantial differentiation between *Aaa* in the Americas and the rest of the non-African populations and also *Aaf* on either side of the East African Rift Valley (Figures S7 and S8). The (delta)K method (*12*) supports a model with five genetic clusters (max Delta K = 6.27), but model likelihoods increase continuously as more genetic clusters are added with the most complex model we evaluated (14 genetic clusters) being the most likely, consistent with differentiation among regional and local populations in the data. Several populations emerge as genetically intermediate between the large genetic clusters (Figures 1B,C,D) including Ngoye (Senegal), Dakar/Thies (Senegal), Luanda (Angola), Rabai (2009, Kenya), and El Dorado (Argentina), which could be explained by a simple serial founder scenario describing the origins of proto-*Aaa* and *Aaa* or a model with a simple split between *Aaa* and *Aaf* followed by admixture among the subspecies. Admixture models with eight or more genetic clusters identify the population from Argentina as a nearly ‘pure’ sample of proto-*Aaa*, with Ngoye, Dakar, Luanda, and others from West Africa being identified as mixtures of an *Aaf* admixture component and the proto-*Aaa* component (Figure 1B). Consistent with previous descriptions (*3*, *13*–*15*), Rabai (Kenya) is identified as an admixture of East African *Aaf* and an Asian *Aaa* cluster (Figures 1B,C,D).

### Origin of invasive form

Genetic studies suggest all invasive populations of *Aaa* are monophyletic with a single origin (*3*, *6*, *7*, *9*, *16*), but when and where invasive *Aaa* first evolved remains a key unanswered question. However, historical context must be considered as *Aaa* was partially or completely eradicated from large regions of the Americas and the Caribbean (*17*). *Ae. aegypti* populations currently found in these regions result from a massive re-invasion after eradication efforts ceased in the 1960’s, so they are unlikely to represent direct descendants of pre-eradication populations (*17*). The exceptions were some *Aaa* populations from the US and certain Caribbean islands that were not eradicated and therefore may represent early invasive *Aaa* populations (*17*, *18*).

To understand where and when invasive *Aaa* emerged, we first need to identify their closest ancestral population(s). Our data show higher genetic diversity and proto-*Aaa* ancestry in an Argentinian population of *Ae. aegypti* consistent with recent studies (*7*, *19*), suggesting the origin of *Aaa* could be described by an *Aaf*-to-proto-*Aaa*-to-*Aaa* stepping-stone model with American proto-*Aaa* as a middle step. However, populations of proto-*Aaa* outside of Africa could represent either a relict population that has survived since an introduction during the Atlantic Slave Trade (AST) or a recent introduction from Africa (*4*, *7*). We conducted cross-coalescent analysis to estimate the split time between El Dorado (Argentina) and Ngoye (Senegal), a proxy for African proto-*Aaa*, and found the timing of the split (Figure 2A) is most consistent with Argentinian populations representing relict populations of proto-*Aaa* introduced around the time of the AST. We also compared El Dorado to Luanda, another African proto-*Aaa* population (*4*, *20*, *21*), and found very similar patterns (Figure S9), confirming the split timing between American and African proto-*Aaa* populations is not sensitive to the choice of African proto-*Aaa* population. The similarity among cumulative migration curves points to close ties among the three proto-*Aaa* populations in our dataset prior to and during the AST. To test whether any of the proto-*Aaa* populations is more closely related to invasive *Aaa* populations using an alternative method, we calculated phylogenetic trees in 10 kb windows along the genome using a set of representative *Aaf*, proto-*Aaa*, and *Aaa* populations and found support for all three populations with 0.41 (95% CI: 0.40-0.42) trees placing Luanda closest to *Aaa*, 0.32 (95% CI: 0.31-0.33) trees placing El Dorado closest, and 0.27 (95% CI: 0.26-0.28) trees placing Ngoye closest. Cross-coalescent analysis and subtree weightings agree that El Dorado is not a recent introduction, but instead appears to be a relict of a highly connected metapopulation of proto-*Aaa* that was spread around the Atlantic basin around the time of the Atlantic Slave Trade.

**Figure 2.**
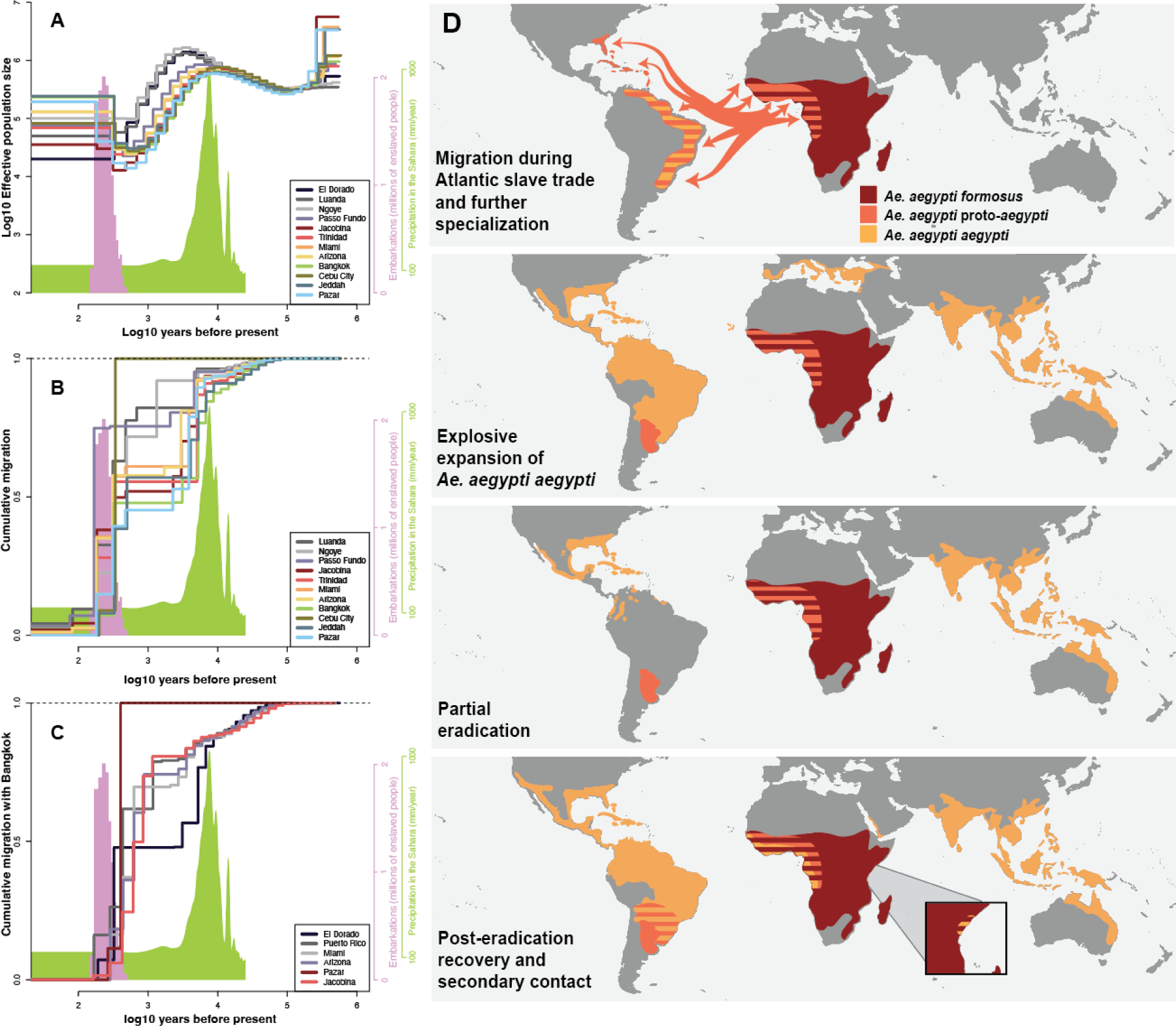
A) Cross-coalescent analysis of effective population size and migration using MSMC2. Effective population size plotted on log10 scale for representative populations as a function of years before present, also on a log10 scale, assuming 12 generations per year. The pink histogram shows the estimated number of slave vessel embarcations scaled by millions of enslaved people (Slave Voyages 2023) (*33*) on the alternative, pink y-axis at right. The green histogram shows precipitation levels in the Sahara (*34*) according to the green scale at right. Both histograms shown for approximate temporal reference. B) Cumulative migration plotted as a function of time on log10 scale (assuming 12 generations per year) between representative populations and El Dorado (Argentina) estimated using MSMC-IM. Cumulative migration is expected to plateau at one going back in time. Pink and green histograms are the same as for A). C) Same as for B) but cumulative migration between representative populations and Bangkok. D) Phased map showing simplified distributions and movements of different subspecies and forms of *Ae. aegypti* according to color. Description of each phase is shown in the bottom left corner of each phase. Orange arrows in the first phase indicate movement of proto-*Aaa* during the AST. Striped areas indicate more than one form or subspecies found in that region at that time.

If proto*-Aaa* populations were introduced during the AST and invasive *Aaa* evolved from proto-*Aaa* in the Americas, then we would expect the split between proto-*Aaa* and invasive *Aaa* populations to come after the split among proto-*Aaa* populations. We estimated the timing of the split between El Dorado proto-*Aaa* and several representative *Aaa* populations using MSMC-IM and discovered support for multiple histories (Figure 2B). The *Aaa* population from Cebu City (Philippines) *Aaa* showed a sharp increase in cumulative migration from ∼0.1 to one going backwards in time around the time of the AST, implying complete population merging with El Dorado at that time. Passo Fundo, Brazil, on the other hand, shared migrants with El Dorado more recently than did Cebu City, but approximately one quarter of the Passo Fundo genome coalesced further back in time on the order of thousands of years, not hundreds. The remaining *Aaa* populations showed similar step-functions in their cumulative migration curves where a significant proportion (∼0.4-0.6) of the genome coalesced with El Dorado around the time of the AST, but the remaining genomic haplotypes remained isolated until around the same time that Passo Fundo showed an ancient increase in migration. We evaluated the robustness of these cumulative migration curves by repeating the analyses with bootstrap resampled genomes and found that while some populations show similar curves for both the empirical and bootstrap replicates (e.g. Arizona, Miami, Pazar, Figure S10), many *Aaa* populations showed both the step-function shape as well as recent population merging with El Dorado similar to Cebu City where the curve reaches one around the time of the AST (Figure S10). We also calculated cumulative migration curves comparing Arizona and Trinidad with Luanda, another proto-*Aaa* population, and recovered both shapes in the bootstrap replicate curves (Figure S11). These results suggest that genomes from *Aaa* and proto-*Aaa* populations comprise a mixture of haplotypes with different histories: haplotypes that coalesce between *Aaa* and proto-*Aaa* around the time of the AST, and haplotypes that coalesce further back in time, around the time when proto-*Aaa* split from *Aaf* ∼5,000 years ago (*5*). The presence of haplotypes in proto-*Aaa* and *Aaa* genomes with such distinct histories could be explained by either a history of substructure within African proto-*Aaa* with variable levels of admixture exchange with *Aaf* or introduction of *Aaf* haplotypes through more recent admixture with proto-*Aaa*. The similarity between cumulative migration between Arizona and both El Dorado and Luanda suggests that recent admixture among proto-*Aaa* and *Aaf* cannot explain the entire signal since El Dorado is geographically isolated from *Aaf*.

Comparing estimated effective population size changes over time can also provide information on when two populations last exchanged migrants (*22*). The effective population size of El Dorado diverges from invasive *Aaa* populations shortly after the African Humid period ∼5,000 years ago (Figure 2A and S12), significantly pre-dating the AST. Notably, invasive *Aaa* populations share the signature of population size increase after the bottleneck when their effective population sizes increased while El Dorado’s effective population size remained at the bottlenecked size (Figure 2A). However, historical migratory exchange among partially isolated proto-*Aaa* populations within could also lead to higher inferred effective population size since some alleles could have spent time in other subpopulations leading to longer coalescent times that could impact effective population size inference (*23*). Alternatively, the extreme demographic shifts in the recent history of invasive *Aaa* populations could have led to loss of some older haplotypes obscuring historical migration events and effective population size shifts. To evaluate the possibility of ancient substructure in proto-*Aaa*, we returned to subtree analysis and quantified the prevalence of a star-like phylogeny specifically testing for topologies where *Aaf*, proto-*Aaa*, and *Aaa* are all reciprocally monophyletic. Although recent secondary contact events and ongoing lineage sorting would both obscure this pattern, we found only 0.0116 of windows show a star-like phylogeny. Together with the historical absence of *Aaa* populations within Africa until recently (*13*, *24*), the most parsimonious scenario is that proto-*Aaa* emerged in West Africa with varying degrees of isolation from *Aaf*, Slave Trade vessels transported populations of proto-*Aaa* to the Americas during the AST, and *Aaa* evolved from proto-*Aaa* in the Americas and spread around the globe.

Historical records and genetic data suggest that Asia, the Pacific region, and the Middle East were colonized by *Aaa* in the 1800’s, well after *Aaa* establishment in the Americas (*7*, *25*), but the source of these populations is not well understood. Our genetic structure analysis, PCA, and genetic distance tree all show that populations from the Middle East, Pacific, and Asia have genetic affinities with populations from the Caribbean (Figure 1), potentially consistent with movement from the Caribbean to these regions as the source of the founding populations. Cross-coalescent analysis of migration, however, comparing the population from Bangkok with representative *Aaa* American populations shows that Bangkok split from all the representative populations around the same time, although the population from Puerto Rico exchanged migrants with Bangkok more recently than the other populations, likely explaining the higher level of shared ancestry with populations from the Caribbean (Figure 2C and Figures S13). This hypothesis is consistent with the presence of *Ae. aegypti* in the Mediterranean Basin in the 1800s (*20*, *26*), near ports involved in trade with the Americas that frequently stopped at Caribbean Islands (especially Santo Domingo) before returning to Europe, providing a potential route for Aaa introduction to Asia through the Suez Canal, although *Aaa* could also have been introduced through trade channels traveling around the horn of Africa to India and Japan.

### Contemporary admixture between *Aaa* and *Aaf*

After historical shipping and trade helped spread *Aaa* throughout the tropical world, modern shipping, trade, and travel have increased global connections with potentially important implications for mosquito populations and viral transmission risk. Our admixture analysis suggests that *Aaa* populations from Southern Brazil are admixed with proto-*Aaa* ancestry, likely from nearby Argentina (Figure 1B). Following the collapse of eradication efforts in South America in the 1960’s, *Aaa* populations likely reinvaded from the North (*27*), and reestablishing throughout eradicated areas, potentially making secondary contact with relict proto-*Aaa* populations upon expansion. We analyzed admixture tracts along genomes from Passo Fundo from Southern Brazil to estimate the timing of admixture between *Aaa* and proto-*Aaa* from El Dorado (Figure S14). We modeled Passo Fundo as a mixture of El Dorado and the closest *Aaa* population, Patos de Minas, and estimated the time of contact to be ∼22 years ago (assuming 12 generations per year, 95% confidence interval 18-27). Considering our Passo Fundo samples were collected in 2017, the admixture we detected occurred in the 1990’s. It is possible that during this time period *Aaa* completed reinvasion from the north into Southern Brazil. Alternatively, the establishment of Mercosur, an economic and political bloc consisting of Argentina, Brazil, Paraguay, and Uruguay, increased trade in this area since 1991, resulting in increased migration and secondary contact between proto-*Aaa* and *Aaa*.

We also searched for contemporary admixture between *Aaa* and *Aaf* populations more broadly using Patterson’s F3. Admixture signals were identified predominantly in coastal populations on both coasts of Africa, with particularly strong signals detected in coastal Senegal, Angola, and Kenya, whereas Ouagadougou, Burkina Faso, was the only inland population with clear evidence of admixture (Figure 3). These results were robust to selection of alternative *Aaa*/*Aaf* ancestral populations (Figure S15). We applied Patterson’s F4 to pairs of *Aaa*/*Aaf* and pairs of proto*-Aaa*/*Aaf* to rule out the possibility that the admixture signals derived from admixture between proto-Aaa African populations and ‘pure’ *Aaf* African populations and find in all cases that the magnitude of F4 was greatest within *Aaa*/*Aaf* admixture pairs (Figure S16), and could not derive from admixture within sub-Saharan Africa. To understand the timing of admixture, we fit a two-pulse admixture model to identify admixture tracts in samples from Rabai, Kenya, and find evidence for first contact to be ∼52 years (assuming 12 generations per year, 95% Confidence Interval 39-71 years) before sample collection and a secondary wave of admixture ∼10 years before sample collection (95% Confidence Interval 7-18 years). The first report of *Aaa* along the coast of East Africa was in 1952, consistent with the presence of an imported domesticated population (*28*). Together, our admixture results indicate that contemporary secondary contact between global *Aaa* and African *Aaf* is extensive in coastal regions.

**Figure 3.**
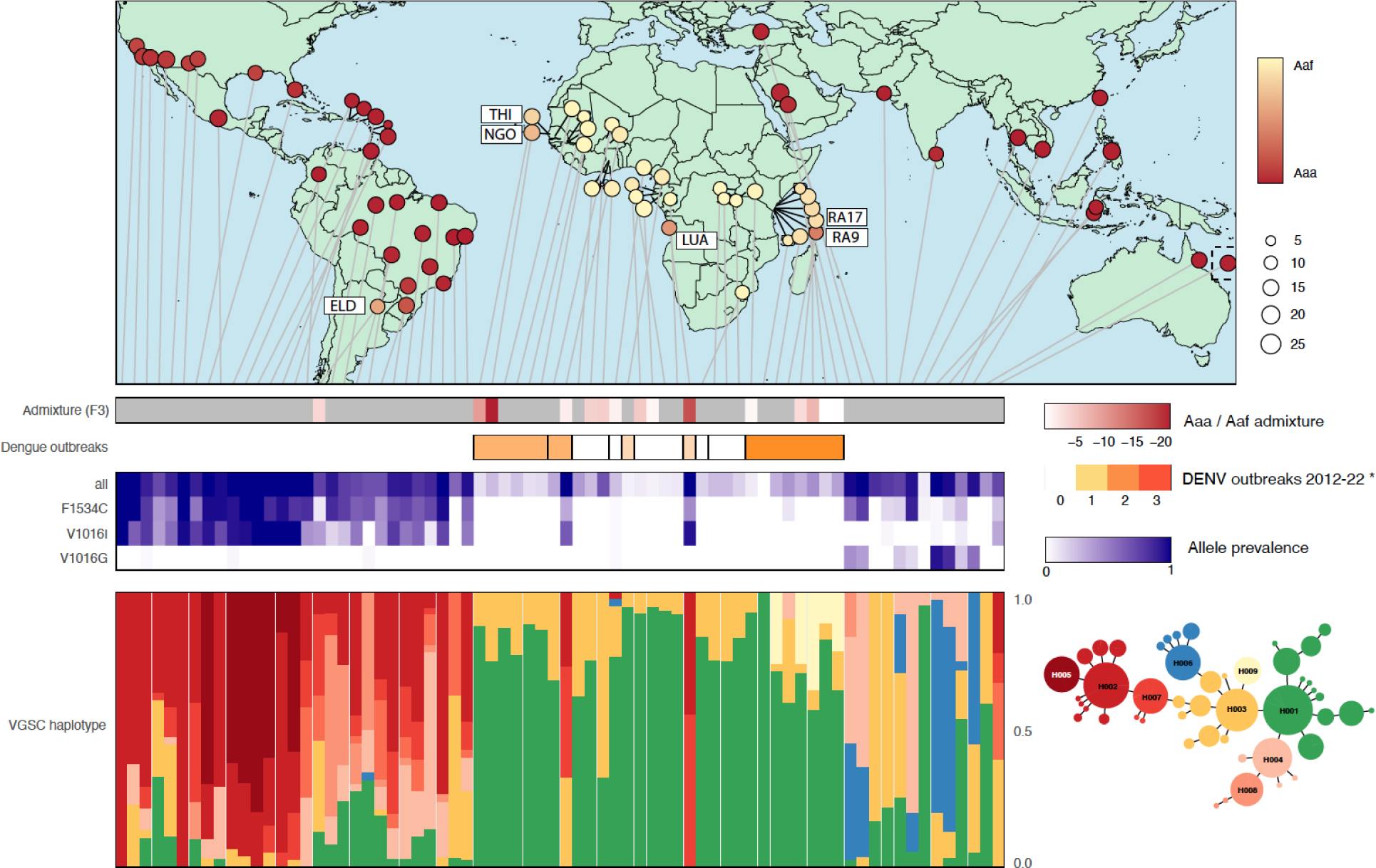
Global map showing evidence of *Aaa/Aaf* admixture proportions from NGSadmix analysis with K=2. Lollipop markers used to offset populations when actual locations were too tightly clustered for visualization, with notable populations given labels as in Figure 1. Light gray lines indicate position of each population in barplots below. The first row shows results of the F3 test for admixture according to scale at the right with darker purple corresponding to stronger evidence of admixture. The third panel shows the number of dengue virus outbreaks in Africa according to Gainor, Harris, and LaBeaud (*30*). The fourth panel shows population-level prevalence of KDR resistant mutations as labeled on the left with colors indicating prevalence according scale on right. The bottom panel shows population-level frequency of each of the top seven resistant KDR haplotypes (labeled H002-H008), the susceptible haplotype (H001), and minor haplotype clusters with colors indicating related haplotypes in the network on the right. See Figure SX for detailed KDR haplotype network.

### Knock-Down resistance mutation sharing

Pyrethroid chemical insecticides are a critical component of modern mosquito control, but the spread of insecticide resistance mutations is threatening the utility of this tool globally. We find multiple SNPs that have demonstrated roles in pyrethroid resistance at high frequency in our dataset (Table 1). Knock-Down Resistance (KDR) SNPs are common in the Americas, with high F1534C prevalence across all regions (USA 76%; Mexico 97%; Brazil 64%), and above 90% frequency across the Caribbean (Table S1). Two mutations at V1016 are mutually exclusive, with V1016I predominating in the Americas, often in linkage with F1534C and V1016G at high frequency across Asia, consistent with earlier reports (*29*).

**Table 1:** Common KDR resistance mutation frequencies summarized by geographical region. Values in parentheses indicate frequency range among populations within each region. Population-level frequencies can be found in Table S1.

| Region | F1534C | V1016I | V1016G | All resistant haplotypes |
| --- | --- | --- | --- | --- |
| East Africa | 0.01 (0-0.1) | 0 (0-0) | 0 (0-0) | 0.23 (0-0.43) |
| West Africa | 0.04 (0-0.44) | 0.09 (0-0.94) | 0 (0-0.03) | 0.29 (0.05-1) |
| Caribbean | 0.92 (0.81-1) | 0.98 (0.94-1) | 0.01 (0-0.03) | 1 (1-1) |
| South America | 0.64 (0-0.97) | 0.4 (0-0.78) | 0 (0-0.03) | 0.84 (0.59-1) |
| North America | 0.77 (0.39-1) | 0.76 (0.37-1) | 0 (0-0.03) | 0.93 (0.72-1) |
| Middle East | 0.66 (0.55-0.77) | 0 (0-0) | 0.4 (0.37-0.43) | 0.99 (0.98-1) |
| Asia | 0.18 (0-0.89) | 0.01 (0-0.07) | 0.32 (0-0.94) | 0.76 (0.08-1) |
| Pacific | 0.11 (0-0.22) | 0.17 (0-0.33) | 0 (0-0) | 0.57 (0.41-0.72) |

Within Africa, many of the populations with strong signals of *Aaa*/*Aaf* introgression also exhibit a high prevalence of KDR resistance SNPs (Figure 3); F1534C and V1016I are both at high frequency in Luanda, Angola (F1534C: 94%; V1061I: 44%) and Ouagadougou, Burkina Faso (65%; 33%). However, this pattern is not universal, with admixed coastal populations in Senegal showing a complete absence of the three major KDR resistance alleles, despite strong evidence of admixture.

To infer the origins and spread of KDR-related pyrethroid resistance in *Ae. aegypti,* we derived a haplotype network from all non-synonymous SNPs across the locus (Figure 3 and Figure S17). While the susceptible (wildtype H001) haplotype represents 31% of the dataset, just eight mutated haplotypes make up the next 58% of the total, each of which contains at least one previously characterized resistance-associated SNP (Figure S18). In all regions outside Sub-Saharan Africa, susceptible haplotypes are in the minority; the average North American population exhibits 93% resistant haplotypes (72-100%), and the mean frequency of resistant haplotypes is similarly high across Asia, albeit across a much wider range (mean 76%; range 8-100%) (Table S1). Stark haplotype frequency differences between Asia and the Americas and the topology of the network support independent origins of pyrethroid resistance in each continent, in agreement with previous reports (*29*).

The prevalence of resistance haplotypes within Sub-Saharan Africa is consistently lower than in the rest of the world, with mean prevalence of resistant haplotypes in West and East Africa being 23% and 29%, respectively, and many populations carrying an entirely susceptible KDR profile. Resistant haplotype H009, which contains only the I915K SNP, is only found in East Africa and may represent a reversion of the locus.

## Discussion

The World Health Organization (WHO) has recorded a ten-fold increase in global dengue cases over the last two decades with over five million cases and 5000 deaths in 80 countries and territories in 2023 alone (*1*). Effective vaccines against dengue, chikungunya, and Zika are not widely available, so control of mosquito vectors is critical to reduce the impact of these diseases on human populations. Improved understanding of the genetic factors enabling the global spread of *Ae. aegypti* including the genetic mechanisms underlying insecticide resistance and their spread, variation in viral transmission capacity, and preference for human bloodmeals, will improve the efficacy of existing tools and facilitate development of new control tools. Investing in biological resources such as an improved reference genome and a database of over 1,200 full genome sequences provides a valuable platform for future studies of *Ae. aegypti*.

The evolution of preference for human hosts and adaptations to human habitats set the stage for the explosive expansion of *Aaa* and increased risk of arboviral diseases around the globe. Recent work showed that the first step in this evolution likely began in the Sahelian region of West Africa approximately 5,000 years ago (*5*), in close proximity to the ancestral range of *Aaf*, but our data show that an additional evolutionary step took place within the Americas.

Cross-coalescent analysis of a population in Argentina and populations in West Africa reveal the presence of a proto-*Aaa* population in the Americas dating back to the time of the AST. Genetic relatedness results, stepwise reduction in genetic variability, and cross-coalescent analyses all point to a *Aaf*-to-proto-*Aaa*-to-*Aaa* stepping stone model, but cross-coalescent and subtree weighting analyses show that the a simple stepping stone model does not capture all the complexities in the data. Our results suggest that proto-*Aaa* may have existed as a series of subpopulations with differing levels of admixture with *Aaf*, and the proto-*Aaa* populations in our dataset are imperfect proxies for the ancestor of *Aaa*. Moreover, current and historic admixture could further complicate cross-coalescent analyses by introducing haplotypes with histories that differ from the history of the focal population. Taken together, our results suggest that the period leading up to and during the AST was characterized by partial separation between proto-*Aaa* and *Aaf* followed by large scale movement and shuffling of proto-*Aaa* populations around the Atlantic coasts of Africa and the Americas introducing populations to an open, human-centered, niche in the the Americas. Canonical *Aaa* then emerged in the Americas leading to rapid expansion throughout the continent and eventually the rest of the tropical world. We screened for signatures of natural selection potentially involved in the transition from proto-*Aaa* to *Aaa* and found putative evidence for adaptation to novel pathogens and feeding environments (Supplemental Text, Figure S19) consistent with niche expansion after the emergence of invasive *Aaa*.

Our analysis indicates that the global spread of *Aaa* continues today, with important implications for control of *Aa. aegypti* and dengue transmission, especially in Africa. Back-to-Africa migration of *Aaa* was first recognized in Tanzania in the 1950s (*28*), and our data suggest such movement of the domestic, human-preferring *Aaa* into Africa is more widespread and common than previously understood. Dengue outbreaks have become more common in Africa and are geographically correlated with the signatures of *Aaa* admixture in coastal regions (Figure 3). While most of Sub-Saharan Africa does not report extensive dengue outbreaks, four out of seven locations that show signals of *Aaf/Aaa* admixture (Angola, Burkina Faso, Kenya, and Senegal) have reported regular or significant dengue outbreaks in recent years (*30*). We also show insecticide-resistant haplotype sharing among populations in the Americas and populations in Africa, pointing to the importation of genetic variants that could hamper mosquito control efforts. The combined effects of increased urbanization and global connectivity have led to an invasion of the ancestral range by the highly domesticated *Aaa* that may already be driving increased dengue transmission (*31*, *32*).

Scientific resources for studying *Ae. aegypti* have lagged behind *Anopheles* malaria vectors, but the Aaeg1200 genome sequence data set adds to the improved genome reference sequence available (*10*) to provide a platform to continue studying this important mosquito disease vector, including evolutionary analysis of neo-sex chromosome formation and chromosomal inversions (Supplementary Text, Figures S31,S32,S33). Our analysis provides answers to some previously unanswered questions, but raises additional questions and only touches the surface of what can be studied with these data. Combined with valuable field and laboratory research, we hope that these data will move the field forward and contribute to the fight against *Ae. aegypti* and the viruses this species transmits to millions of humans around the globe.

## Supporting information

TableS2.PBS_Scan_for_Natural_Selection

TableS4.Population_Sample_Information

## Methods and Supplemental Text

### Sample Collection

Mosquito samples were provided by a number of contributors either as carcasses or DNA extractions. Samples were collected from 2009-2019. See Table S4 for sample sizes, location, and other collection information.

### Whole genome re-sequencing and processing

We sequenced whole genomes using the Illumina HiSeq X and 4000 platforms. When adult or larval carcasses were available, we extracted DNA using the protocol described previously in Crawford et al. (*1*). Briefly, individual carcasses were homogenized in lysis buffer using steel grinding balls on a SPEX grinder. Lysate was transferred to a new plate as input for DNA extraction on Chemagic360 according to the manufacturer’s protocol. DNA sequencing library preparation was performed using a PCR-free protocol also described in detail in Crawford et al. (*1*). Briefly, at least 150 ng of gDNA from each sample were fragmented to target size between 350 and 400 basepairs, double-side size selected to narrow the size range to 350-400 basepairs, end repaired, A-tailed, and adapter-ligated. Purified libraries were pooled and sequenced on the Illumina HiSeq 4000 and HiSeqX platforms to generate 2x150 basepair reads.

DNA sequence reads for samples published in Rose et al. (*2*) have been deposited to NCBI SRA under accession number PRJNA602495. The remaining samples have been deposited under accession number XXXXXX.

### Short-read mapping and genome reference updating

We mapped Illumina paired-end reads to the *Ae. aegypti* L5 genome reference (NCBI WGS Project NIGP01; (*3*)) using BWA *mem* (v0.7.16)(*4*) with default parameters, then sorted using samtools *sort* (1.1)(*5*), and duplicate reads were marked using Picard’s *markdup* function (2.1.0, http://broadinstitute.github.io/picard/).

To minimize the effects of SNP variants and genetic distance on short read mapping, we conducted an iterative process to update the reference sequences. This routine was described previously in Rose et al. (*2*), and the resulting reference sequence was used for mapping *Aaf* populations for this study as well. We analyzed various aspects of read mapping for each iteration and found that the number of reads mapping, the number of sites covered, and the number of heterozygous sites discovered all increased, while mismatches between reads and the reference decreased, as shown in Figure S5 in Rose et al. (*2*). We made an analogous updated reference specifically for *Aaa* populations as well using the same method. Briefly, a random selection of 100 *Aaa* males were aligned to the L5 *Ae. aegypti* reference, and then the reference was updated with the consensus allele from the pileup. This process was repeated three times and then used this updated reference for remapping all *Aaa* samples.

After re-mapping all short reads to updated reference sequences, we conducted indel realignment and read clipping. We used GATK (v3.8-1-0-gf15c1c3ef)(*6*) for indel realignment on each population individually with default settings. After indel realignment, we clipped overlapping read pairs using the clipOverlap function of the bam program within the bamUtil analysis kit (v. 1.0.14) (*7*).

### Outgroup sequences

We used publicly available sequence data for *Aedes bromeliae* to assemble an outgroup sequence. We downloaded fastq files (accession numbers SRX2323182, SRX2323157, and SRX2323117) from NCBI SRA for an *Ae. bromeliae* individual. These short reads were aligned to the updated *Aaf* reference sequence using the same pipeline described above including read clipping, but no indel realignment. We created a fasta sequence from the short data using ANGSD (-minMapQ 20 -minQ 10 -only_proper_pairs 1 -remove_bads 1 -uniqueOnly 0 -doFasta 3 -doCounts 1 -setMaxDepth 150 -setMinDepth 8).

We sequenced four *Aedes mascarensis* individuals from a laboratory colony using the same protocols as for the full *Ae. aegypti* panel.

### Close relative screen

To identify close relatives within each population sample, we calculated relatedness among all individuals using NgsRelate (*8*). GLF input files were prepared using ANGSD (*9*) at 60 Million randomly selected sites from the robust sites set described below and GATK genotype likelihood model (-GL 2), as recommended by the authors of NgsRelate. We further limited the analysis to sites with read depth greater than seven reads and less than 46 reads, and applied the SNP-pval flag with a value of 1e-6. We ran NgsRelate with a minimum allele frequency of 0.05 and plotted the KING-robust and R1 statistics to evaluate relatedness.

To confirm the KING-robust and R1 statistics behaved as expected with our samples, we calculated the physical distance between samples, all *Aaf*, collected for Rose et al. (*2*) based on the location of ovicups used for collections. We binned sample pairs into four categories: 1) cupmates collected from the same ovicup, 2) pairs of samples from different cups but less than 50 meters apart, 3) pairs of samples from cups separated by 50-100 meters, and 4) pairs of samples from cups greater than 100m apart. We plotted KING-robust on R1 for each category separately (Figure S20). As expected, close relatives (KING-robust > 0.25 and R1 > 0.5) were predominantly found in the cupmate and <50m categories with only two exceptions in each of the other categories.

We then plotted the same statistics for all populations (Figure S21) with *Aaa* and *Aaf* populations separated by color. In contrast to *Aaf* populations where most pairs had KING-robust and R1 values consistent with low relatedness (lower left quadrant), *Aaa* populations followed a different pattern with elevated R1 values overall and a near continuous distribution extending into the high-relatedness quadrant (Figure S21). Although the relatedness statistics followed the expected distribution in *Aaf* populations, the statistics appeared elevated in *Aaa* populations. The authors of NgsRelate conducted simulations to show that these statistics can be robust to non-equilibrium demographics, but we suspect that the demographic histories of the *Aaa* populations likely deviate substantially from the relatedness models underlying KING-robust and R1. As such, we used the standard thresholds (KING-robust < 0.25 and R1 < 0.5) to identify unrelated pairs and removed one individual of every pair that fell outside those bounds for all *Aaf* populations but not *Aaa* populations. Relatedness filtering resulted in the removal of 98 individuals from *Aaf* populations leaving 412 *Aaf* and 794 *Aaa* individuals used in all downstream analyses.

### Robust site identification

For each analysis, we identified genomic locations considered robust for analysis by applying a series of filters and only keeping sites that passed all filters. First, we generated a pileup VCF annotated with a series of quality statistics and tests using samtools mpileup (1.6-2-gf068ac2) and the following flags: ‘-q 10 -Q 20 -I -u -t SP,DP’. Putative variant sites were called from this VCF using BCFtools (1.6, https://samtools.github.io/bcftools/) with ‘-f GQ -c’ flags, which were then filtered using SNPcleaner.pl from ngsQC (https://github.com/tplinderoth/ngsQC) with the following settings ‘-k MIN_IND -u 2 -D MAXD -a 0 -f 1e-4 -H 1e-6 -L 1e-6 -b 1e-10’. For population samples with more than 12 individuals, MIN_IND, or the minimum number of individuals with data at a given site, was set to 12. For population samples with less than 12 individuals, MIN_IND was set to the number of individuals in the sample. The total maximum depth across all individuals in a population sample, MAXD, was set to 45 ⨉ MIN_IND. In all cases, the autosomal genome was divided into 10 MB chunks and analyzed in parallel using GNU parallel (*10*). This pipeline produced a set of sites, including both putative variant and invariant sites, for each population separately that have passed all of the filters that could be considered robust for analysis of that population. To obtain a global set of robust sites for analysis of the full panel, we found the intersection of robust sites sets across all *Aaa* populations and *Aaf* populations separately.

To further filter sites that are potentially difficult to map in the genome, we calculated the total depth of reads with map quality >= 20 for all African and non-African samples on their respective updated references at sites identified in the robust set above. Supplementary Figure S22 shows density curves for each set up to 20k reads. For individuals mapped to the African reference, 1,180,248,427 (98.76%) sites had at least one read. For the non-African reference, 1,169,822,108 (97.89%) sites were covered by at least one read. While both sets show clear local maxima at 6410 and 9701 for African and non-African, respectively, they differ in the density of lower-depth sites, with a sharp increase in lower-depth sites in the African set. Regional genomic biases in accessibility impacting read mapping rates likely contribute substantially to variation in read depth across the genome. We excluded sites in the African set with a total read depth greater than 50% above the 6,410 maximum, or 9,615 reads. Using the same logic, we excluded sites with a total read depth greater than 14,552, for the non-African set. High read-depth sites were excluded from the *Aaa* and *Aaf* robust sites sets from above and then the filtered sets were merged to find the intersection to arrive at a final set of 1,138,636,693 sites for analysis.

### Variant calling

We called SNPs using the likelihood ratio test in ANGSD and a p-value threshold of 1e-6 in the *Aaf* (n=425) and *Aaa* (n=778) sets separately. We identified 120.68 million variant sites in *Aaf* and 66.02 million variant sites in *Aaa*. We filtered all variants to include only sites with data present for at least 80% of individuals and the minor allele was found on at least 10 chromosomes. We then found the union of the filtered *Aaf* and *Aaa* variant sets for each chromosome resulting in a total of 141.42 million unique variants.

In addition to the full set, we generated three linkage disequilibrium-pruned variant sets to facilitate genome-level analysis of independent loci. First, we used PLINK (*11*) to calculate linkage disequilibrium (LD) in 100 variant windows with a step size of and an LD cutoff of 0.1 resulting in a set of 7,161,365 from the Aaa variant set. From this LD-pruned set, we randomly selected one million sites (1e6 set). From the 1e6 set, we randomly selected a smaller set of 100,000 sites. For analysis, we filtered the gVCFs to generate a set of variant VCFs with each of these LD-pruned variant sets.

### Haplotype phasing

We phased genomes as described in Rose et al. (*12*), first pre-phasing variants spanned by single sequencing fragments using HAPCUT2 (*13*), then statistically phasing chromosomes using SHAPEIT4 (version 4.2) (*14*). In order to manage memory usage, we phased different global regions (North America, South America, African samples not previously analyzed in Rose et al. (*12*), and Asia) in separate SHAPEIT4 runs. We then used bcftools merge to combine phased regional genome sets. For each SHAPEIT4 analysis, we included the Rose et al. (*12*) set of phased genomes as a reference panel.

### Genetic differentiation and structure

We calculated genetic differentiation using multiple approaches. First, we used ANGSD to prepare a beagle-format genotype likelihood file at the unlinked 1M SNP set described above and used NGSadmix (*15*) to estimate admixture proportions assuming (K) 2-14 population components. We applied a minimum minor allele frequency threshold of 0.05, and missingness tolerance of 0.05, and a minimum number of individuals with a data filter of 1100 resulting in a filtered set of 292,710 SNPs for analysis. To compare among models with different numbers of population components, we ran each K level 20 times with different starting seed values and submitted output of all replicates to the CLUMPAK server for analysis using the best K method (tau.evolseq.net/clumpak/bestK.html). The output of both the Evanno (*16*) best-K and model probability analyses are shown in Figures S23 and S24, respectively.

Second, we used the same unlinked 1M SNP set for PCA. We calculated the covariance matrix using PCangsd (version 0.982, https://github.com/Rosemeis/pcangsd.git) (*17*) and -minMaf threshold of 0.01. Eigenvectors were calculated using the eigen function in R (*18*). For Figure 1, we found the largest admixture component from the admixture analysis with K=8 and set the color of the individual in the PCA plot to match the color in the admixture plot. We also plotted subsets of the full panel by first subsetting the covariance matrix and then recalculating the eigenvectors. Analysis of only non-African *Aaa* populations is shown in Figure S25 and proto-*Aaa* and *Aaf* populations are analyzed and shown in Figure S26.

Third, we calculated genetic distance as *D_xy_*. To calculate *D_xy_* while avoiding any biases associated with hard SNP and genotype calling, we used realSFS (*19*) and ANGSD to calculate 2D site frequency spectrum (SFS) pairwise between all individuals and used the entries of the 2D SFS to quantify the number of differences and the total number of sites. The 2D SFS was calculated at all robust sites after removing all sites inside and within 100 basepairs from boundaries of annotated exons. For computational efficiency and to avoid the sex-differentiated region on chromosome 1, we limited this analysis to 84 one megabase windows on chromosomes 2 and 3 excluding centromeric regions as defined for the AaegL5 assembly. We used this same protocol for the two outgroups, *Ae. bromelia* and *Ae. mascarensis*, and included them directly in the overall distance matrix. The neighbor-joining tree was calculated and plotted using the ape package (*20*) in R (*18*).

### Site frequency spectra

To avoid any biases associated with hard SNP calling, we calculated unfolded site frequency spectra (SFS) directly from sequencing read data using ANGSD and realSFS for each population separately. First, we calculated site allele frequency (.saf) files at global robust sites on all three chromosomes (unplaced scaffolds excluded) using *doSaf* in ANGSD with *Ae. bromeliae* specified as the ancestral sequence. Second, we used realSFS to estimate the genome-wide SFS for each population. For plotting, allele counts were converted to proportions to facilitate comparison among populations.

### Nucleotide diversity

We calculated nucleotide diversity (*π*) by calculating folded site allele frequency files for each population. We then used realSFS to estimate folded site frequency spectra, calculate theta statistics for each population using the thetaStat program (*21*) distributed with ANGSD, and conducted sliding window analysis with 50kb non-overlapping windows. We summarized *π* using smoothed-loess functions in R (*18*) and plotted along chromosomes 1-3 (Figures S3 and S4). We also calculated density curves for each population based on all 50kb windows (Figure S27) and summarized the distributions by calculating the mean and standard deviations across all windows for each population and plotted on the global map (Figure S28) using the R package *rworldmap* (REF).

### Linkage disequilibrium

We calculated *r^2^* using ngsLD (*22*). For efficiency, we extracted one megabase regions from each of the six chromosomal arms. On the p arms, the regions started at 110MB on all three chromosomes. On the q arms, the region started at 270 MB on chromosome one, and 320MB on chromosomes two and three. These regions were extracted from the full genome beagle and then each population was extracted to make population specific input files. ngsLD was run with the following flags --probs --max_kb_dist 50 --min_maf 0.05 --ignore_miss_data --rnd_sample 0.1 --extend_out to randomly select 10% of sites, apply a minimum allele frequency filter and limit comparisons to SNPs within 50 kb of each other. Decay curves were fitted using the fit_LDdecay.R script included with ngsLD and plotted (Figure S6).

### Cross-coalescent analysis

All these analyses were performed following the methodology outlined in Rose et al. (*12*). In summary, for the cross-coalescent analyses, we first utilized the bam files corresponding to each sample to generate a genome-specific mask for every chromosome and sample. This was achieved using the bamCaller.py script from the MSMC tools package (https://github.com/stschiff/msmc-tools). Subsequently, we obtained the final genotype for each sample by merging its phased genotype from the *Aedes aegypti* phased reference panel (as inferred in Rose et al. (*2*, *12*) with its original genotype obtained using samtools 1.9 (mpileup -B -q20 -Q20 -C50). Next, we filtered the SNPs using both the specific mask and the general mask (*2*), retaining only the best SNPs that met the following criteria: under non-selection between specialist-generalist, uncallable sites, absence from the sex locus or on repetitive and centromeric regions. Finally, we inferred the cross-coalescent events by conducting MSMC2 (*23*) analysis followed by MSMC-IM (*24*) analysis, to fit the isolation-with-migration model to the inferred cross-coalescent estimations. These analyses were performed with default parameters. Bootstrapped MSMC2 and MSMC-IM analyses were conducted using ten replicates of three 400 Mb chromosomes constructed with resampled blocks of 20 Mb. For plotting, we used the same approach as Rose et al. (*12*) including a scaling factor corresponding to a mutation rate (*μ*) of 4.85 x 10^-9^ and generation time (*g*) of 0.067 years. Effective population size and cumulative migration curves for bootstrap replicates are shown in Figures S9 - S13.

### Ancestry tracks analysis

We used AncestryHMM (*25*) to detect evidence of admixture over the genome of two different populations: Passo Fundo from Brazil and outdoor-Rabai collected in 2017 from Kenya

For Passo Fundo, El Dorado (Argentina) population was selected as a representative of the proto-*Aaa* ancestor and Patos de Minas (Brazil) as a proxy of *Aaa* as the closest, non-admixed Brazilian population to Passo Fundo.

For Rabai 2017 analysis, African mosquitoes from different populations but with a clean Aaf ancestry (see NGSadmix results: 3 from Arabuko, 4 from Kwale, 3 from Shimba Hills, and 2 from Skukusa) were used as representatives of *Aaf*, while individuals from Jeddah were used as the proxy for the *Aaa* donor population.

AncestryHMM was applied following the detailed instructions on (*12*), with minor modifications: after filtering in the best sites as for cross-coalescent analysis, the input data was obtained by considering a recombination rate of 1.7e-9 M/b. The admixture timing was estimated for each comparison. An analysis of 100 bootstrap samples with blocks of 2000 informative sites was used to infer the 95% CI for the admixture timing.

### Subtree weighting analysis

To quantify support for various phylogenetic relationships along the genome, we used a method similar to *Twisst (26)* where we built neighbor-joining trees based on genetic distance among a representative set of individuals within 10kb windows. First, we called consensus fasta sequences for eight individuals using ANGSD. We used the following commands for each individual: ‘-doFasta 3 -explode 0 -doCounts 1 -setMaxDepth 50 -setMinDepth 8 -minMapQ 20 -minQ 10 -only_proper_pairs 1 -remove_bads 1 -uniqueOnly 0’. We included one individual each from PK10 (Senegal), Ouahigouya (Burkina Faso), Luanda (Angola), Ngoye (Senegal), El Dorado (Argentina), New Orleans (US), Playa Puerto Rico (US), and Cebu City (Philippines). We then subdivided the genome into 10kb windows, calculated the number differences among all pairs, and built neighbor-joining trees using the *ape* package (*20*) in R (*18*). We excluded windows where one or more comparisons in the matrix had less than 2000 sites with data. We tested for monophyly among certain groups using the *is.monophyletic* function in *ape*. To avoid biases from back-to-Africa *Aaa* admixture in proto-*Aaa* populations, we excluded any trees where *Aaa* and proto-*Aaa* were not monophyletic with respect to *Aaf.* We conducted bootstrap analyses by resampling 10kb windows with replacement (1,000 replicates) and repeating the analysis for each bootstrap replicate.

### F3 and F4 admixture inference

We next examined contemporary patterns of long distance vector migration by identifying recent admixture events or secondary contact.

Putatively unadmixed parent populations from each subspecies were selected based on prior studies (*2*). *Aaf* populations included: Franceville, Gabon; Bantata and Kedogou, Senegal. *Aaa* populations included: New Orleans, USA and Ho Chi Minh City, Vietnam.

To preserve admixture signals related to LD while avoiding misleading genetic signals, repetitive regions and those of consistently low or high coverage were removed from the full SNP set followed by removal of SNPs with a minor allele frequency below 0.01 and random subsampling to 1e7 SNPs genome-wide.

Admixture was examined using Patterson’s *F3* applied genome-wide with block jackknifing for each of our sampling sites, using unadmixed *Aaa* and *Aaf* parents. Directionality of the admixture was established with the Patterson’s *F4* test, using Skukusa as an outgroup, due to its basal position in the phylogeny (Figure 1). Both tests were implemented through scikit allel (github.com/cggh/scikit-allel). To avoid false positives from cryptic population structure we repeated all F3 and F4 tests with alternate parent populations.

To distinguish between scenarios of *Aaa*/*Aaf* admixture and *Aaf*/proto-*Aaa* admixture, Patterson’s F4 test was also applied to pure *Aaf* populations with the putative proto-*Aaa* population from El Dorado, Argentina.

### KDR genotyping and haplotype network reconstruction

Initial genotyping on the assembled chromosome of AaegL5 (*3*) did not result in a call for the well characterized F1534C variant. In the likelihood that large and highly structured haplotypes in this region were assembled as alternative haplotypes, we aligned all alternative haplotypes to their most probable analogue in the core genome using minimap2(*27*), allowing up to 1% sequence divergence and alternative matches within 95% of the primary alignment [-ASM10 -p0.95]. Alternative haplotype NW_018735214.1 aligned across the 5’ end of KDR encompassing many of the known resistance loci. Variants called across this alternative haplotype were spliced into the original locus allowing the recovery of known alleles for F1534C, V1016G and V1016I, and others.

Reads were realigned to this reconstructed region using BWA (*4*) and individual read-based phasing was performed using WhatsHap (*28*); the resulting phase blocks were phased across the whole dataset using SHAPEIT4 (*14*). Haplotype networks for this region were reconstructed across all non-synonymous coding SNPs using GeneHapR (*29*).

### Scans for positive selection

*Ae. aegypti* has undergone a series of phenotypic shifts in recent evolutionary history, beginning with the shift from forest-dwelling host generalist phenotypes to preferring human hosts and living around human settlements, and later further adapting to modern human landscapes and interventions such as insecticides (*30*–*32*). Rose et al. (*2*) identified several genomic regions that are highly differentiated between generalist and human-preferring populations in West Africa. To discover genomic regions targeted by natural selection during the more recent process of human specialization and adaptation to modern landscapes, we scanned the genome for regions that are highly differentiated in North American populations (Miami, New Orleans, and Mexico) when compared to a proto-*Aaa* population (El Dorado), after polarizing with a West African *Aaf* population (PK10) to isolate signals specific to North American populations. North American populations were chosen as they likely represent the closest proxy to early New World populations introduced during the AST, since most of the original founding South American populations were eradicated in the 1950s (*33*).

We scanned the genome for highly differentiated regions using the Population Branch Statistic (*34*), which is a polarized measure of genetic differentiation intended to localize differentiation to one branch on a three-branch population tree. To minimize any impact of admixture on signals of differentiation, we used admixture-corrected allele frequencies obtained using NGSadmix (*15*). We ran NGSadmix on the full set of genotype likelihoods at SNPs described above with the minimum number of individuals with data (-minInd) set to 500, no minimum allele frequency filter (n = 67,182,646 SNPs after filtering), and the Q matrix with admixture proportions at K=8 obtained from genetic structure analysis described above. By supplying an existing Q matrix, NGSadmix assumes admixture proportions for each individual according to this matrix, and then estimates allele frequencies at each SNP under the assumed admixture model. We calculated PBS at SNPs with a minor allele frequency of at least 0.05 in the target population using sample sizes of 155, 18, and 231 for the target, contrast, and outgroup populations, respectively. We counted the number of individuals in each of the admixture components with admixture proportions of at least 0.5 to define sample sizes used in PBS calculations. We calculated PBS for individual SNPs as well as in 20-SNP windows. After removing windows greater than 10,000 basepairs in size, we ranked windows based on PBS scores, then ranked the top 1,000 windows based on the number of individual SNPs in the top 0.01% of the SNP-wise distribution genome-wide. We obtained gene names and exon boundaries from the AaegL5 AGWG GFF hosted on Vectorbase.org (version 66, VectorBase-66_AaegyptiLVP_AGWG.gff). We plotted individual window values as well as the rolling mean of 30 windows.

The strongest window of differentiation shared among several North American populations (Population Branch Statistic (PBS) = 0.5713) was located on the q arm of chromosome two and centered at 401.75 MB (Figure S29). The window of increased differentiation is approximately two megabases wide and centered over a region with relatively few annotated genes (Figure S29). The gene, AAEL025736, at the center of the peak, is comprised of 18 exons spanning ∼477 kb and is annotated as a Toll-like receptor Tollo, a transmembrane receptor involved in antimicrobial and antiviral immune responses (*35*). The second strongest window of differentiation is located on chromosome one centered at 79.75 MB (Figure S30). The width of the peak is somewhat smaller compared to the first peak, but it covers a more gene dense area (Figure S30). The gene at the center of the peak, AAEL017005, is the RYamide receptor that pairs with RYamide neuropeptides and is thought to play a role in regulating feeding and digestion (*36*). Additional regions of differentiation were identified and are listed in Table S2.

### Neo-sex chromosome evolution

Unlike well known insect species such as *Drosophila melanogaster* and *Anopheles gambiae* who have fully-formed sex chromosomes, *Aedes aegypti* has three autosomes with the male determining sex locus located near the centromere of chromosome one (*37*–*40*). Suppressed recombination and male-female differentiation around the sex determining locus are key early steps in the formation and growth of an incipient X and Y chromosome system (*41*). Fully evolved X and Y sex chromosome systems usually involve a degenerate Y chromosome with few genes, and the evolutionary processes leading to such degeneration is highly dependent on effective population size (*41*). Taking advantage of the large sample sizes and broad sampling in the Aaeg1200 dataset, we asked whether the extensive genetic drift and history of population bottlenecks in *Aaa* has altered male-female differentiation around the sex locus relative to *Aaf*.

To identify differences in allele frequencies between males and females, we randomly selected 150 males and females from *Aaa* and *Aaf* populations separately and calculated minor allele frequencies using ANGSD (-minMapQ 20 -minQ 10 -remove_bads 1 -uniqueOnly 0 -minInd 120 -capDepth 45 -doSaf 1 -domaf 2 -GL 1 -doMajorMinor 5). We limited the analysis to robust sites described above and polarized major and minor alleles to the reference sequence. We further limited analysis to only sites where both male and female frequencies were calculated from at least 135 individuals and the sum of the minor allele frequencies exceeded 0.01 to remove invariant sites. To identify sites likely to be found on the male-specific haplotype carrying nix, we filtered based on male frequencies, requiring male minor allele frequency to be greater than 0.48 and less than 0.52. We plotted female minor allele frequencies at individual SNPs, as well as frequencies averaged within non-overlapping one megabase windows.

We estimated high-resolution allele frequencies from 150 males and 150 females and female allele frequencies at candidate male-only SNPs (male frequency 0.5 +/- 0.02) along chr1 for *Aaf* and *Aaa*. In both subgroups, female frequencies were well correlated with candidate male-only SNPs, as expected under normal combinatorial exchange among the sexes (Figure S31). In *Aaf*, male-female differentiation is highest with fixed female alleles in the region directly surrounding the male-determining gene *Nix,* but the footprint of differentiation extends into the q-arm of the chromosome, over a total of approximately 67 Mb (MB 128-205, Figure S31). The general pattern and position of differentiation are similar in *Aaa*, but highly differentiated and fixed female allele frequencies are found over a much broader area extending to ∼MB 205, with moderate differentiation extending another 5 MB into the q arm (MB 128 - 210, Figure S31). Our coalescent analysis above provides evidence for a history of reduced effective population sizes in *Aaa* relative to *Aaf*, and here we show that the degree of male-female differentiation has increased in *Aaa*, especially at the boundary of the footprint, suggesting that incipient Y chromosome evolution may be accelerated by strong genetic drift in *Aaa* relative to *Aaf*.

### Chromosomal inversions

Large genomic structural rearrangements, such as chromosomal inversions, change the recombination and regulatory context for genes captured in the rearrangement. Chromosomal inversions, in particular, can play a role in speciation and local adaptation, among other evolutionary processes (*42*). Previous studies suggest the presence of large structural variants in *Ae. aegypti (43–47)*, generating a large number of putative chromosomal inversion calls. While some of these studies have seemingly detected the same variants (*45*, *46*), there has been little insight into the age, importance, or global distribution of these inversions, and none of these studies have developed molecular markers for the inversion itself, which would enable a wider and more detailed survey to be carried out. The availability of large scale genome re-sequencing data for *Ae. aegypti* now allows us to investigate inversion polymorphisms by searching for the demographic signal of increased genetic linkage and divergence of the two inversion haplotypes, rather than for the positional rearrangement, and to use these genetic signals to derive informative markers for these inversions and facilitate future studies.

We first removed repetitive regions and those of consistently low or high coverage from the full SNP set followed by removal of SNPs with a minor allele frequency below 0.01 and random subsampling down to 1e7 SNPs genome-wide.

We developed a pipeline to identify and validate putative inversion calls directly from the demographic data using Lostruct (*48*) (Figure S32). Lostruct (*48*) performs windowed PCA followed by similarity scoring to identify linked blocks of population structure along the genome, and has previously been employed to scan for novel chromosomal inversions (*49*). Lostruct was applied to all countries with panmictic populations and more than 20 samples in the Aaeg1200g dataset. Similarity matrices were calculated between all 0.5 MB non-overlapping windows along the chromosome and demographic blocks were identified by hierarchical clustering using pvclust [ P < 0.01 after 1000 bootstraps with multiscale resampling (au)] (*50*). Candidates larger than 150 MB or smaller than 2.5 MB were removed.

To distinguish putative inversions from other regions of suppressed recombination, we re-ran the PCA for each candidate region, k-means clustered (n=3), and tested for both variability within [ sum squares < 5] and between clusters [sum squares > 25] as well as the centrality of the putative homokaryotype cluster [ < 10% deviation from the midpoint of the two homokaryotype clusters] as performed previously (*51*, *52*). To allow for inversions which do not dominate the PC1 signal, we retested after rotation of the PC1/2 positions around the origin, preferring those with lower angles of rotation. Inversion candidates for which the heterokaryotype samples did not exhibit either increased heterozygosity or significant F3 (*53*) were removed. Within countries, inversions that overlapped positionally by 0.5 MB or more and exhibited identical k-means cluster assignments were merged.

Inversion SNP markers were identified within each country by association with the k-means clusters (chi-square, P<=1e-7). Between countries, inversions were merged where these markers were in LD with each other (r^2^ > 0.25). The inversion haplotype that was predominantly ‘ref’ genotypes was arbitrarily defined as the uninverted form; the inversion karyotype was estimated for each sample as the median genotype of all marker SNPs. Tests for Hardy-Weinberg equilibrium were applied at both the population and country levels.

We used a PCA-based test to scan the Aaeg1200 panel at the country level and identified a total of 10 inversions ranging in size from 2.5 MB - 44.5 MB, covering close to 10% of the genome (Figure S33). The inversions predominated on chromosome 1, where 7 inversions represented 28% of the chromosome length with numerous proximal or overlapping inversions. A subset of the inversions found on chromosomes 1 and 2 are positionally similar to inversions identified in a recent study based on Hi-C data (*47*), although we found evidence for inversions not identified in that study. Interestingly, we found no robust evidence for inversions on chromosome 3 in contrast to the Hi-C study that found seven inversions on that chromosome, further underscoring differing sample sets and sensitivities among the two studies. Inversion frequencies ranged from below 1% to greater than 50% (Table S3) with all inversions at moderate (>5%) frequencies found in both *Aaa* and *Aaf* subspecies. While this study would not be powered to find inversions that are entirely fixed between *Aaa* and *Aaf* it is notable that, as in Redmond *et al.* (*46*) and Liang et al. (*47*), no inversion was found close to fixation between subspecies.

## Supplementary Figures

**Figure S1:**
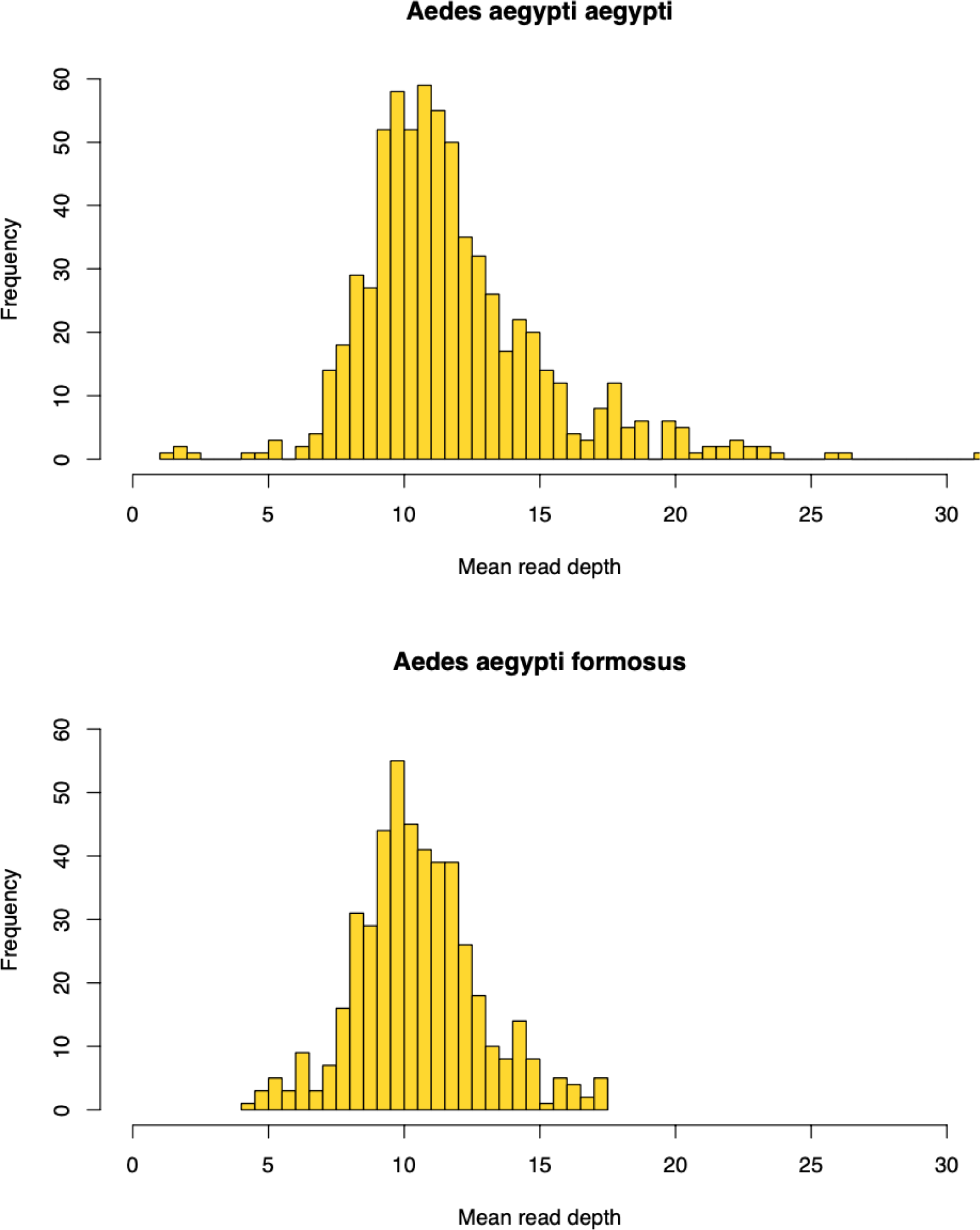
Mean read depths. Histogram plots summarizing mean read depth for *Aaa* (top) and *Aaf* (bottom) individuals. Read depth was calculated using ANGSD (see methods) and averaged for each individual. Supplementary Table X provides mean read depth and read depth distributions for each individual.

**Figure S2:**
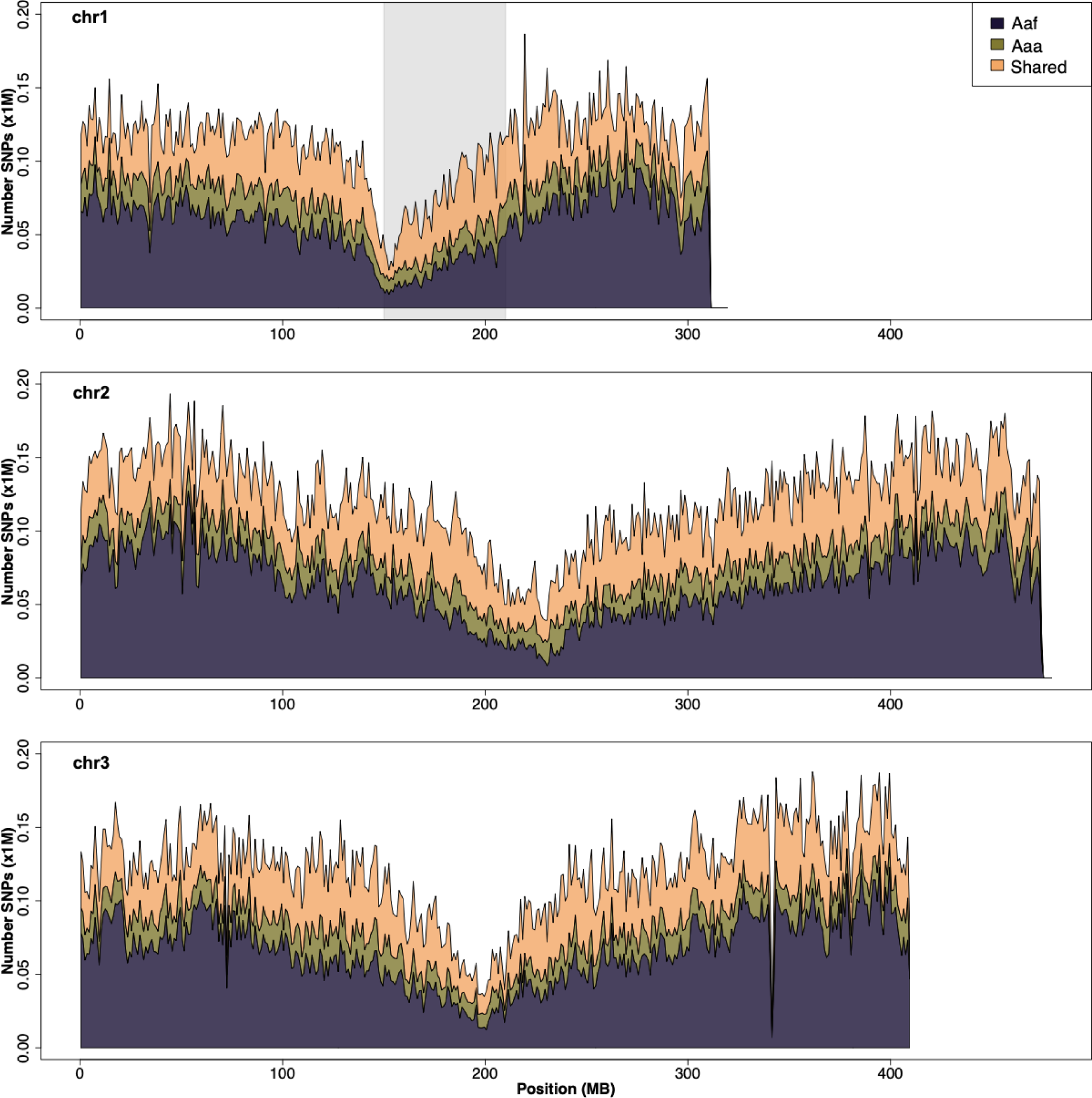
Number of SNPs shared and private among subspecies. Stacked sum plot of the number of SNPs per one megabase block along each chromosome (1 on top, 2 middle, 3 on bottom) with each color showing the number of SNPs that are either private to non-African (*Aaa*) populations, African (*Aaf*) populations, or shared and found to be segregating in populations both subspecies. The gray shaded area shows the highly differentiated region between males and females.

**Figure S3:**
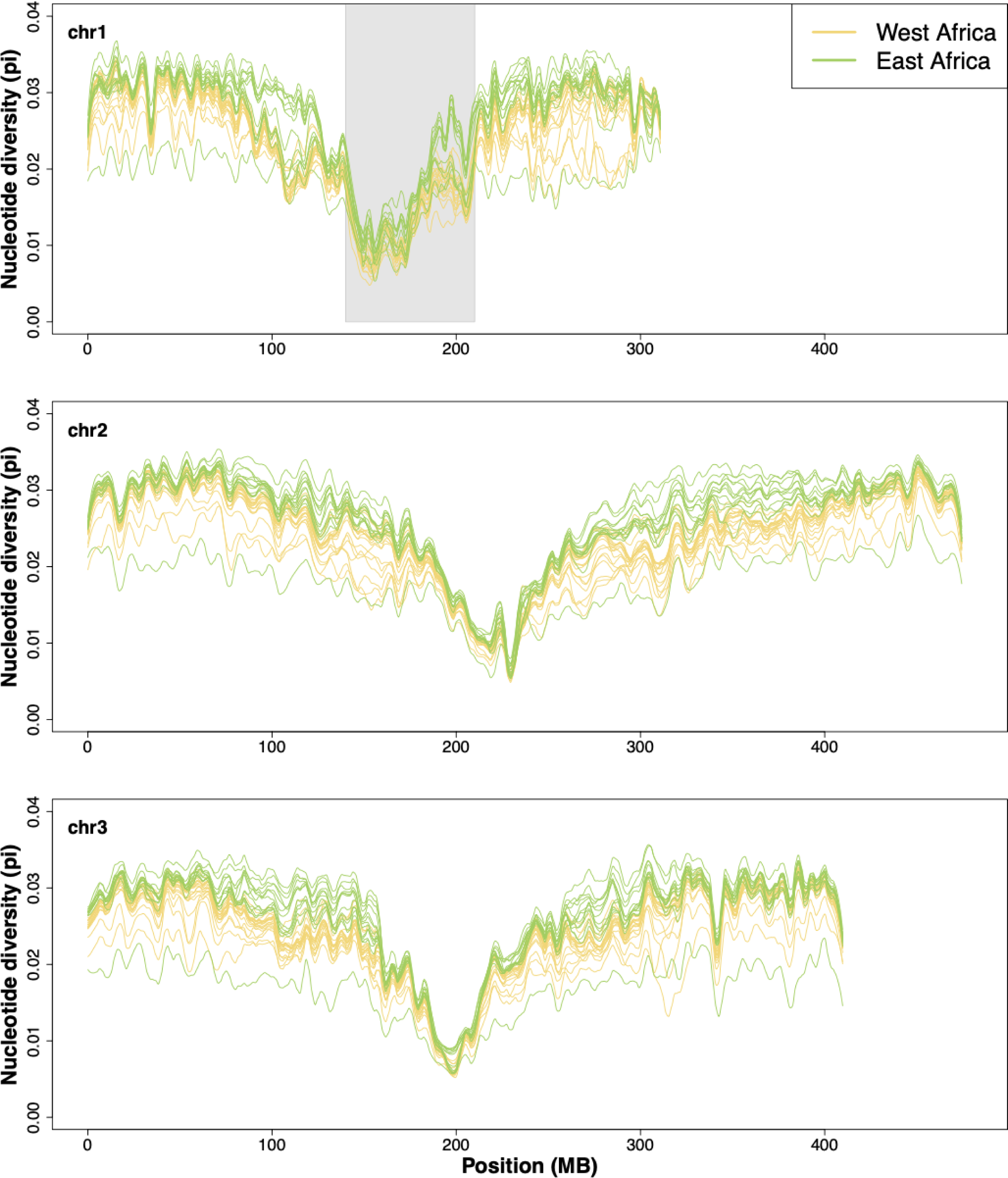
Nucleotide diversity variation along chromosomes in *Aaf*. Nucleotide diversity variation along chromosomes in Aaf for chr1 (top), chromosome 2 (middle), and chromosome 3 (bottom). Nucleotide diversity (*π*) was calculated in 1MB windows using ANGSD and summarized using the loess.smooth function in R. Colors indicate the region of origin according to the legend. The gray bar indicates the sex-differentiated region.

**Figure S4:**
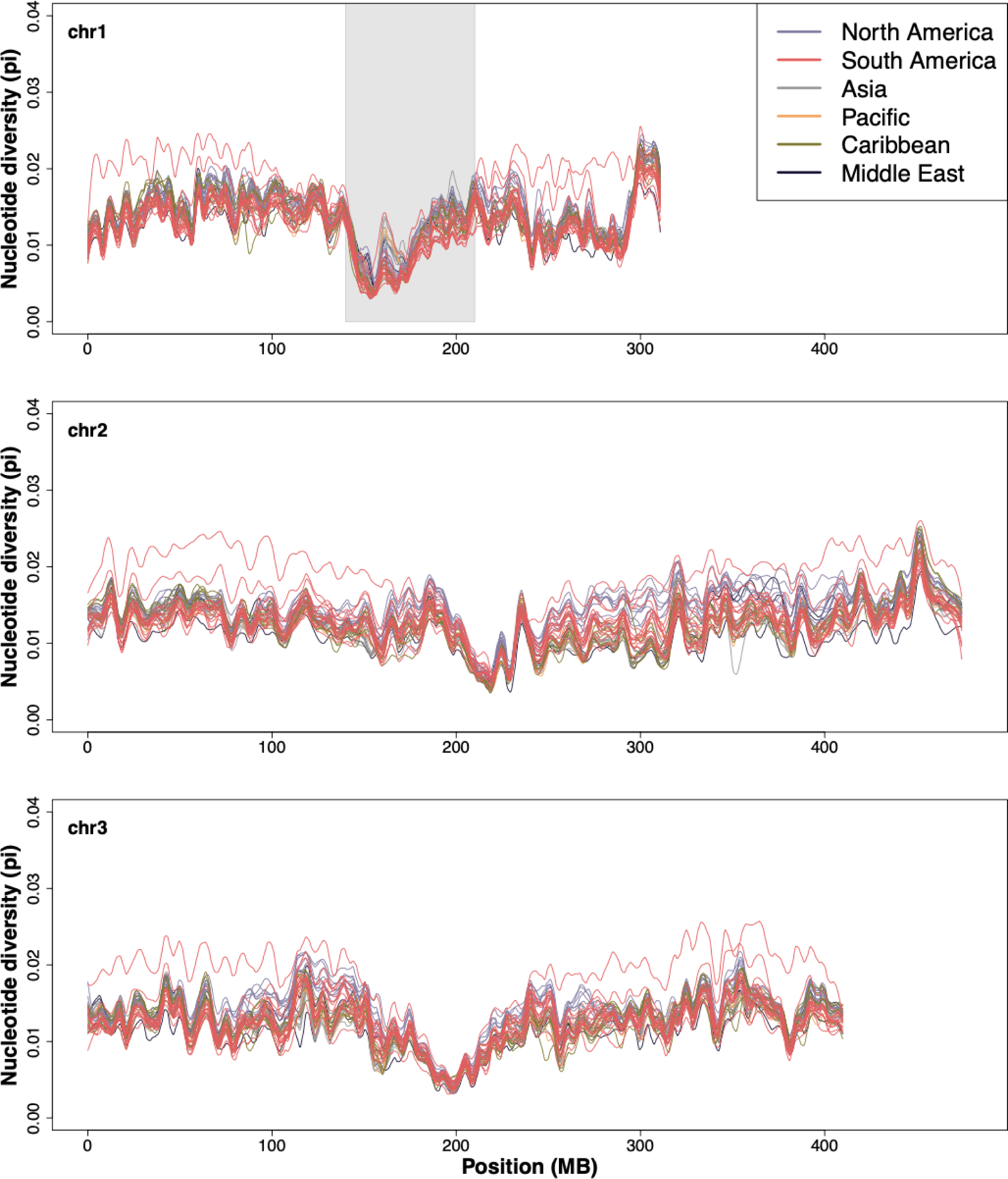
Nucleotide diversity variation along chromosomes in *Aaa*. Nucleotide diversity variation along chromosomes in Aaa for chr1 (top), chromosome 2 (middle), and chromosome 3 (bottom). Nucleotide diversity (*π*) was calculated in 1MB windows using ANGSD and summarized using the loess.smooth function in R. Colors indicate the region of origin according to the legend. The gray bar indicates the sex-differentiated region.

**Figure S5:**
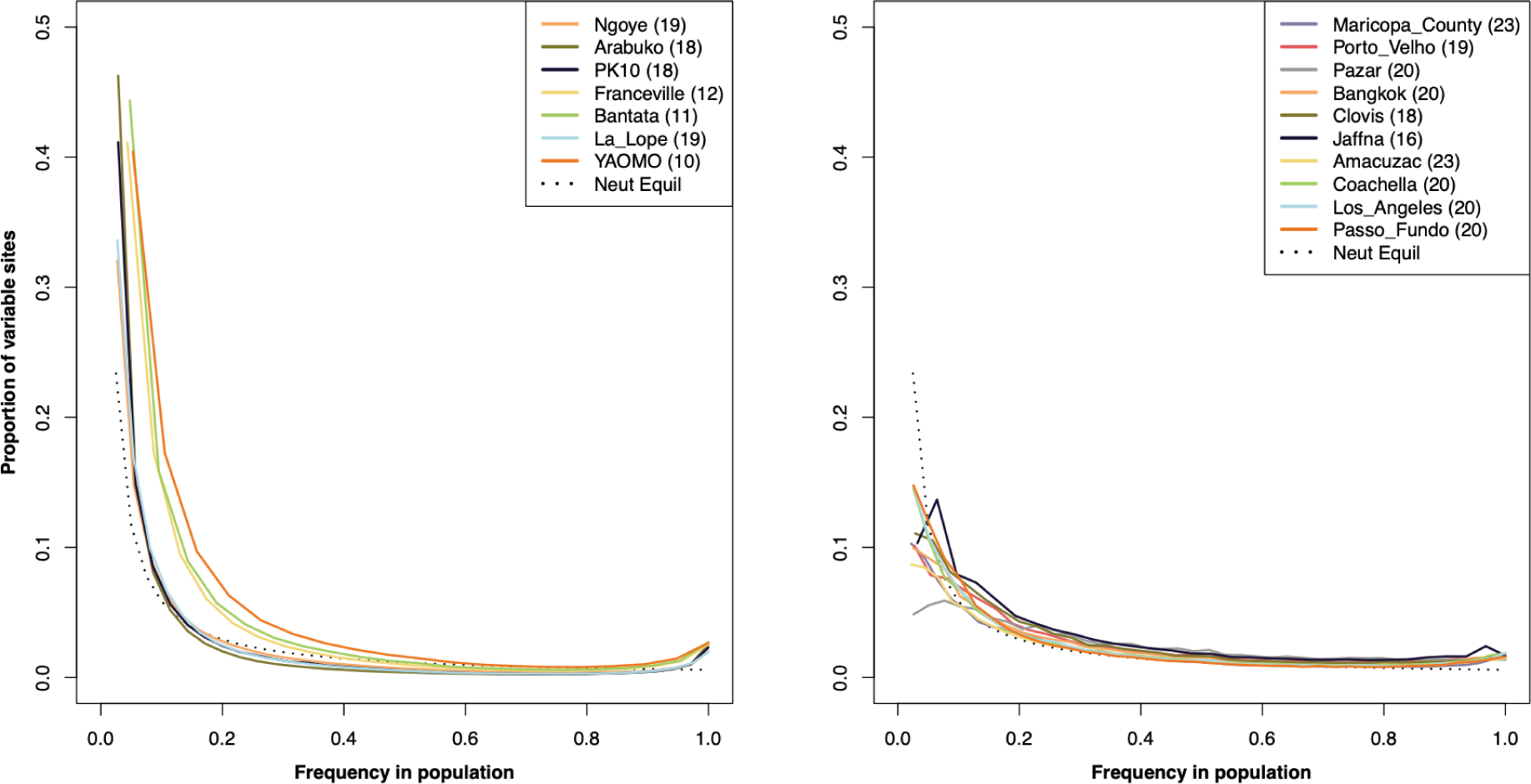
Frequency of derived mutations in populations. Site frequency spectra for representative populations of *Aaf* (left panel) and *Aaa* (right panel). Spectra were calculated using realSFS from robust sites across all three chromosomes (see Methods). Variant frequencies were converted from counts to proportions to facilitate comparison among populations. The dotted line in each panel shows expected site frequency spectrum under a neutral equilibrium model.

**Figure S6:**
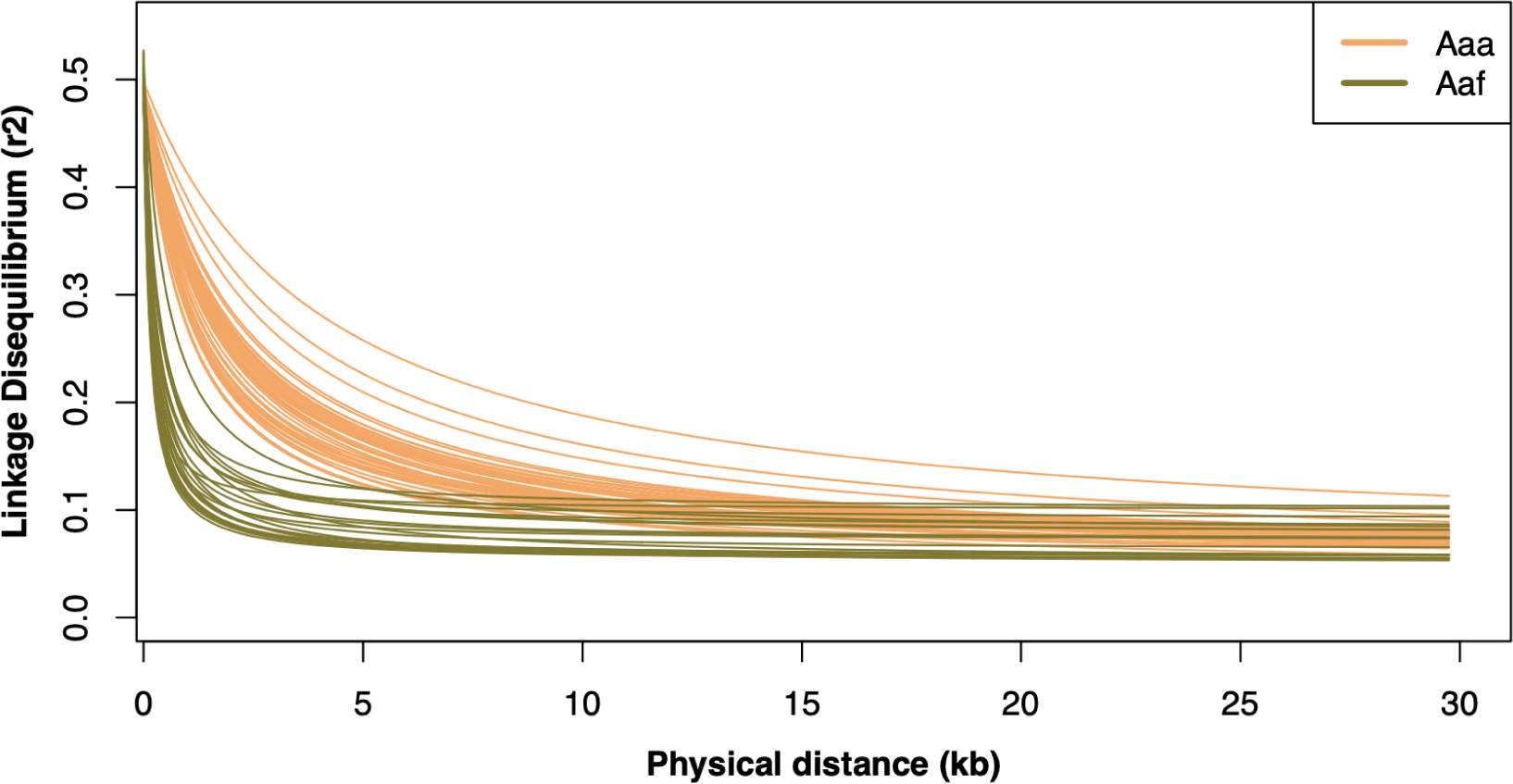
Decay of linkage disequilibrium as function of physical distance. Linkage disequilibrium decay curves shown as a function of physical distance among SNPs. The linkage of disequilibrium estimator, *r*^2^, was calculated using ngsLD using SNPs in 1 megabase windows from the center of each chromosomal arm on all three chromosomes. Only populations with at least 10 individuals are shown.

**Figure S7:**
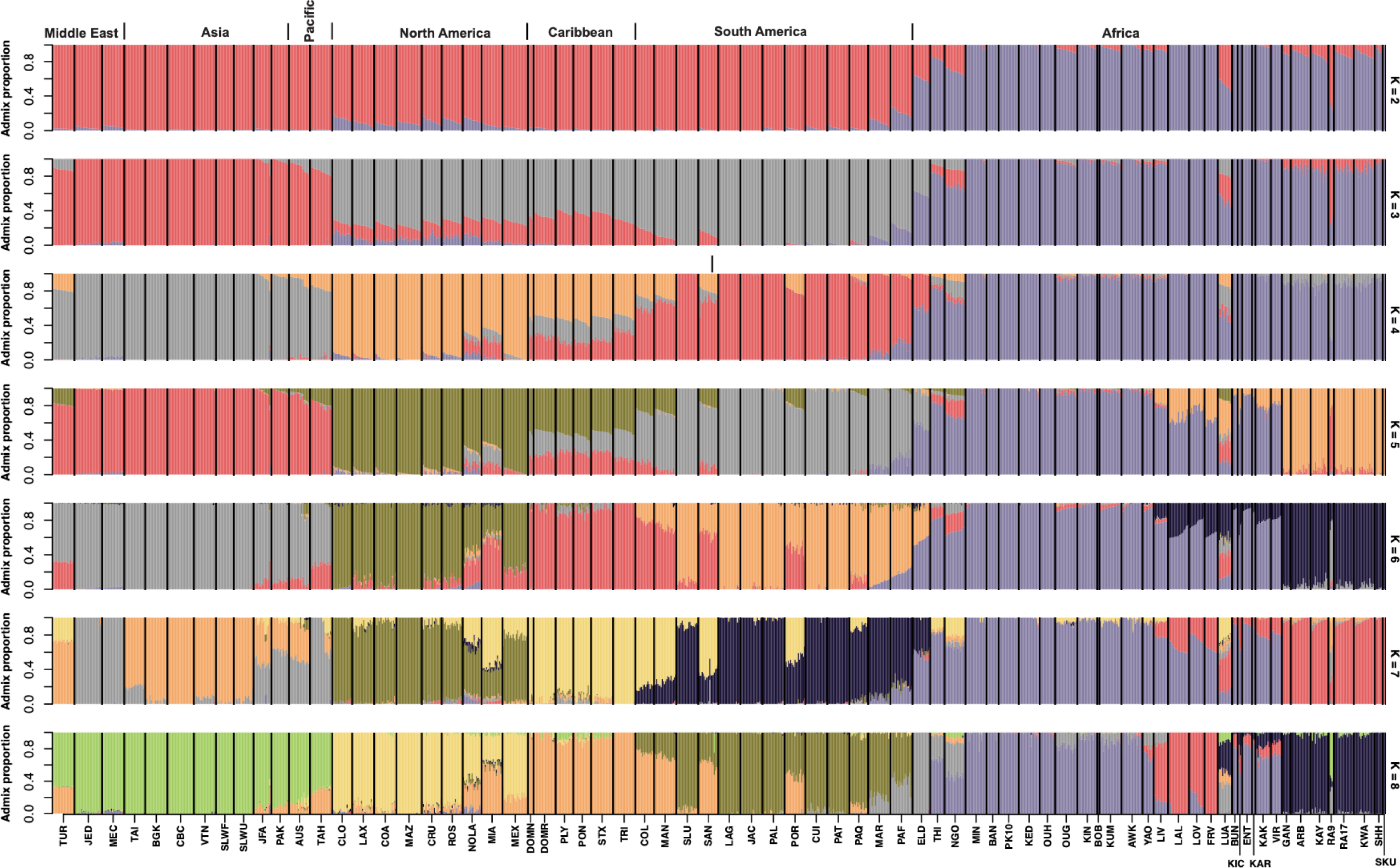
Admixture analysis with 2-8 ancestral populations. Admixture analysis of all 1,206 individuals using NGSadmix. Each row shows results assuming different numbers of ancestral populations ranging from K=2 (top) to K=8 (bottom). Each bar represents an individual genome with colors indicating ancestry proportions assigned to each ancestral population. Populations are labeled along the bottom and full names are in Table S4.

**Figure S8:**
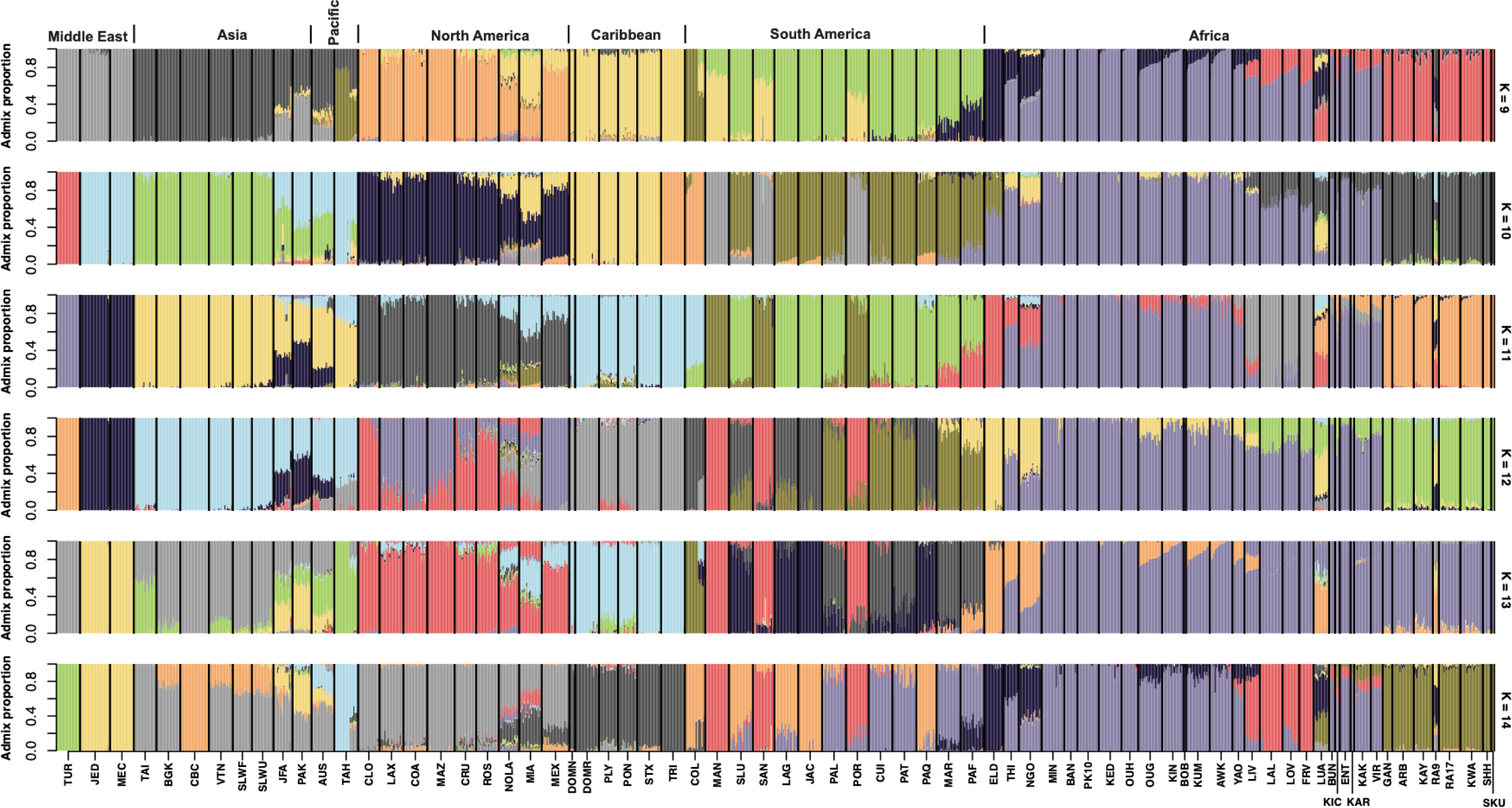
Admixture analysis with 9-14 ancestral populations. Admixture analysis of all populations using NGSadmix. Each row shows results assuming different numbers of ancestral populations ranging from K=9 (top) to K=14 (bottom). Each bar represents an individual genome with colors indicating ancestry proportions assigned to each ancestral population. Populations are labeled along the bottom and full names are in Table S4.

**Figure S9:**
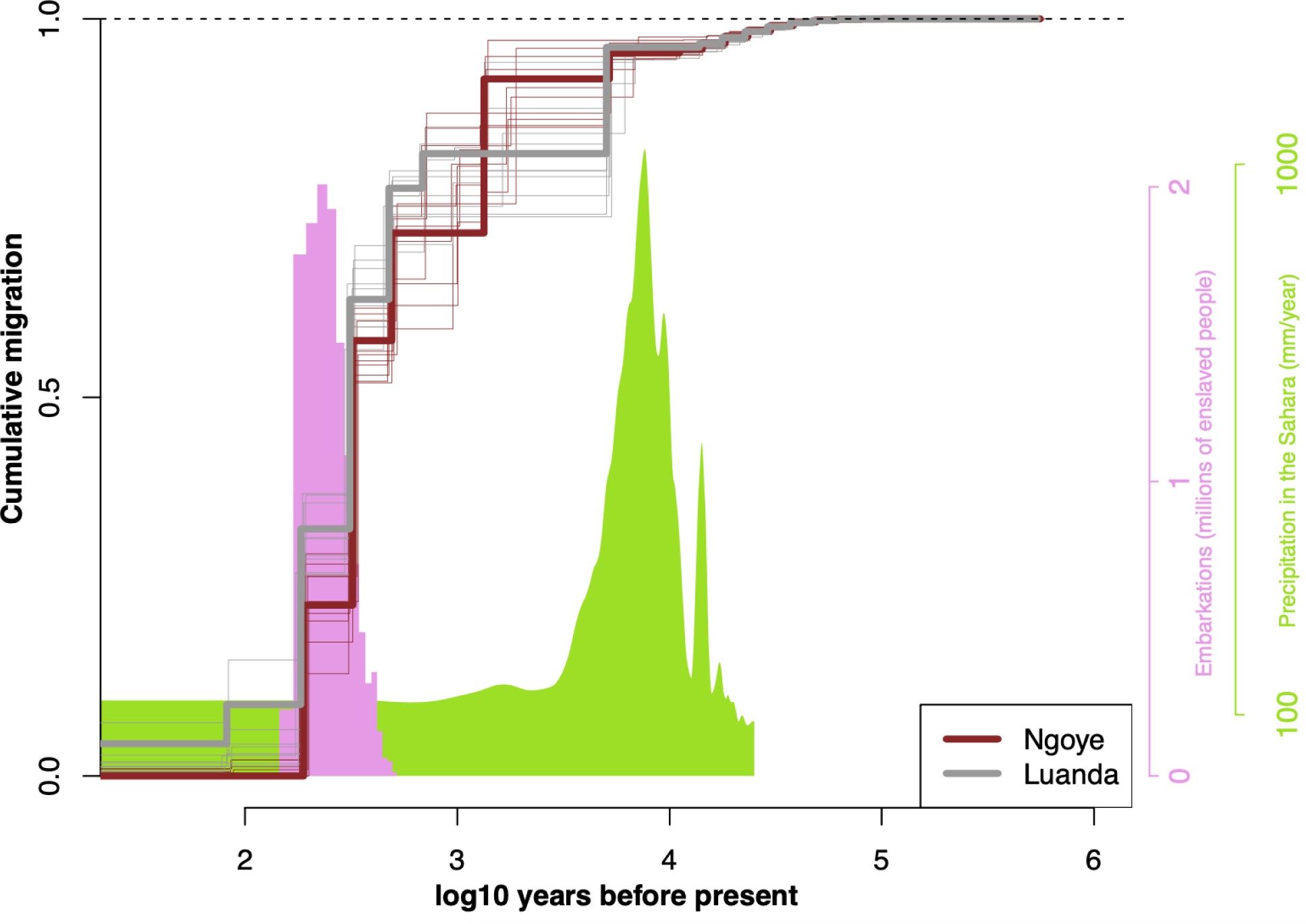
Cumulative migration between El Dorado and African proto-*Aaa* populations. Cross-coalescent analysis using MSMC-IM of El Dorado, Argentina with Luanda, Angola and Ngoye, Senegal. Cumulative migration plotted as a function of time on log10 scale (assuming 12 generations per year) with bootstrap replicate curves shown in the same colors as the empirical data but with a smaller line weight. Cumulative migration is expected to plateau at one going back in time. The pink histogram shows the estimated number of slave vessel embarcations scaled by millions of enslaved people (Slave Voyages 2023) on the alternative, pink y-axis at right. The green histogram shows precipitation levels in the Sahara (mm/year; Beserra et al. 2006) according to the green scale at right.

**Figure S10:**
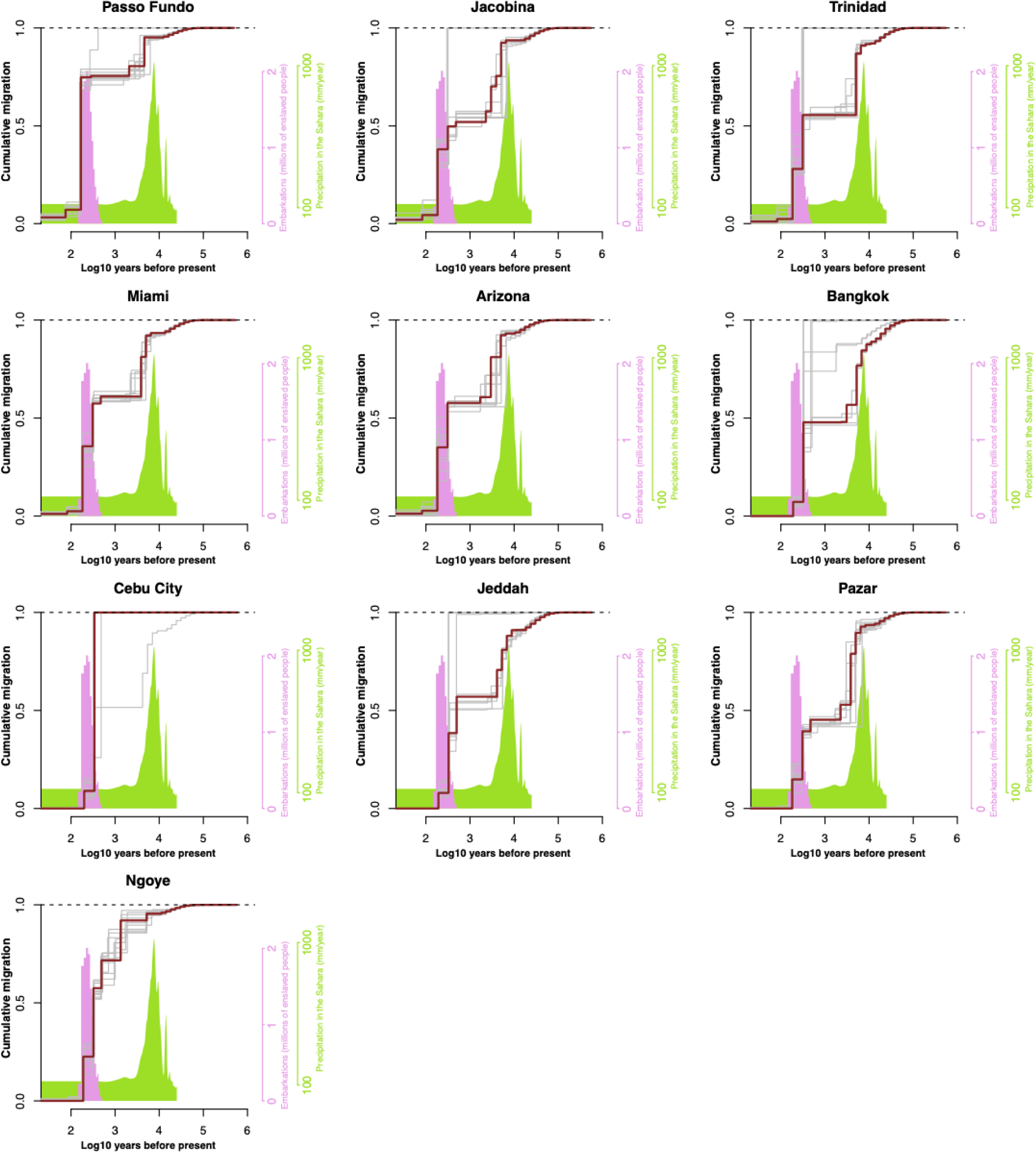
Cumulative with El Dorado estimated using MSMC-IM with bootstraps. Cross-coalescent analysis using MSMC-IM of El Dorado, Argentina and representative populations as in main text Figure 2. Cumulative migration plotted as a function of time on log10 scale (assuming 12 generations per year) with bootstrap replicate curves shown in gray. Cumulative migration is expected to plateau at one going back in time. The pink and green histograms are the same as for Figure S9.

**Figure S11:**
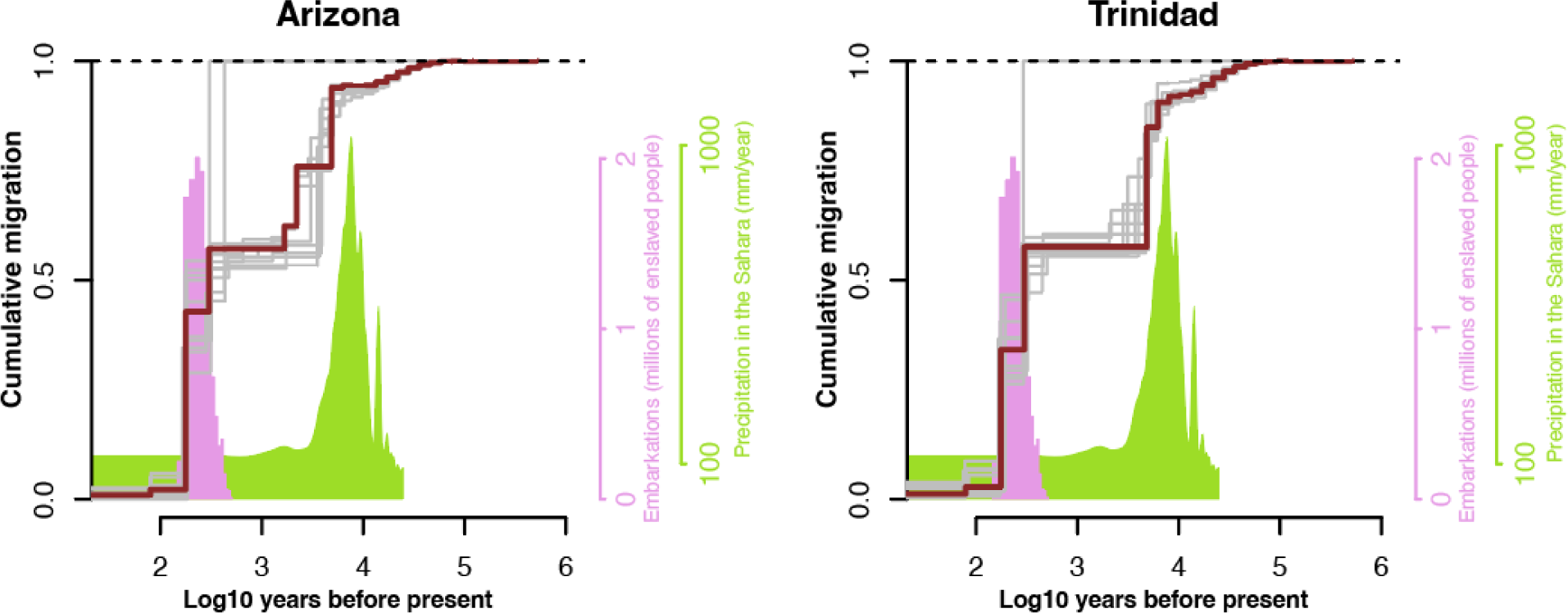
Cumulative with Luanda estimated using MSMC-IM with bootstraps. Cross-coalescent analysis using MSMC-IM of Luanda, Angola and *Aaa* populations as in main text Figure 2. Cumulative migration plotted as a function of time on log10 scale (assuming 12 generations per year) with bootstrap replicate curves shown in gray. Cumulative migration is expected to plateau at one going back in time. The pink and green histograms are the same as for Figure S9.

**Figure S12:**
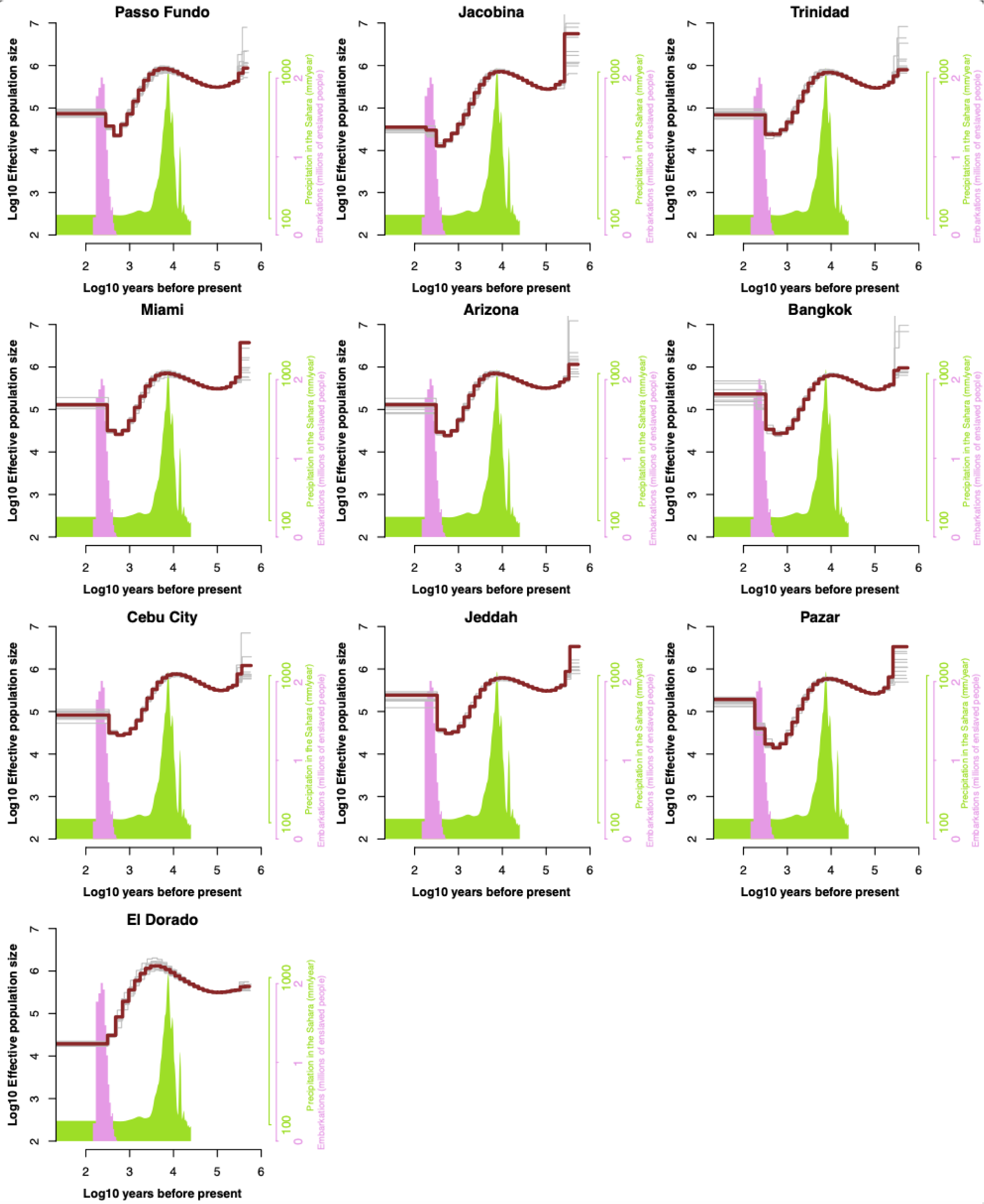
Effective population size estimated using MSMC2 with bootstraps. Effective population size curves on Log10 scale as a function of time also on Log10 scale with bootstrap curves shown in gray. The populations are the same as for main text Figure 2. The pink and green histograms are the same as for Figure S9.

**Figure S13:**
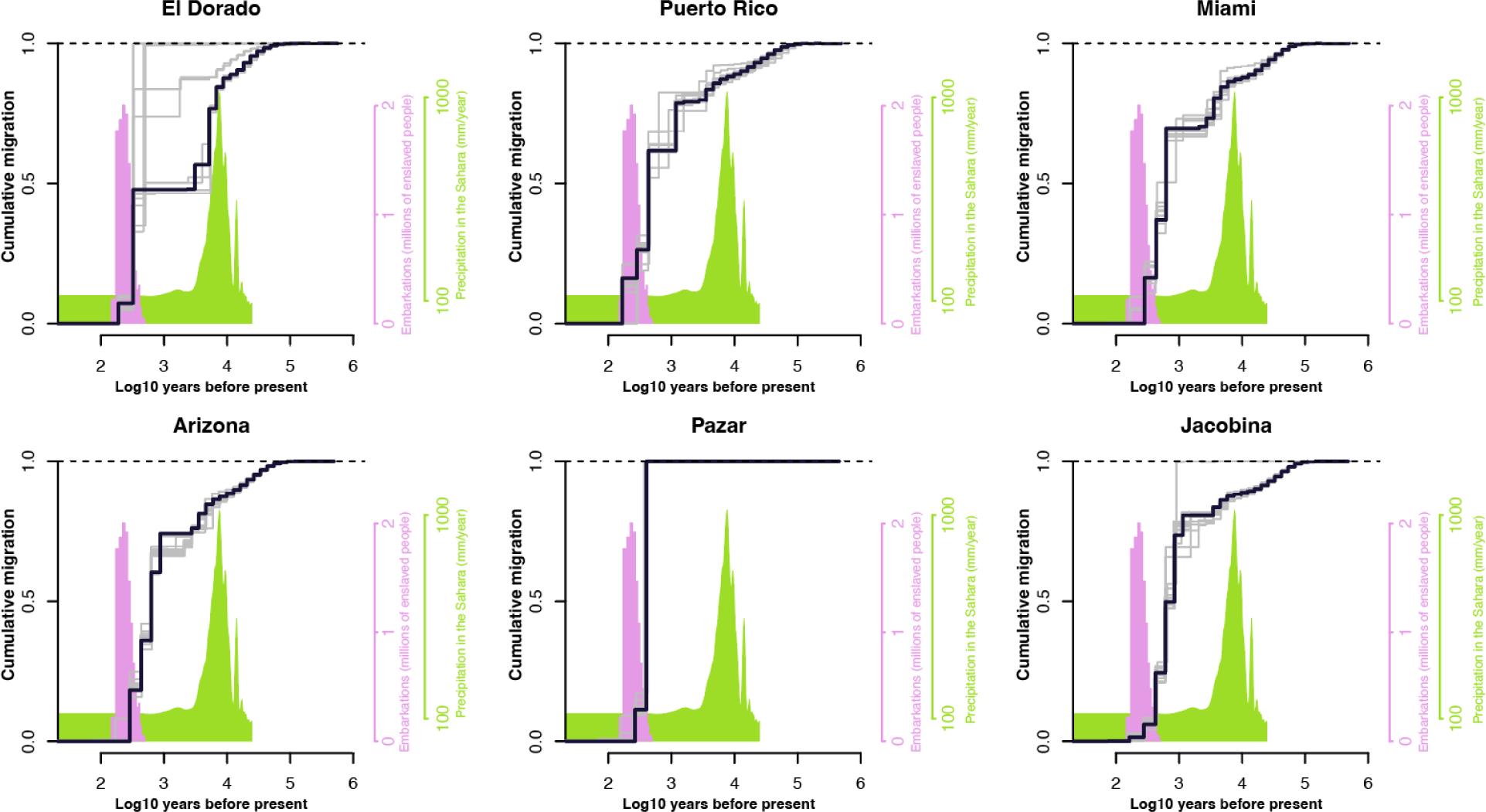
Cumulative with Bangkok estimated using MSMC-IM with bootstraps. Cross-coalescent analysis using MSMC-IM of Bangkok, Thailand and representative proto-*Aaa* or *Aaa* populations as in main text Figure 2. Cumulative migration plotted as a function of time on log10 scale (assuming 12 generations per year) with bootstrap replicate curves shown in gray. Cumulative migration is expected to plateau at one going back in time. The pink and green histograms are the same as for Figure S9.

**Figure S14:**
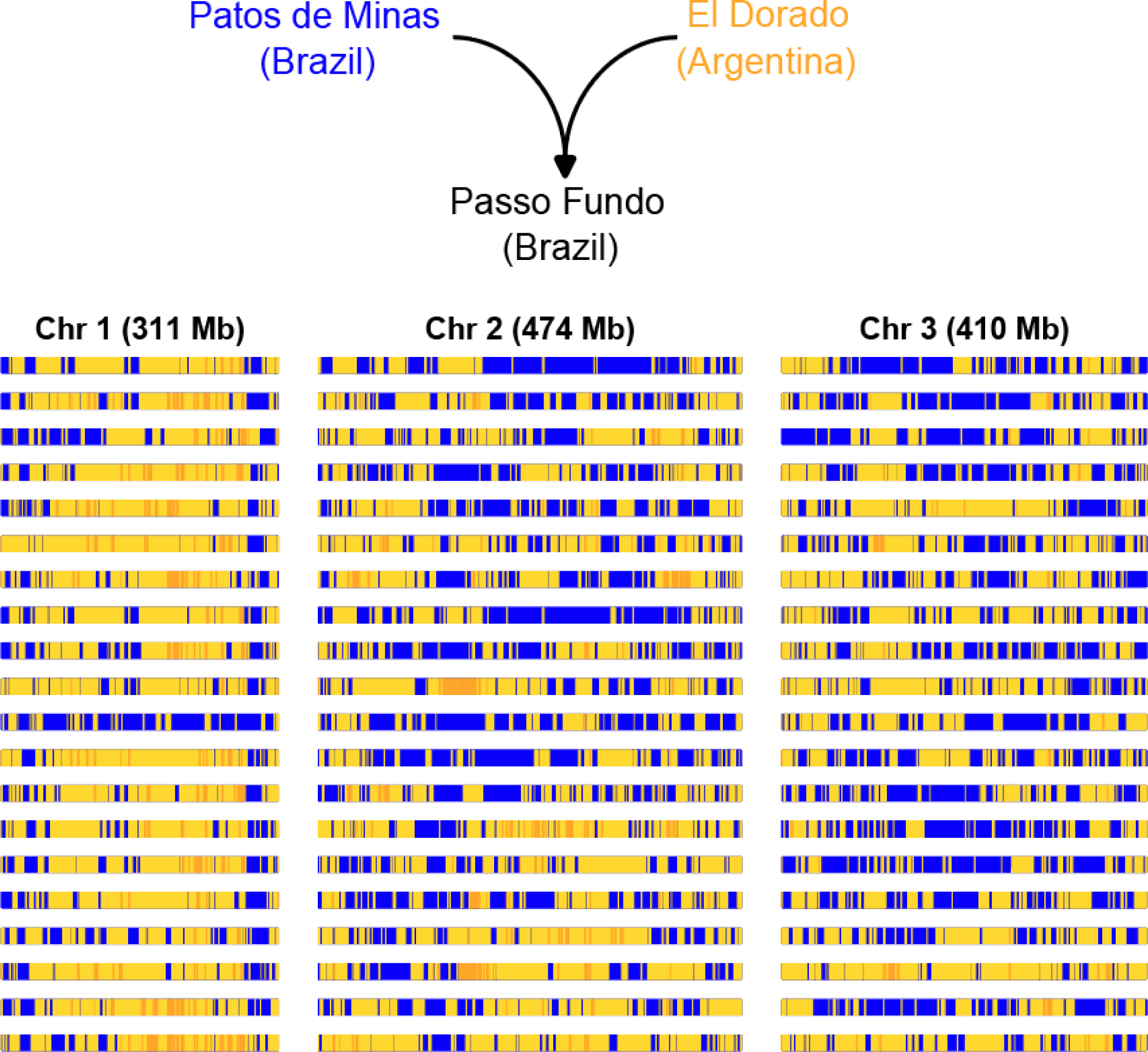
Admixture tracts from secondary contact in Passo Fundo. Admixture tract analysis of Passo Fundo population from southern Brazil using El Dorado as proto-*Aaa* proxy parental population and Patos de Minas as *Aaa* proxy parental population. Ancestry tracts were identified using AncestryHMM (REF, see methods). Each row represents one diploid genome from Passo Fundo where blue represents chromosomal segments homozygous for *Aaa* ancestry, gold represents chromosomal segments homozygous for proto-*Aaa* ancestry, and light yellow represents heterozygous regions.

**Figure S15:**
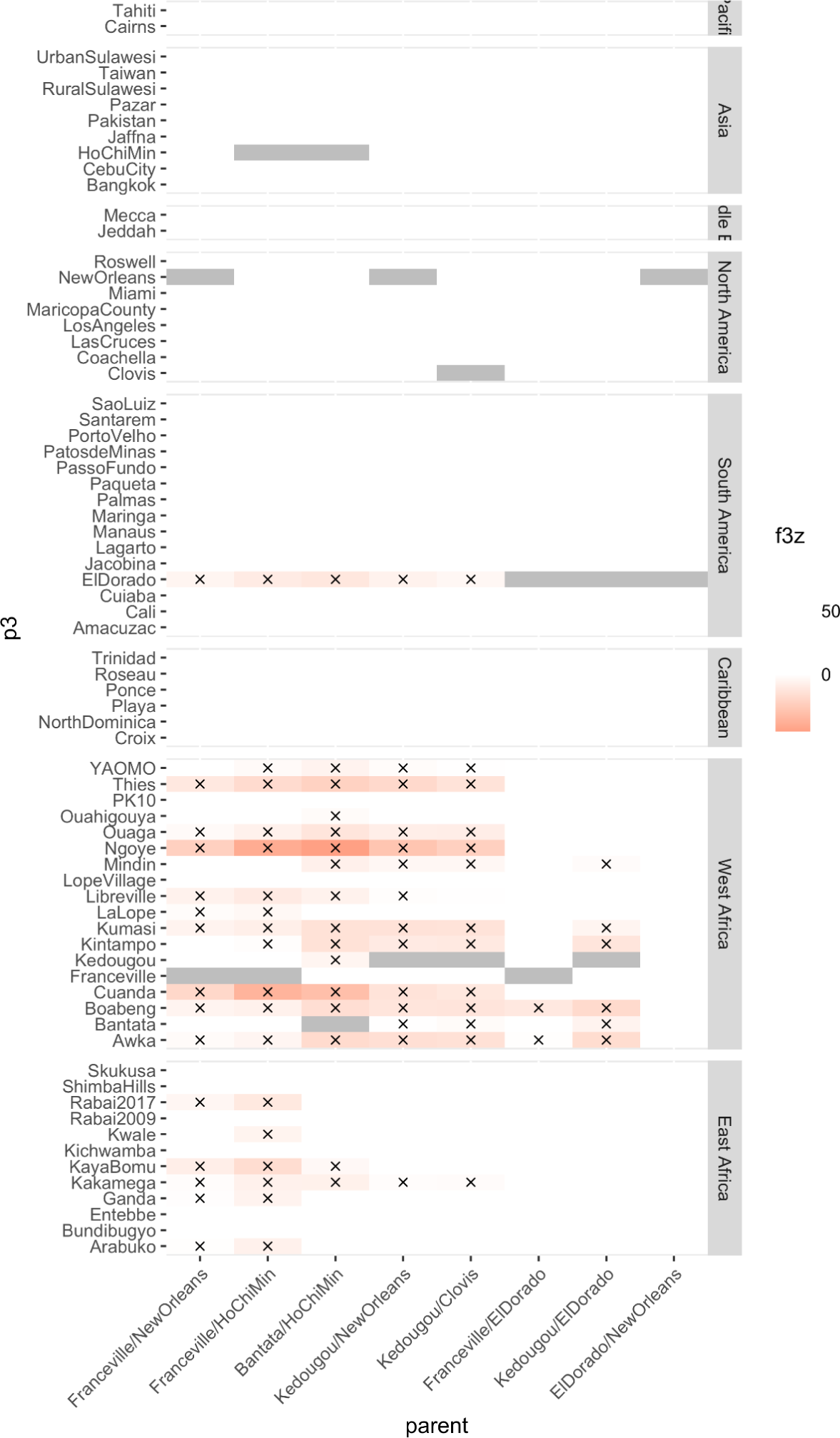
F3 tests for admixture with alternative parental populations. F3 admixture test results for populations by row with alternative parental donors in each column. Populations separated by region as labeled on the right. Colors indicate Z score for the F3 test according to scale on the right with red indicating negative values and therefore stronger evidence for admixture. X marks indicate significantly negative test results (F3 Z-score < -0.3) and gray boxes indicate skipped tests since target population matched one parental.

**Figure S16:**
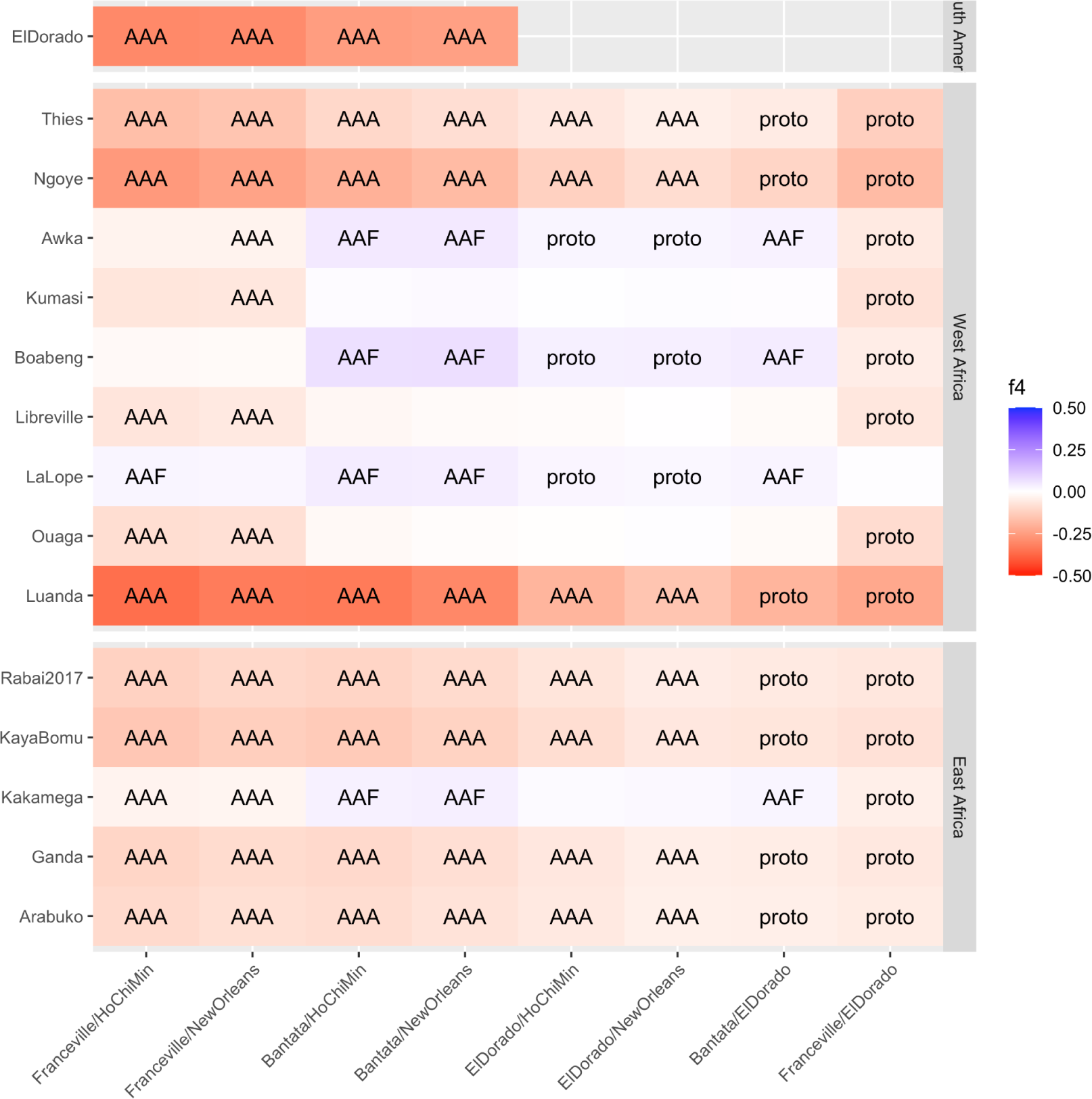
F4 test for admixture with alternative parentals. F4 admixture test results for recipient populations by row with alternative parental donors in each column. The *Aaf* population from Skukusa (South Africa) was used as the outgroup. Populations separated by region as labeled on the right. Colors indicate Z score for the F4 test according to scale on the right with stronger colors indicating stronger evidence for admixture, and color indicating direction of admixture as indicated with labels on each cell.

**Figure S17:**
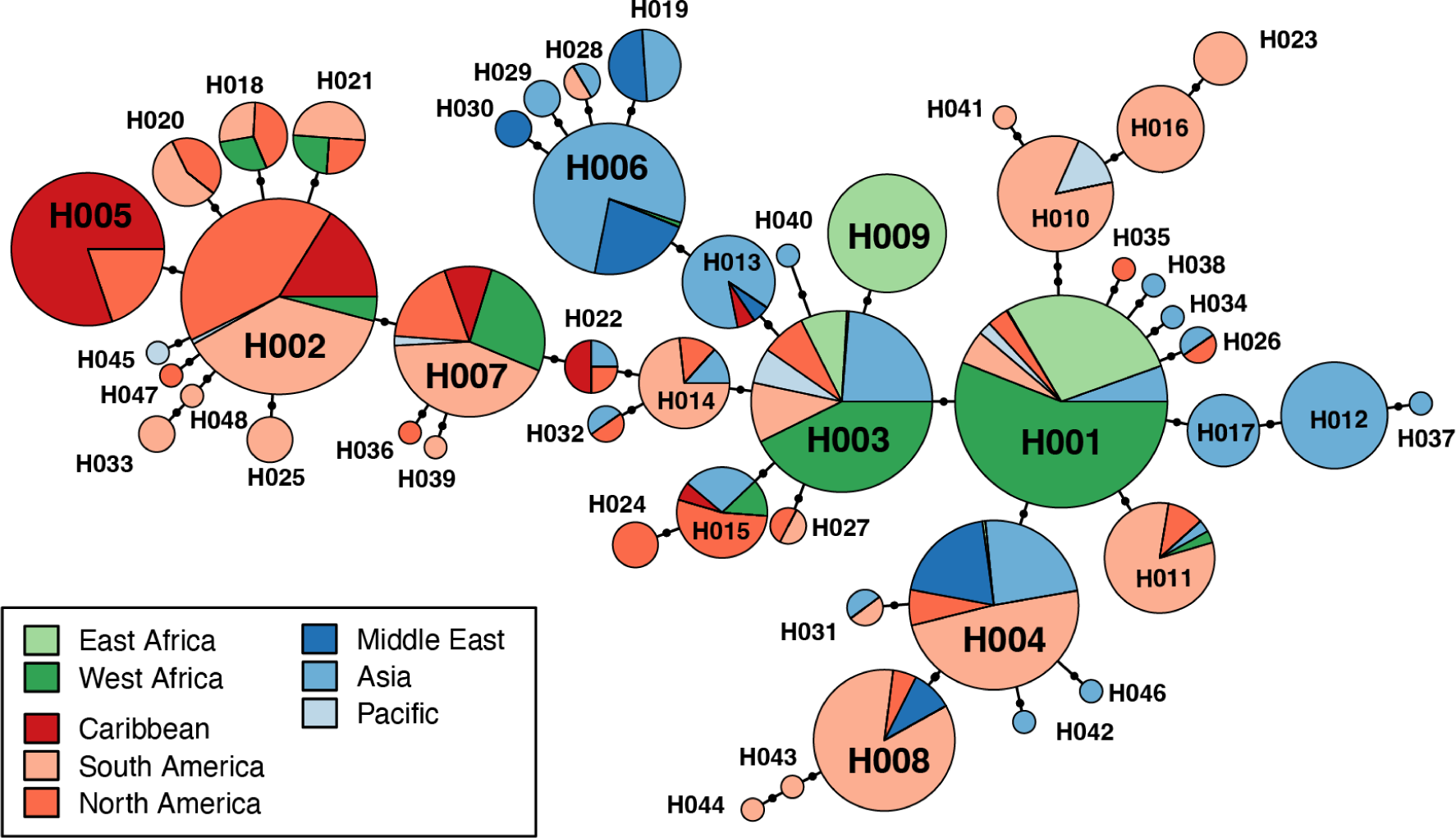
Haplotype network of Knock-Down Resistance mutations. Haplotype network calculated from a set of known Knock-Down Resistance mutations and other non-synonymous mutations at the Voltage-Gated Sodium Channel locus, with genomic positions and haplotype names listed in Figure S14. H001 is the susceptible haplotype. The size of each pie indicates frequency in the dataset with color corresponding to region according to legend.

**Figure S18:**
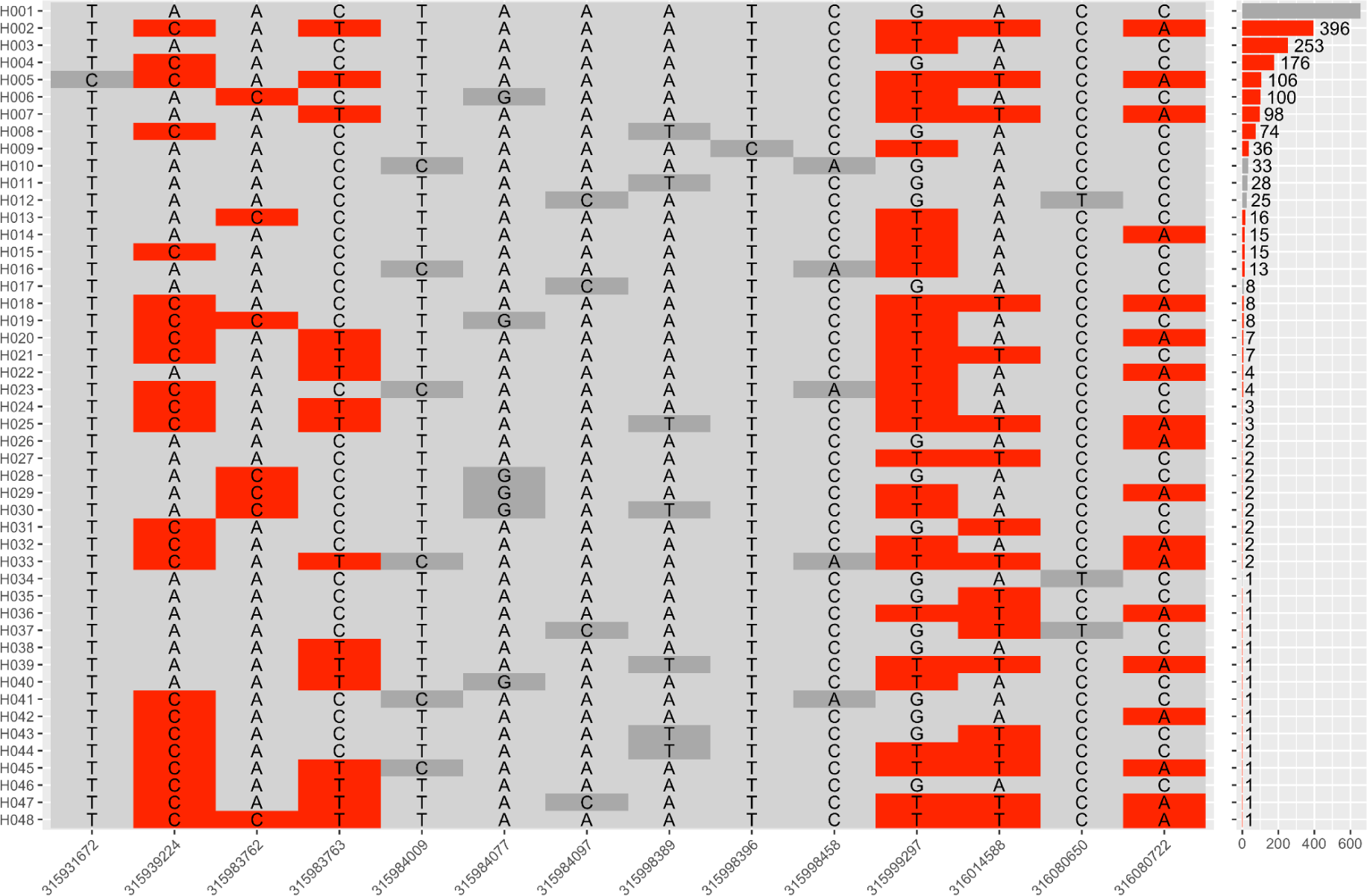
Knock-Down Resistance mutation haplotypes and frequencies. Each row corresponds to a haplotype with names labeled on the left and frequencies across the entire Aaeg1200 panel on the right. Light gray indicates major allele at each position. Dark gray indicates a non-synonymous mutation (only alleles with frequency > 0.01 included). Red indicates known resistance mutations: F1534C (315939224), V1016G (315983762), V1016I (315983763), I915K (315999297), S723T (316014588), V410L (316080722). The fully susceptible haplotype is labeled H001.

**Figure S19:**
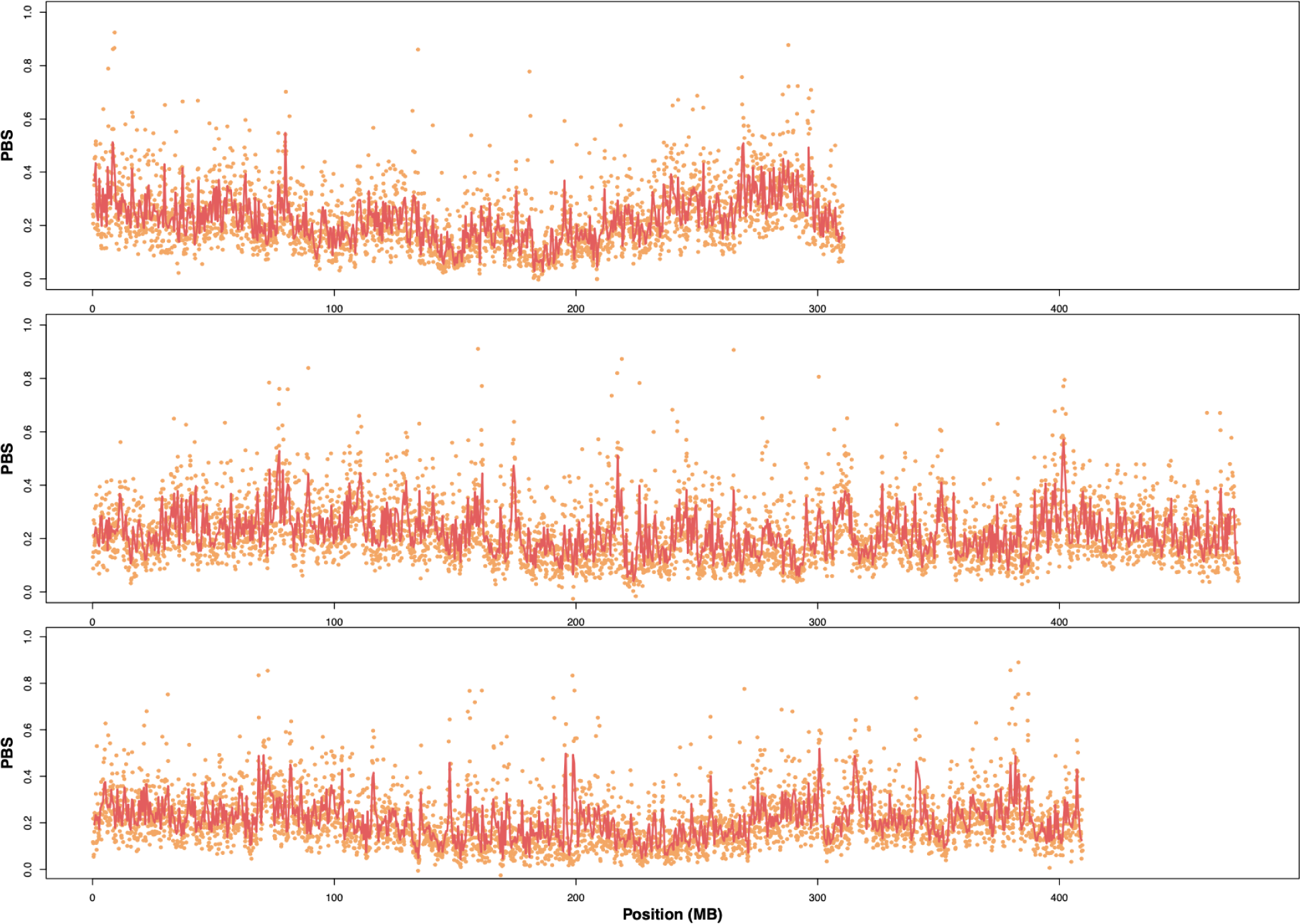
Sliding window analysis of genetic differentiation in North American populations. Population Branch Statistic (PBS) plotted as function of genomic position along each chromosome (chr1 on top, chr2 middle, chr3 on bottom) with North American populations as focal *Aaa* population compared to proto-*Aaa* populations with West African *Aaf* as the outgroup (see methods). Each dot shows PBS value for non-overlapping 100kb windows, and the red line shows PBS values for non-overlapping 500kb windows.

**Figure S20:**
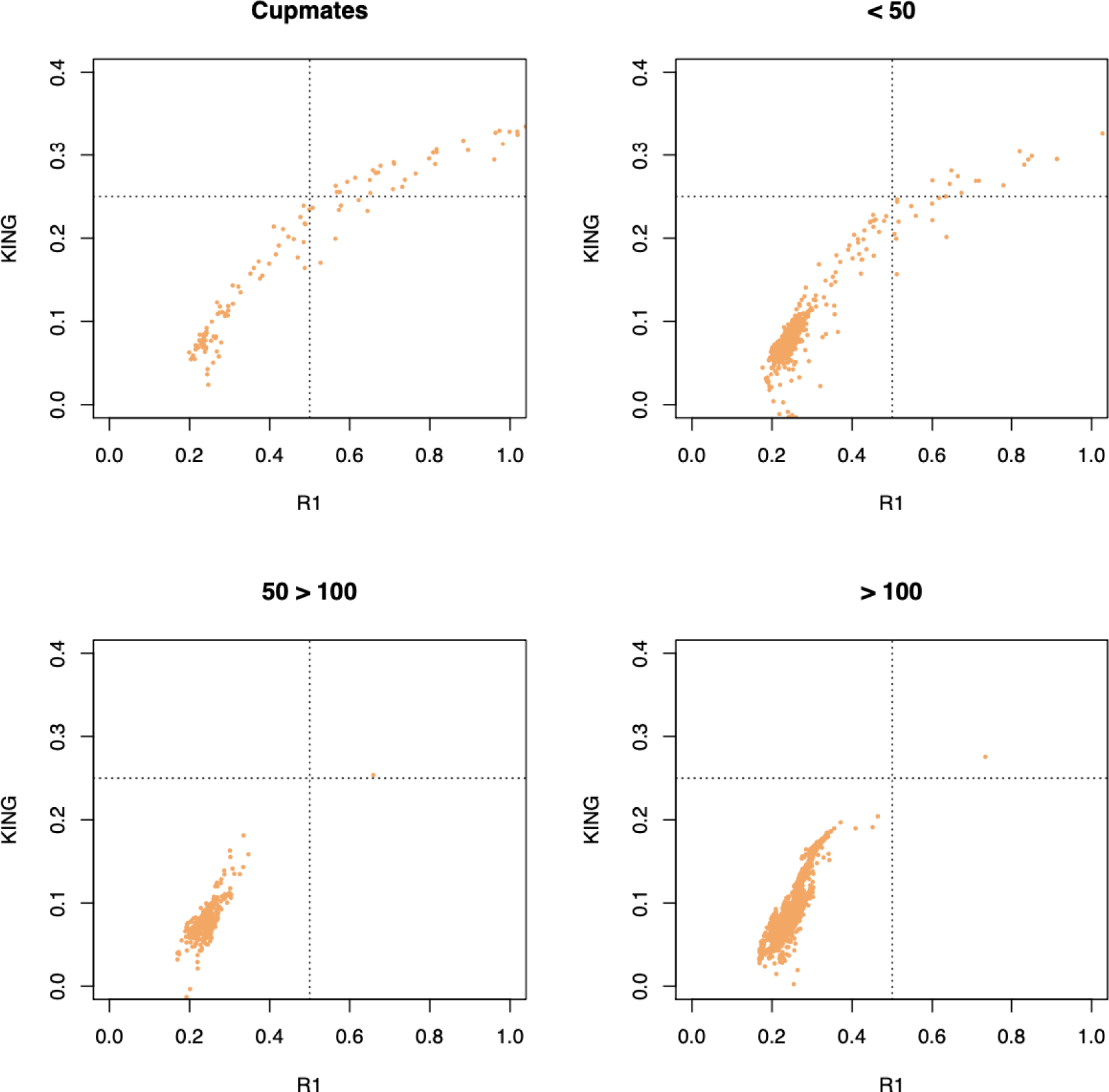
Relatedness analysis of individuals from ovitrap collections. Relatedness statistics (KING-robust and R1) calculated using NGSrelate shown for individual pairs according to physical proximity of collection locations. Cupmates indicates the two individuals were collected in the same ovicup. The remaining panels correspond to collections made from ovicups that were located <50 meters, between 50 and 100 meters, or more than 100 meters apart from each other. The dotted lines show the standard thresholds for first degree relatives so parent-offspring pairs and siblings fall in the upper right quadrant (Waples et al. 2019)

**Figure S21:**
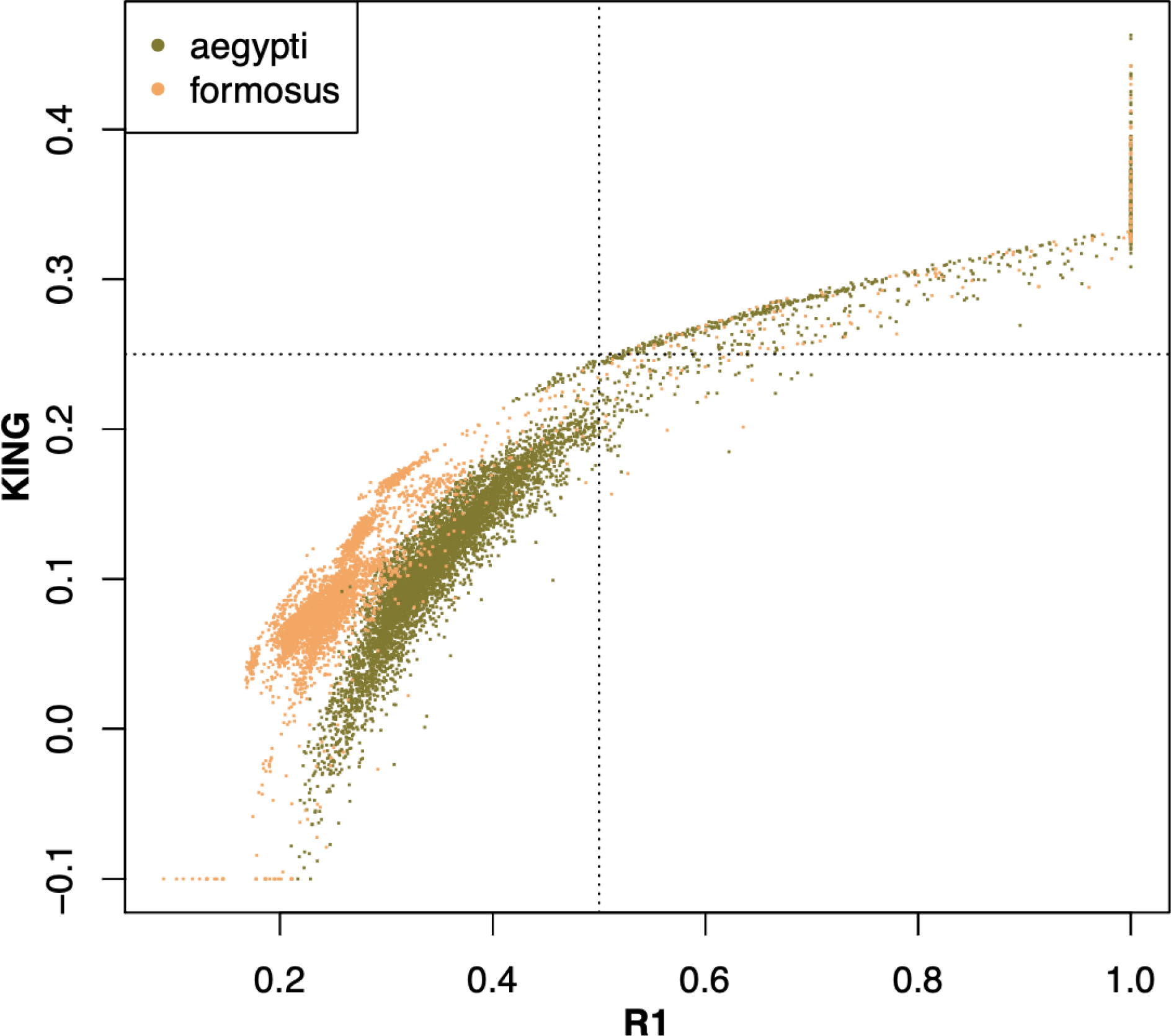
Within population relatedness analysis for all individuals. Relatedness statistics (KING-robust and R1) calculated using NGSrelate shown for individual pairs and colored according to subspecies. The dotted lines show the standard thresholds for first degree relatives so parent-offspring pairs and siblings fall in the upper right quadrant (Waples et al. 2019)

**Figure S22:**
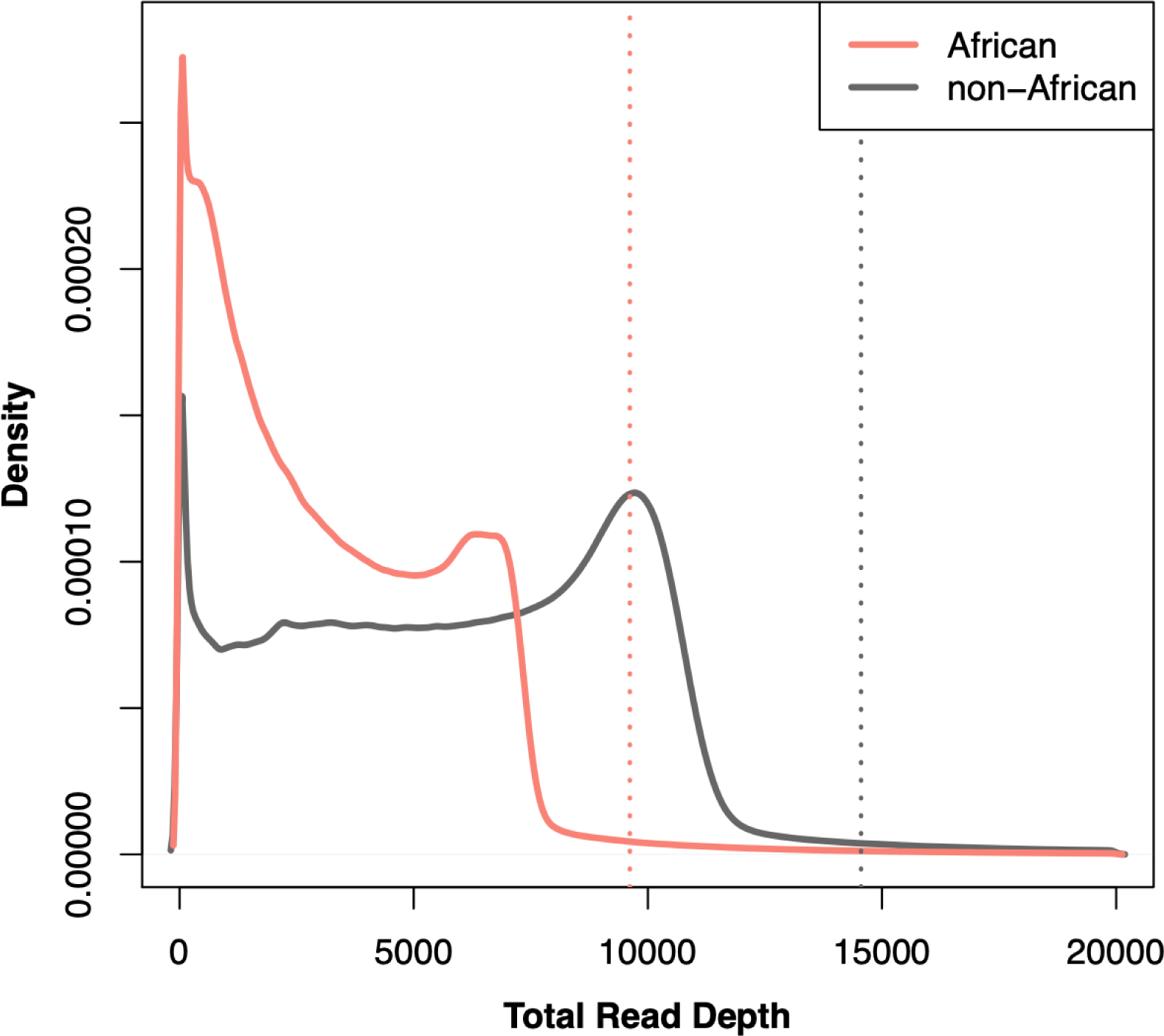
Total read depth across all samples for each subspecies. Total read depth per site calculated across all African (*Aaf*) and non-African (*Aaa*) individuals. Total read depth was calculated using ANGSD at all sites without robust site filters to find sites with outlier read depths. Dotted lines show the maximum total read depth for each set according to color.

**Figure S23:**
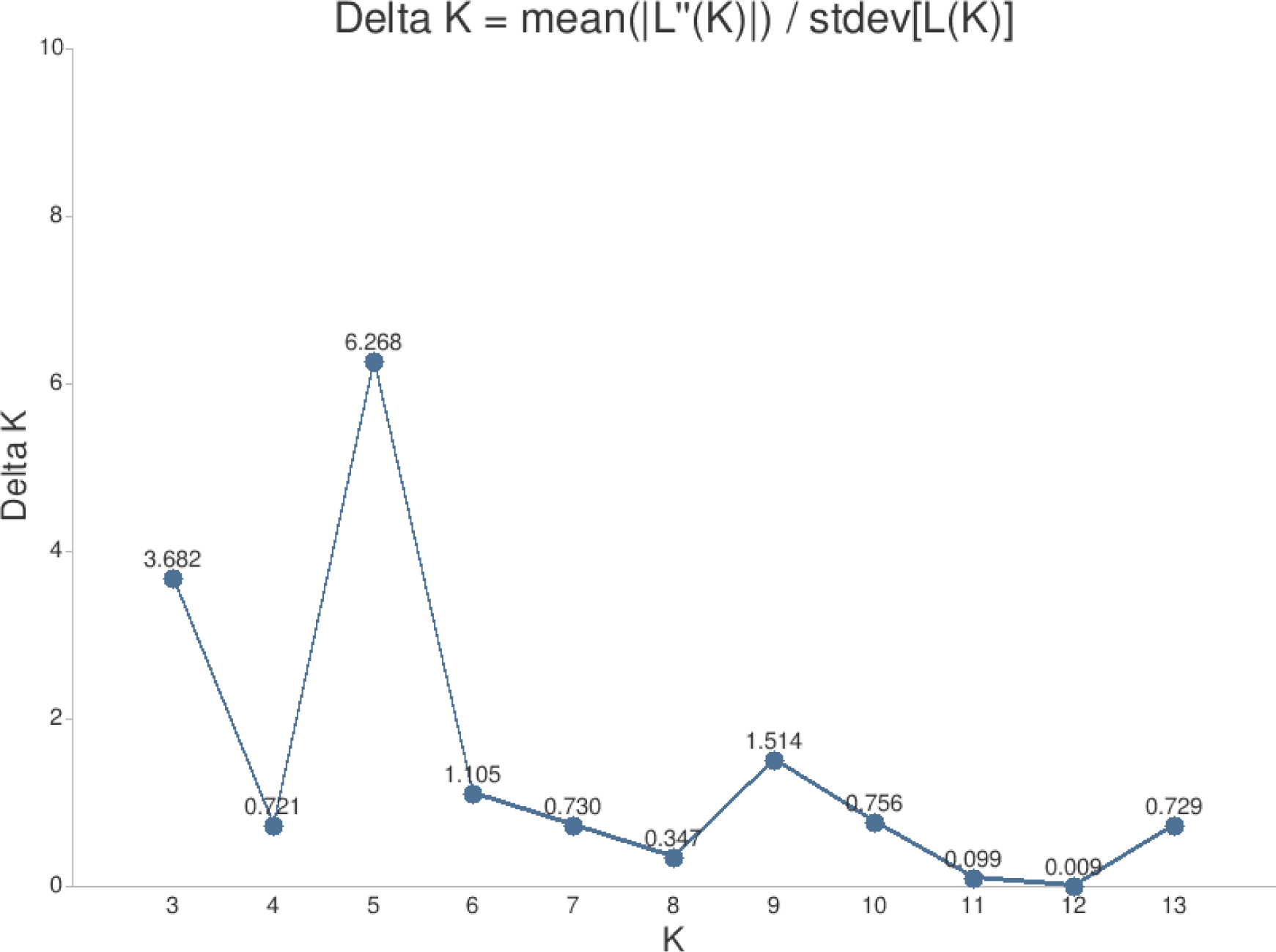
Evanno method to find best K for admixture analysis. Output from 20 replicate runs of admixture analysis with NGSadmix were submitted to CLUMPAK (tau.evolseq.net/clumpa) for evaluation.

**Figure S24:**
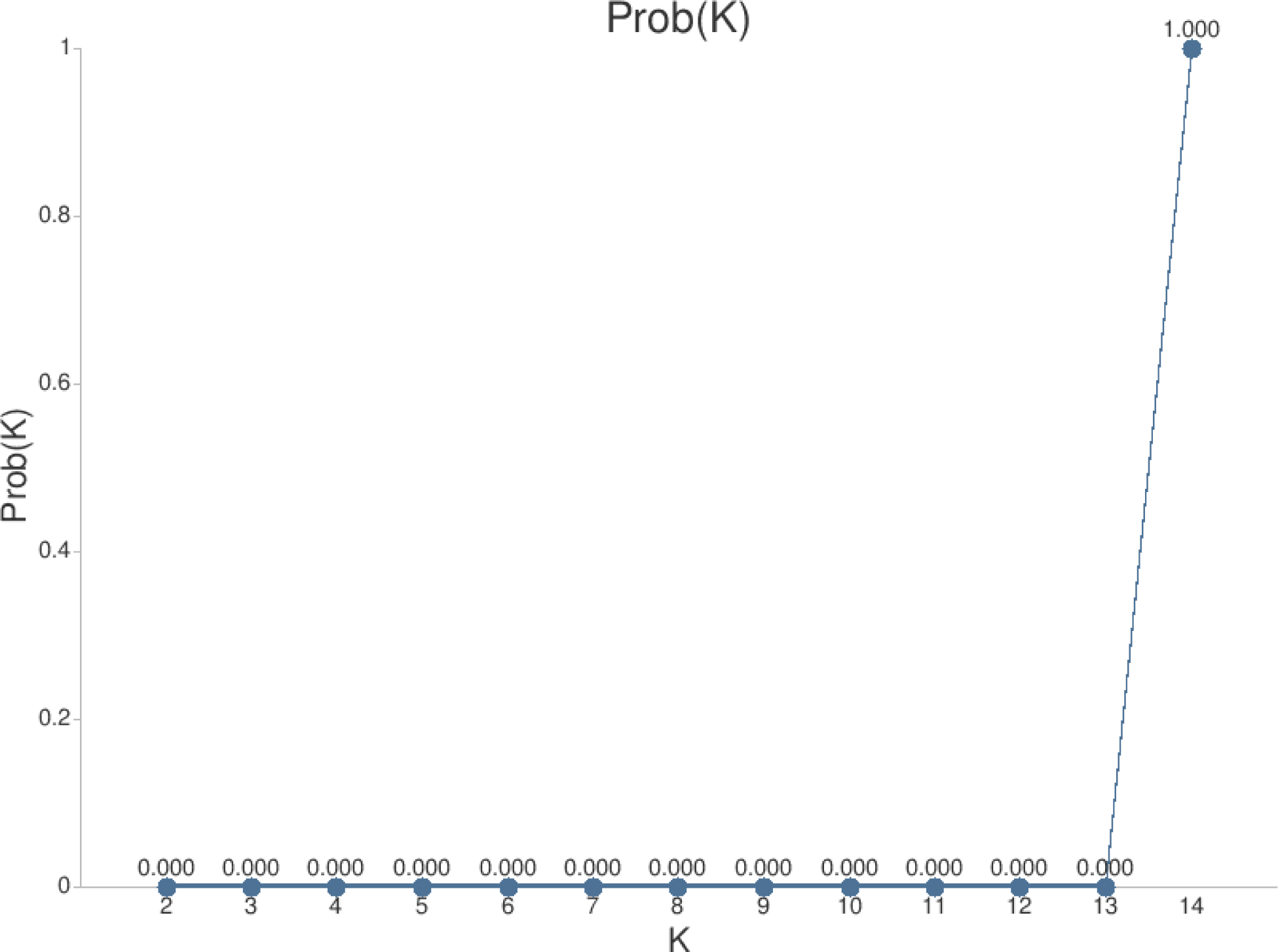
Evaluation of admixture analysis probabilities with CLUMPAK. Output from 20 replicate runs of admixture analysis with NGSadmix were submitted to CLUMPAK (tau.evolseq.net/clumpa) for evaluation. Using median values of Ln(Pr Data) the k for which Pr(K=k) is highest: 14

**Figure S25:**
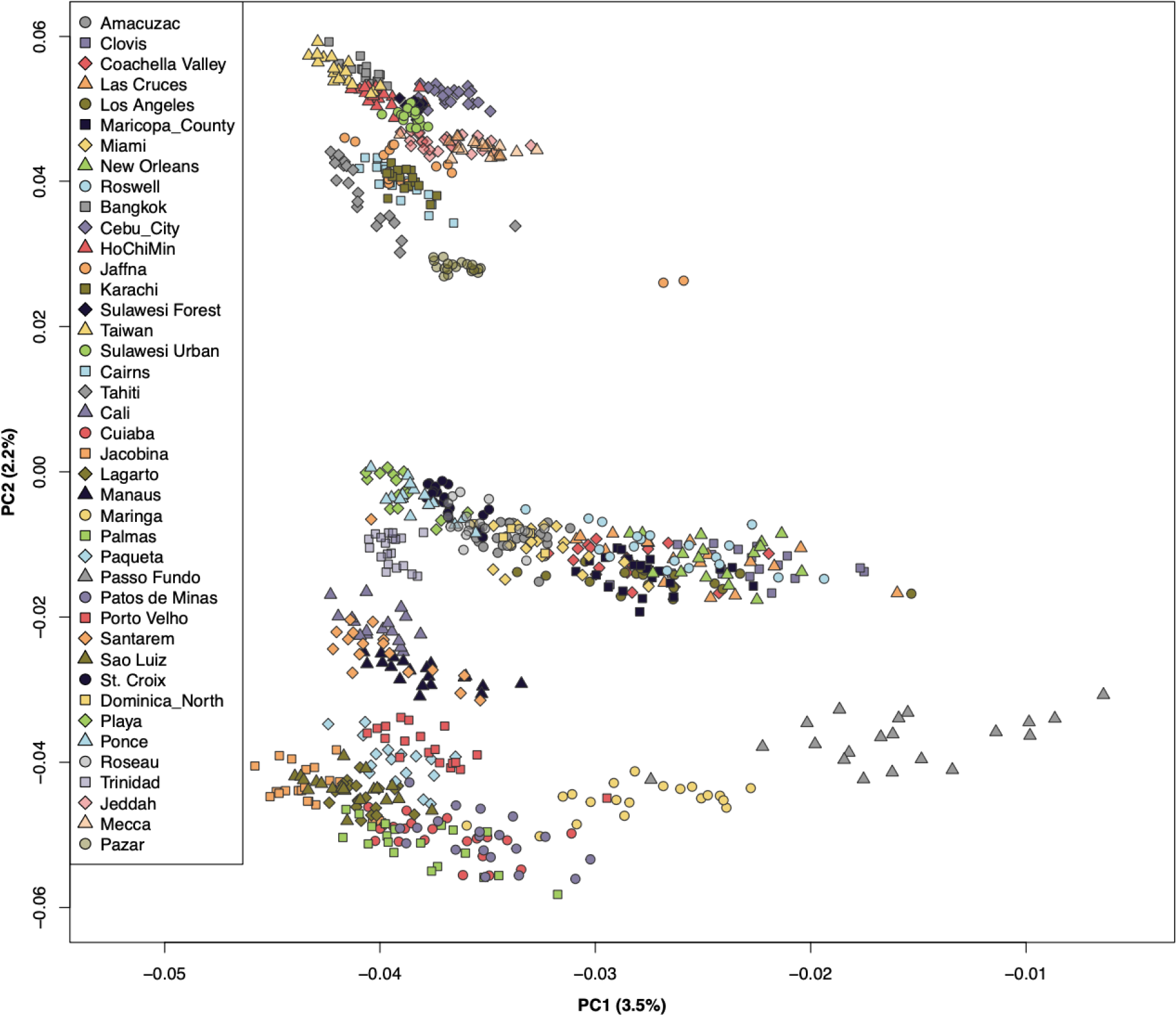
Principal components analysis of *Aaa* populations. The first two principal components shown on x and y axes with percent of genetic variation explained shown in parentheses in axis labels. Each dot shows position for an individual with population identity shown by color and point shape according to legend.

**Figure S26:**
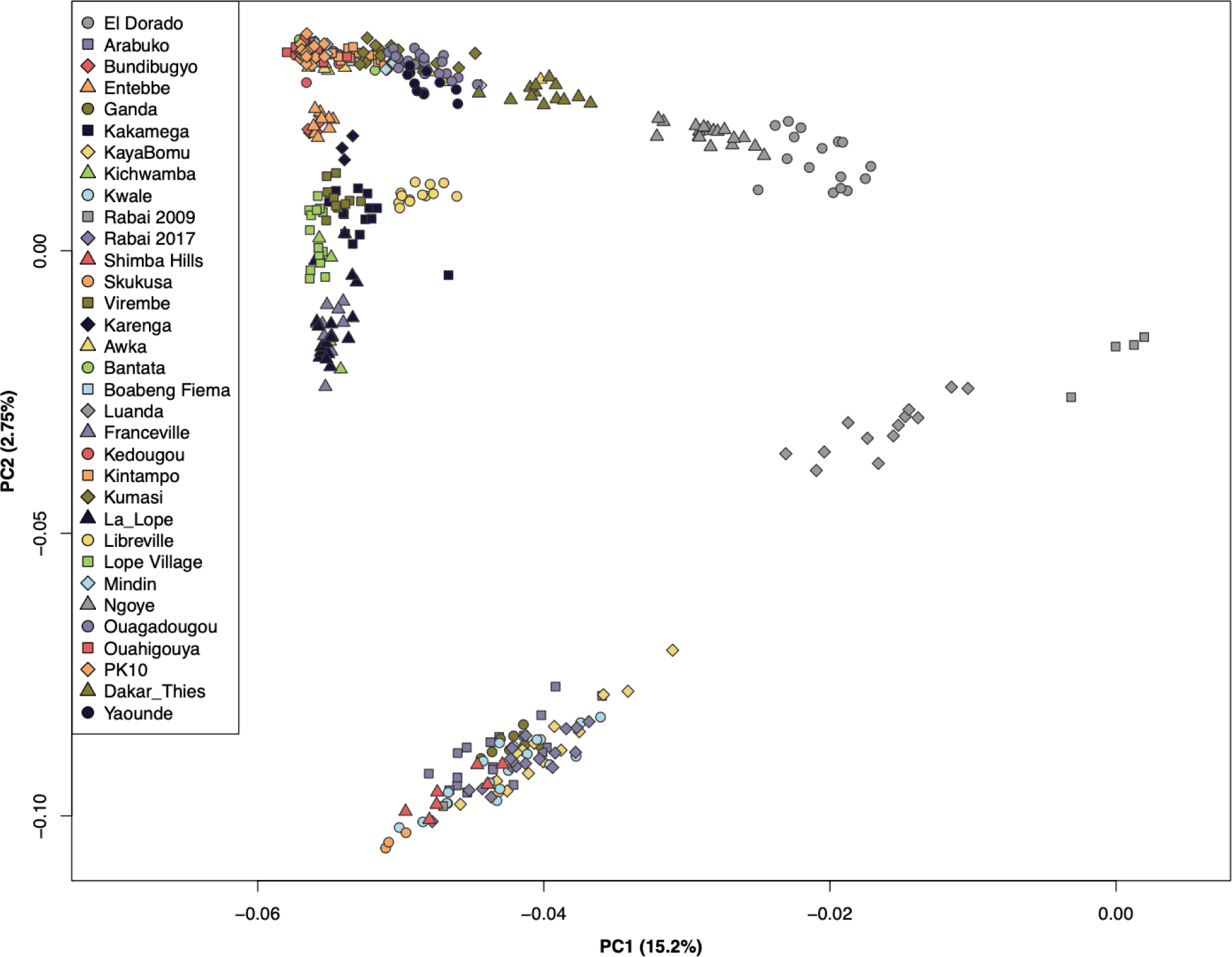
Principle components analysis of *Aaf* and proto-*Aaa* populations. The first two principal components shown on x and y axes with percent of genetic variation explained shown in parentheses in axis labels. Each dot shows position for an individual with population identity shown by color and point shape according to legend.

**Figure S27:**
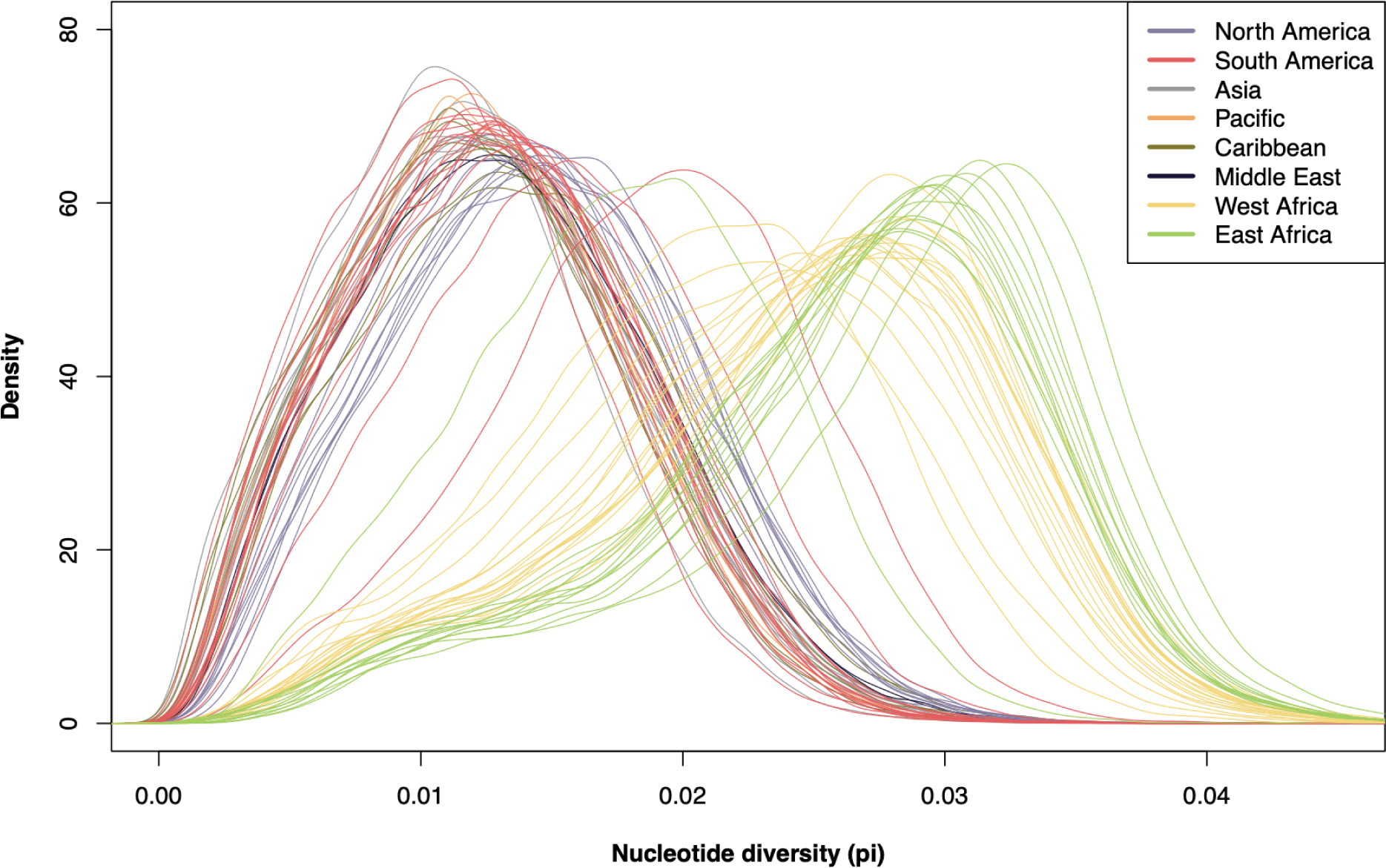
Distribution of nucleotide diversity. The nucleotide diversity *π* was calculated using ANGSD in one megabase windows along all three chromosomes for each population separately. The distribution was summarized using the density function in R and colors correspond to region of origin as indicated in the legend.

**Figure S28:**
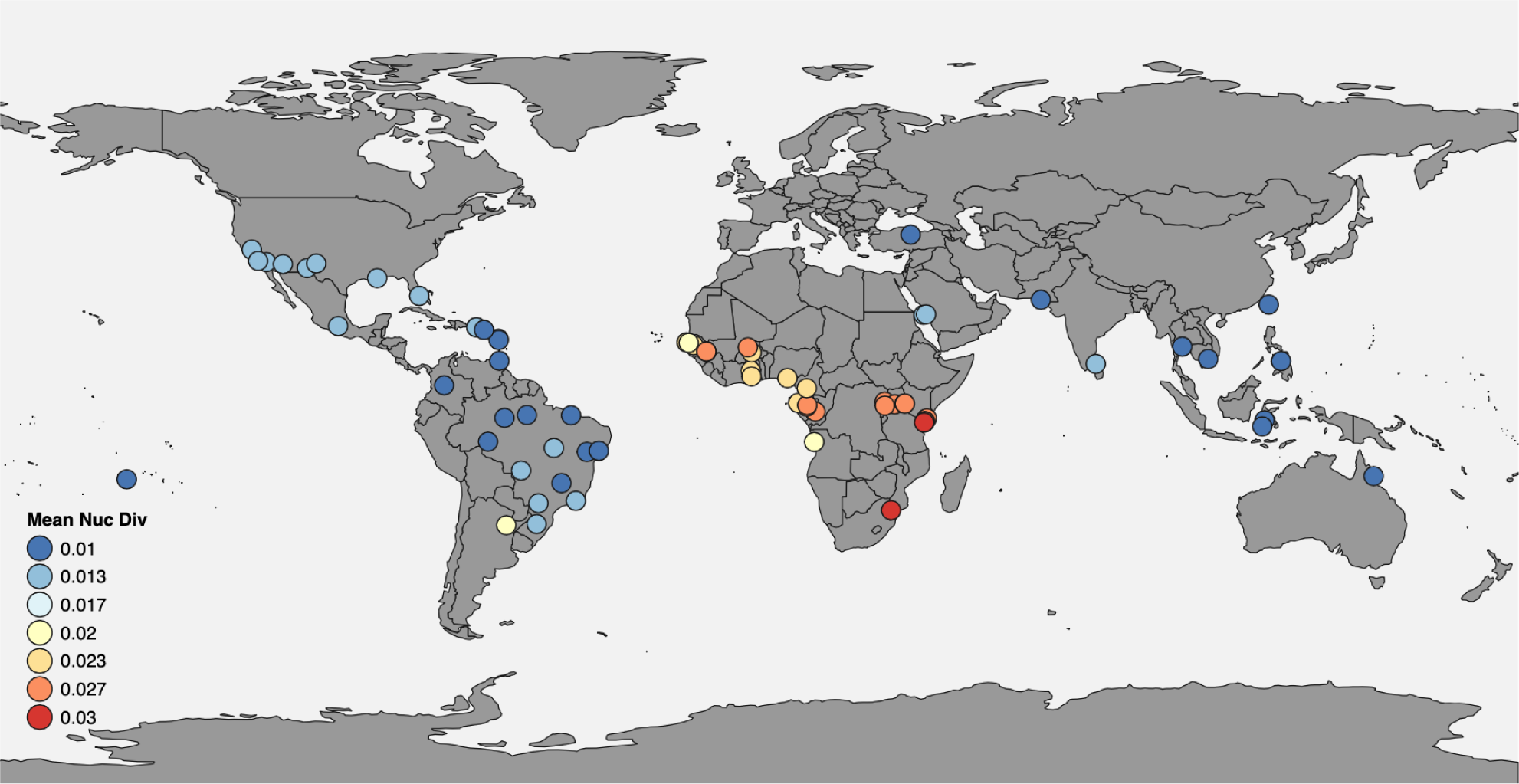
Mean nucleotide diversity varies globally. Mean nucleotide diversity (*π*) was calculated from 1 MB windows across all chromosomes, rounded into bins as indicated by the legend and plotted according to sampling locations.

**Figure S29:**
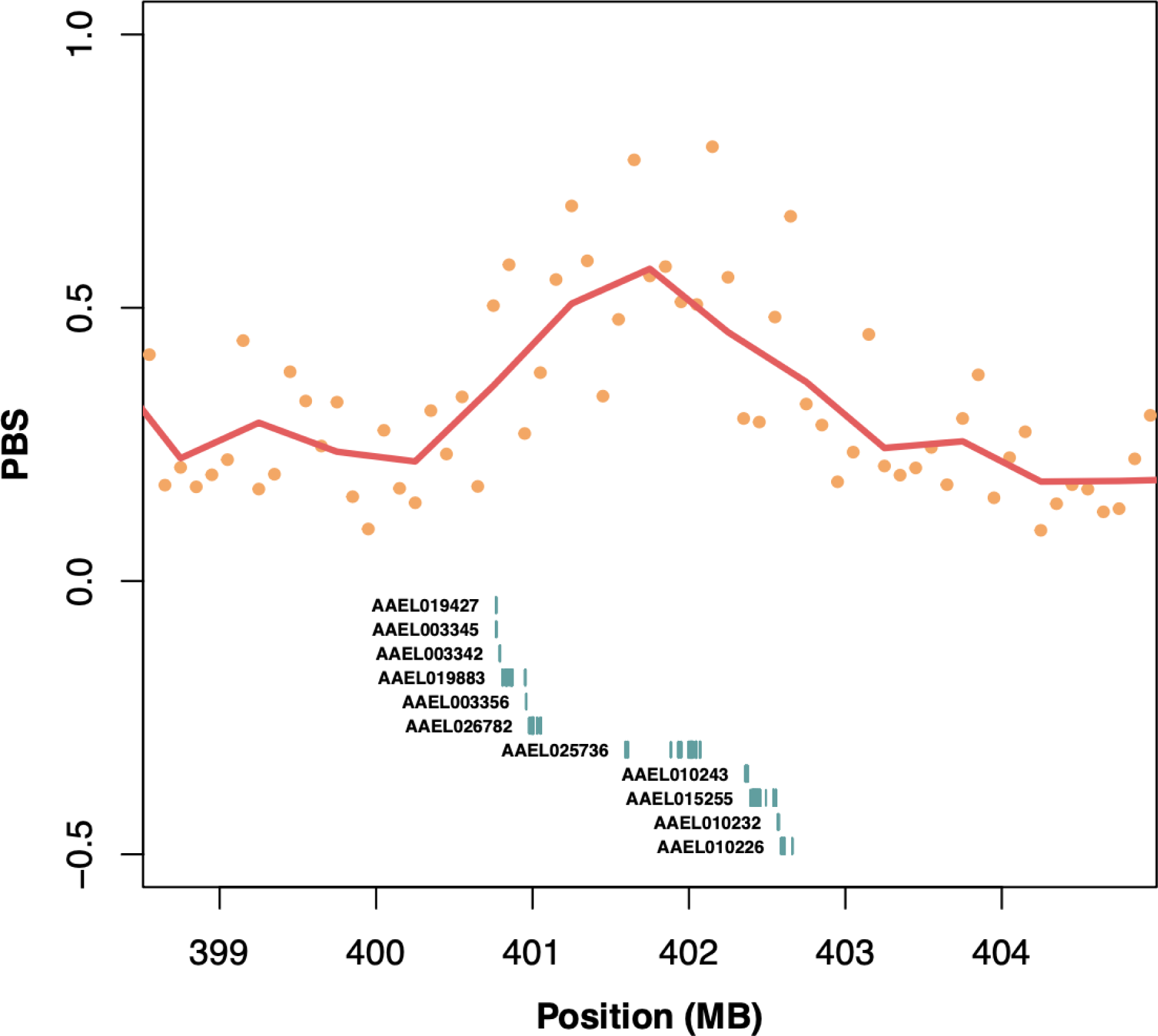
The strongest window of genetic differentiation in North American populations. Population Branch Statistic (PBS) plotted as a function of genomic position on chromosome chr1 at second highest peak with North American populations as focal *Aaa* population compared to proto-*Aaa* populations with West African *Aaf* as the outgroup (see methods). Each dot shows PBS value for non-overlapping 100kb windows, and the red line shows PBS values for non-overlapping 500kb windows. Exons from VectorBase-66_AaegyptiLVP_AGWG.gff shown below.

**Figure S30:**
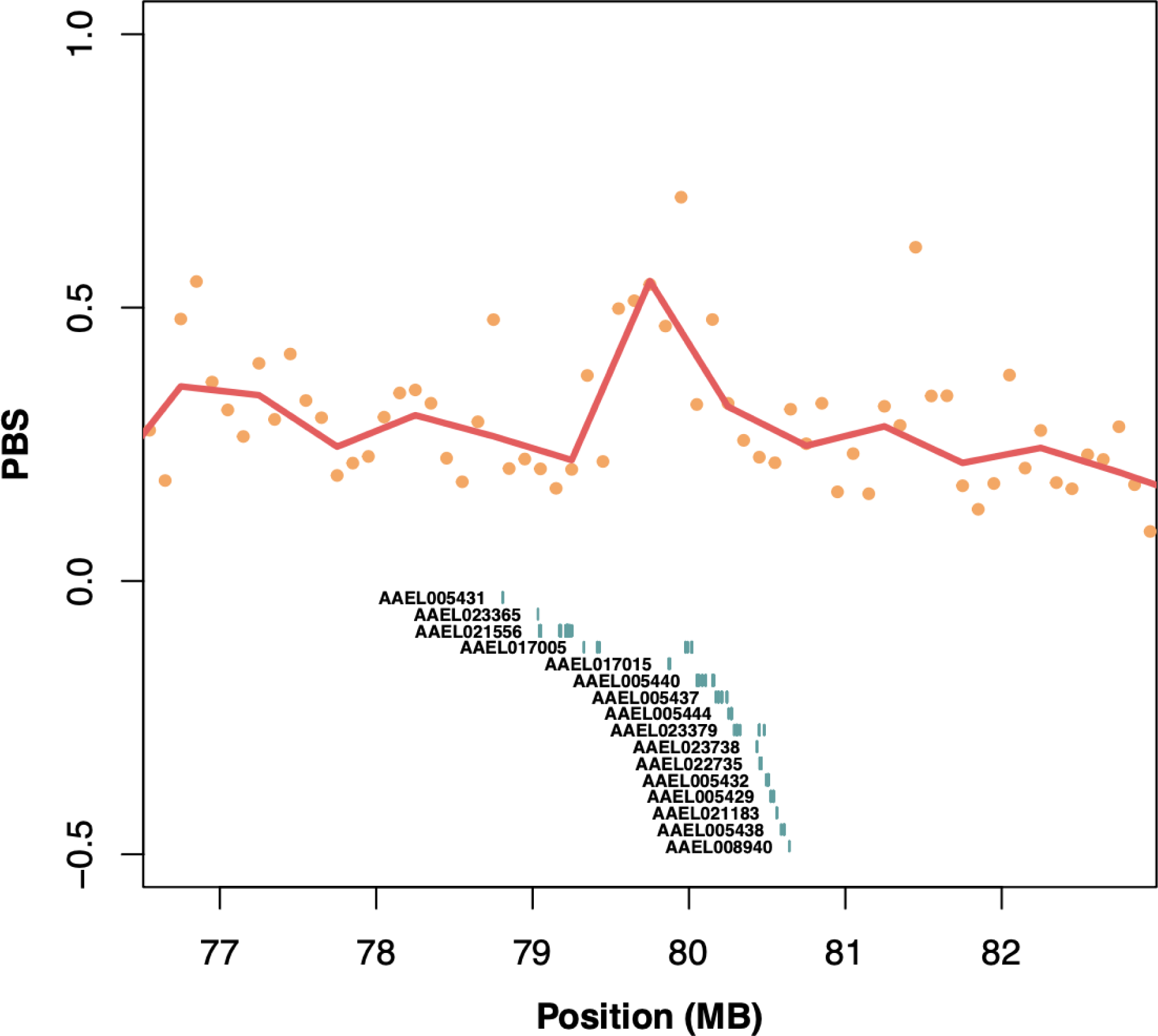
Second strongest window of genetic differentiation in North American populations. Population Branch Statistic (PBS) plotted as a function of genomic position on chromosome chr1 at second highest peak with North American populations as focal *Aaa* population compared to proto-*Aaa* populations with West African *Aaf* as the outgroup (see methods). Each dot shows PBS value for non-overlapping 100kb windows, and the red line shows PBS values for non-overlapping 500kb windows. Exons from VectorBase-66_AaegyptiLVP_AGWG.gff shown below.

**Figure S31:**
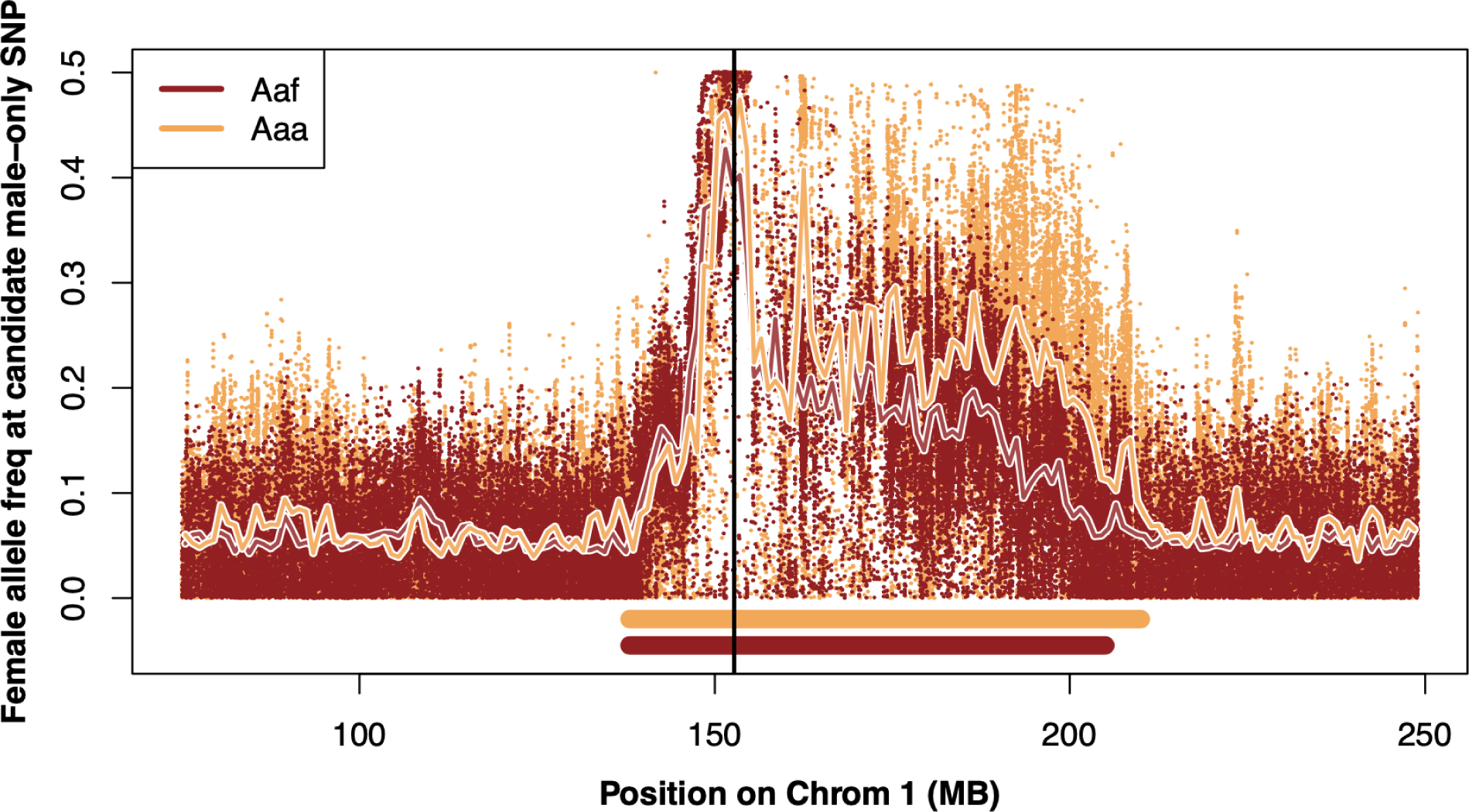
Male-female differentiation in *Aaa* and *Aaf*. Allele frequency differences between males and females at variable sites at intermediate frequencies (0.5) in males. Dots show frequencies for individual SNPs and lines show smoothed over one megabase windows. Red shows frequency for *Aaf* and Gold shows frequencies for *Aaa*. Vertical black line shows position of the male determining gene *nix*. Colored bars below indicate regions of excess differentiation with colors according to legend.

**Figure S32:**
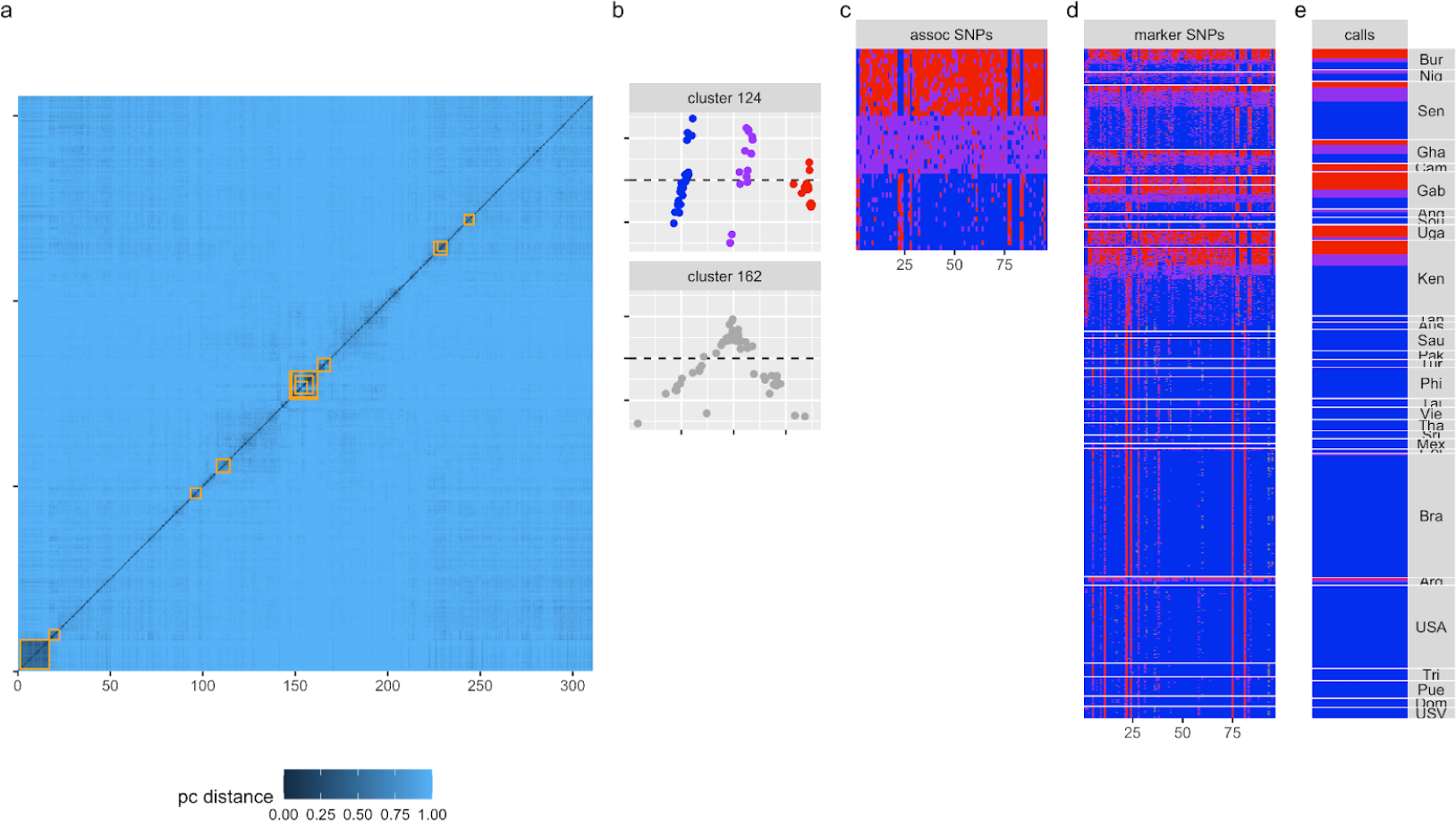
Putative inversion discovery pipeline. (a) a distance matrix is generated using Lostruct in 0.5mb blocks within all large country sample sets (N>25); regions of shared demographics (orange) are identified using pvclust; (b) putative inversions are tested via PCA and Kmeans for clustering into 3 distinct clusters with one admixed central cluster; (c) for validated inversions within their source country SNPs are associated (chi-square, P<=1e-7) with clusters 1 and 3; (d) marker SNPs are genotyped in all samples; (e) inversions calls are made using the dominant allele call for all marker SNPs

**Figure S33:**
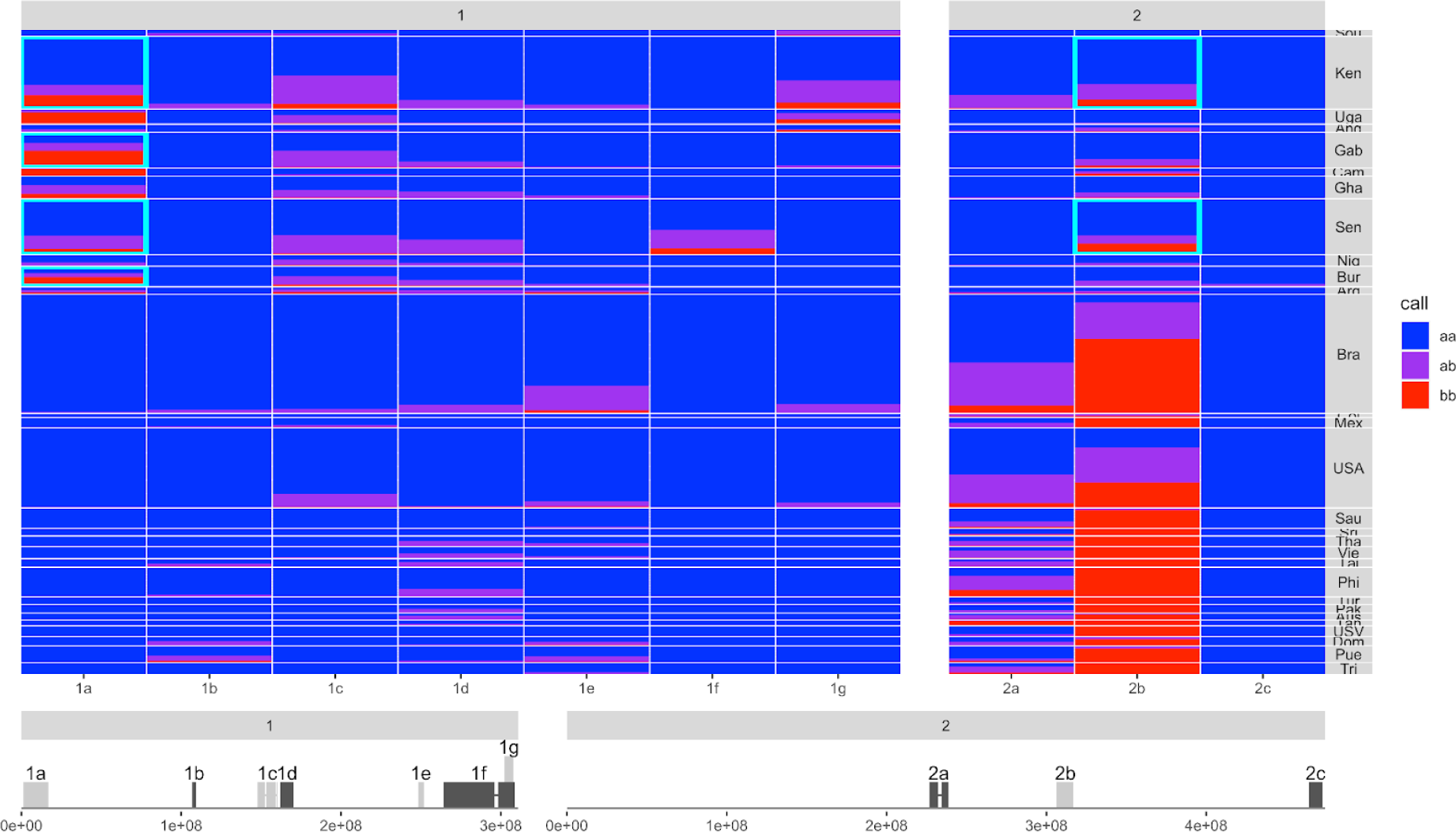
Frequency and location of candidate chromosomal inversions. Inverted regions cover a total of 10% of the *Ae. aegypti* L5 genome with frequencies ranging from less than 1 to 57% globally. Ordering of samples by continent and country here shows clear differences in frequency between *Aaa* and *Aaf* samples, though no inversion that is found at above 5% frequency is private to either subspecies. Deviations from Hardy Weinberg equilibrium in several African populations (cyan blocks) result from a deficit of heterokaryotes and may indicate selection acting on these karyotypes in these locations.

**Table S1.**
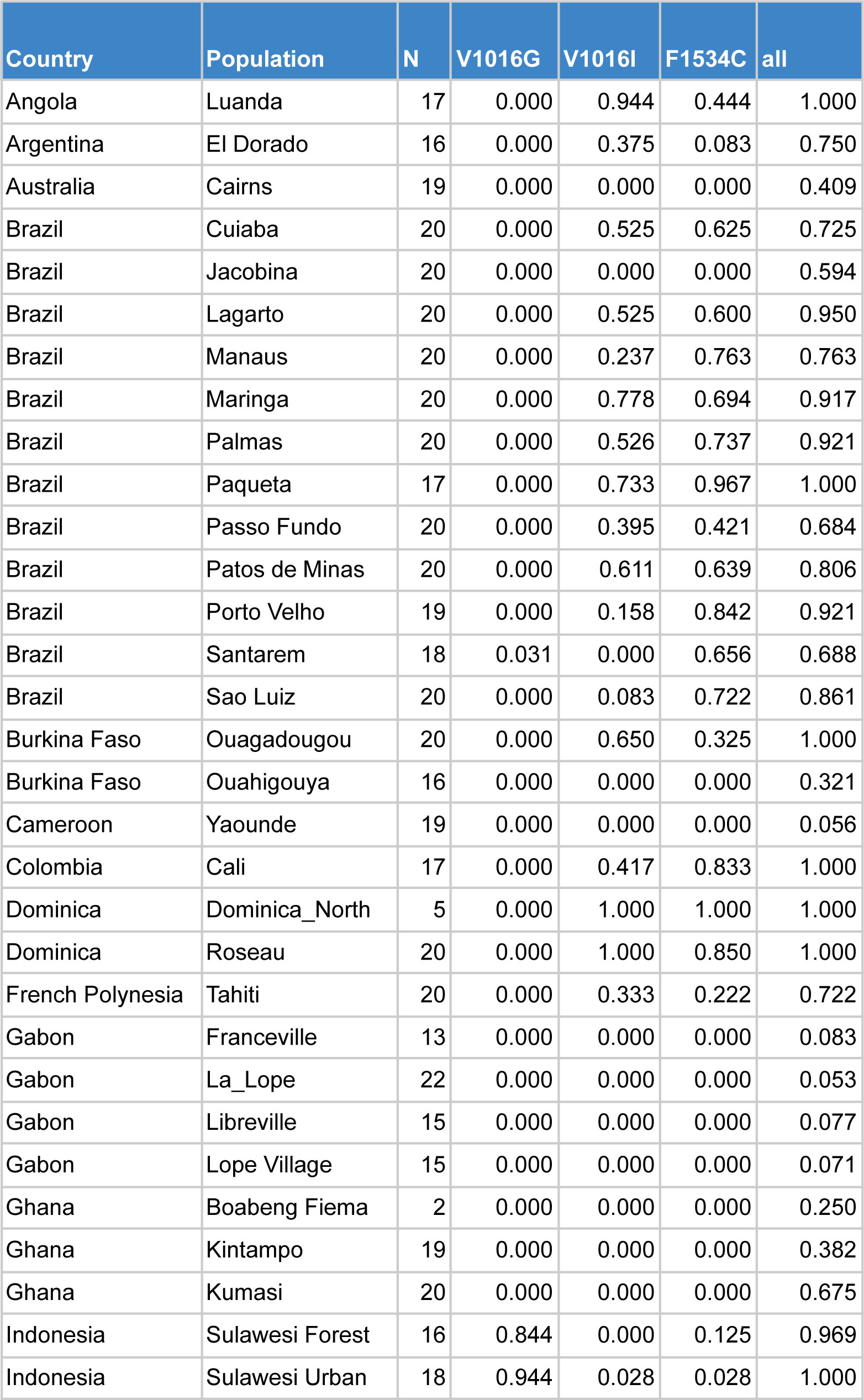
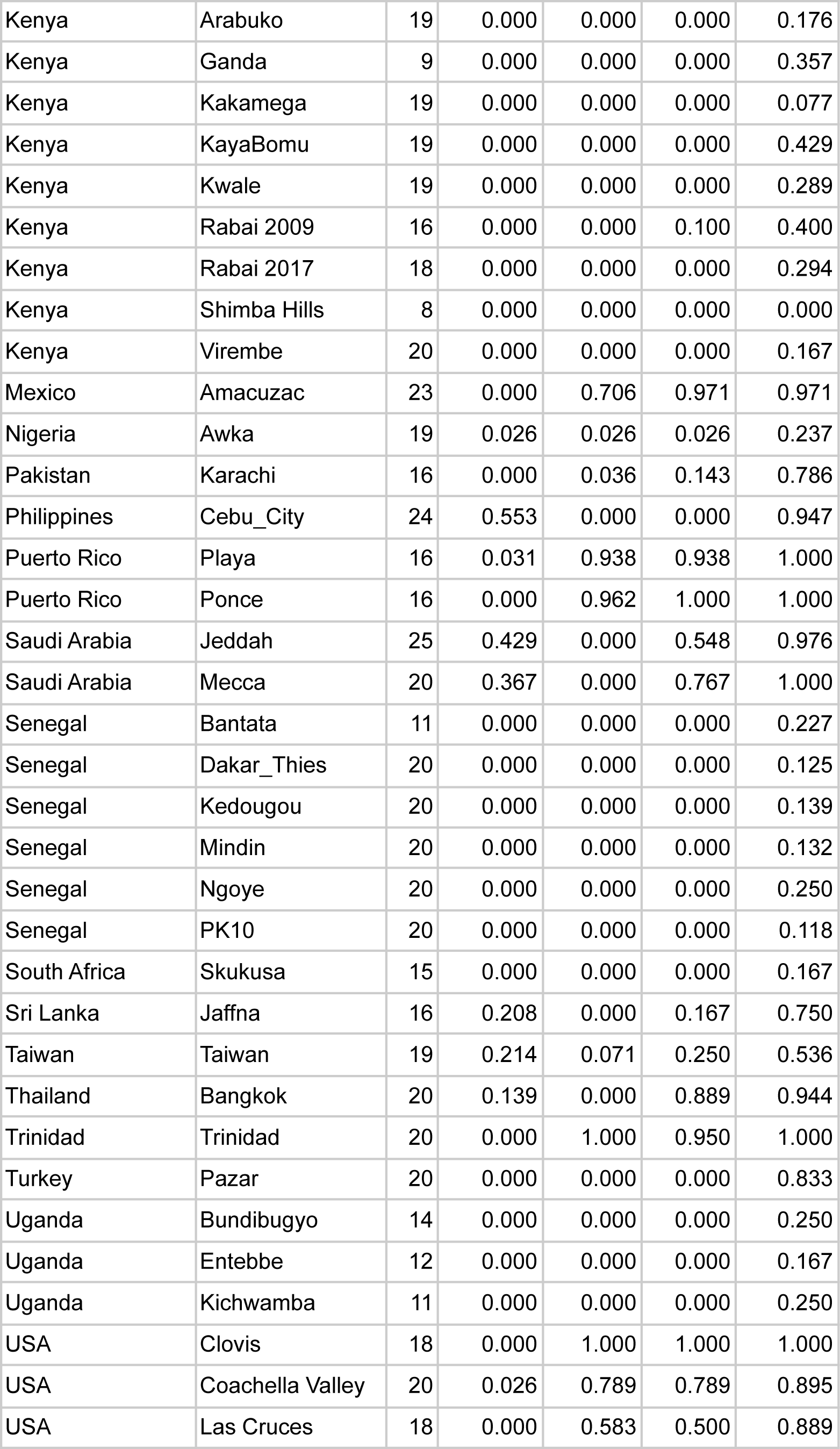
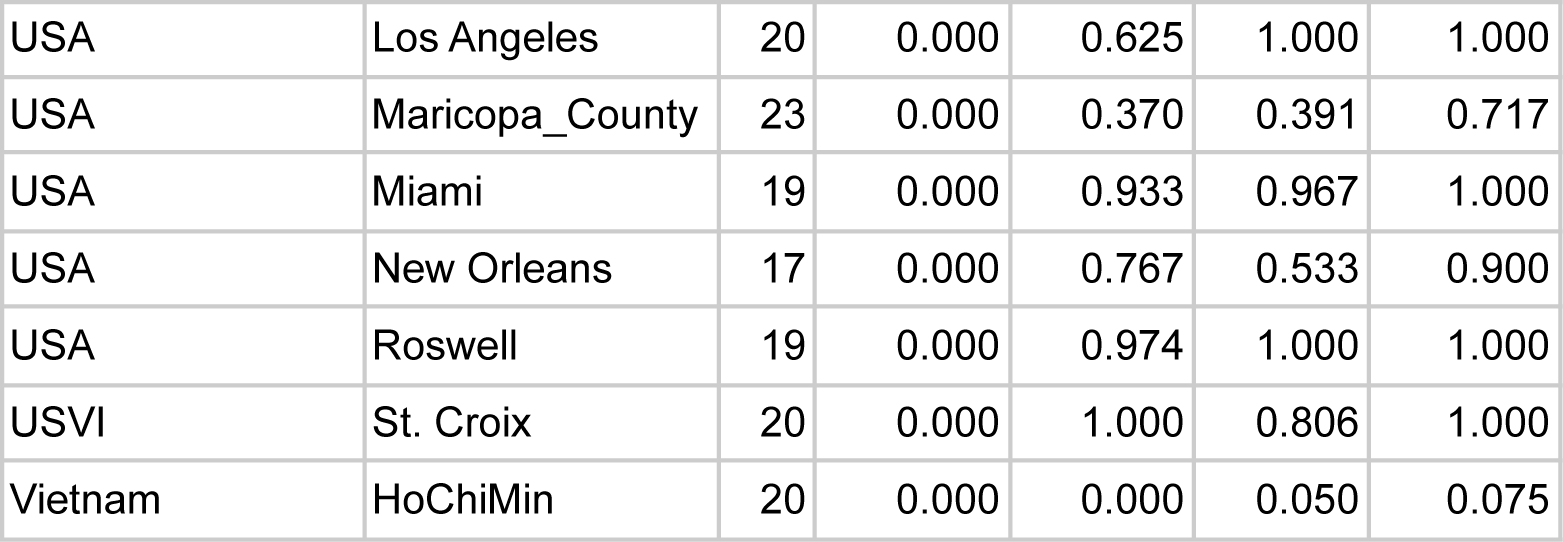
VGSC allele frequencies by population.

**Table S2: PBS Scan for Natural Selection.**

Included as a separate Excel Spreadsheet.

**Table S3:**
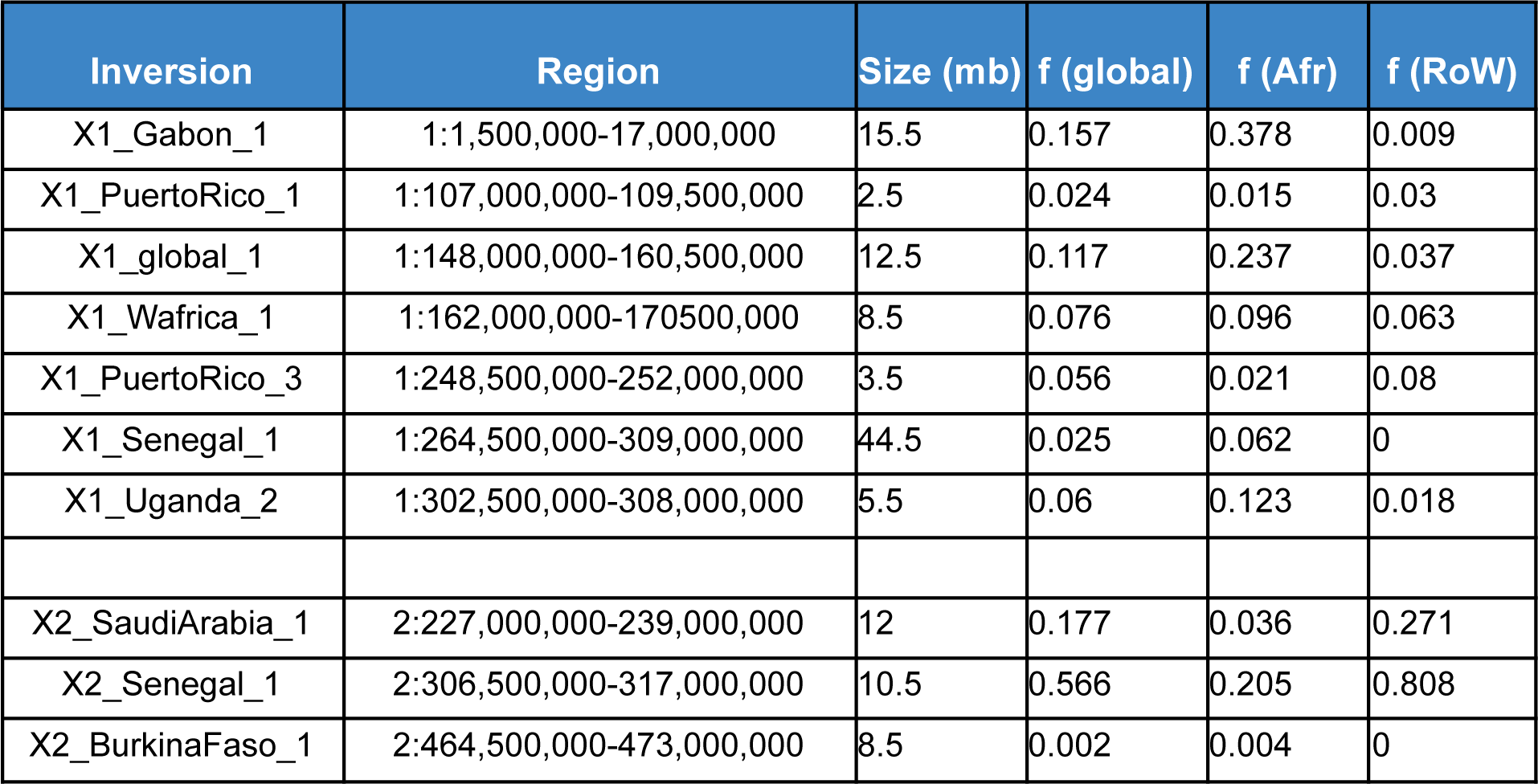
Candidate chromosomal inversions.

